# Expression changes confirm genomic variants predicted to result in allele-specific, alternative mRNA splicing

**DOI:** 10.1101/549089

**Authors:** Eliseos J. Mucaki, Ben C. Shirley, Peter K. Rogan

**Author notes:** Correspondence: Peter K. Rogan, Ph.D. SDRI 201A, Schulich School of Medicine and Dentistry The University of Western Ontario London ON Canada N6A 5C1 E.

## Abstract

Splice isoform structure and abundance can be affected by either non-coding or masquerading coding variants that alter the structure or abundance of transcripts. When these variants are common in the population, these non-constitutive transcripts are sufficiently frequent so as to resemble naturally occurring, alternative mRNA splicing. Prediction of the effects of such variants has been shown to be accurate using information theory-based methods. Single nucleotide polymorphisms (SNPs) predicted to significantly alter natural and/or cryptic splice site strength were shown to affect gene expression. Splicing changes for known SNP genotypes were confirmed in HapMap lymphoblastoid cell lines with gene expression microarrays and custom designed q-RT-PCR or TaqMan assays. The majority of these SNPs (15 of 22) as well as an independent set of 24 variants were then subjected to RNAseq analysis using the ValidSpliceMut web beacon (http://validsplicemut.cytognomix.com), which is based on data from the Cancer Genome Atlas and International Cancer Genome Consortium. SNPs from different genes analyzed with gene expression microarray and q-RT-PCR exhibited significant changes in affected splice site use. Thirteen SNPs directly affected exon inclusion and 10 altered cryptic site use. Homozygous SNP genotypes resulting in stronger splice sites exhibited higher levels of processed mRNA than alleles associated with weaker sites. Four SNPs exhibited variable expression among individuals with the same genotypes, masking statistically significant expression differences between alleles. Genome-wide information theory and expression analyses (RNAseq) in tumour exomes and genomes confirmed splicing effects for 7 of the HapMap SNP and 14 SNPs identified from tumour genomes. q-RT-PCR resolved rare splice isoforms with read abundance too low for statistical significance in ValidSpliceMut. Nevertheless, the web-beacon provides evidence of unanticipated splicing outcomes, for example, intron retention due to compromised recognition of constitutive splice sites. Thus, ValidSpliceMut and q-RT-PCR represent complementary resources for identification of allele-specific, alternative splicing.

## Introduction

Accurate and comprehensive methods are needed for predicting impact of non-coding mutations, in particular, mRNA splicing defects, which are prevalent in genetic disease (Krawczak *et al*. 1992; Teraoka *et al*. 1999; Ars *et al*. 2003; Spielmann and Mundlos, 2016; Gloss and Dinger, 2018). This class of mutations may account for as much as 62% of point mutations (López-Bigas *et al*. 2005). Large transcriptome studies have suggested that a large fraction of genome-wide association studies (GWAS) signals for disease and complex traits are due to SNPs affecting mRNA splicing (Park *et al*. 2018). ValidSpliceMut (Shirley *et al*. 2019) presents evidence of altered splicing (Viner *et al*. 2014; Dorman *et al*. 2014) for 309,848 validated genome splice-variant predictions (Shirley *et al*. 2013). The majority of mutations were associated with exon skipping, cryptic site use, or intron retention, and in these cases ValidSpliceMut assigns a molecular phenotype classification to all variants as either aberrant, likely aberrant or inducing alternative isoforms.

While allele-specific alternative splicing can predispose for disease susceptibility (Park *et al*. 2018), these genetic variations also are associated with common phenotypic variability in populations (Hull *et al*. 2007). Soemedi *et al*. (2017) determined that 10% of a set of published disease-causing exonic mutations (N=4,964) altered splicing. Their analysis of a control set of exonic SNPs common among those without disease phenotypes revealed a smaller proportion (3%) that altered splicing (N=228). However, we recently showed that splice-altering, common SNPs are considerably more abundant in tumor genomes in the ValidSpliceMut web-beacon (http://validsplicemut.cytognomix.com; Shirley *et al*. 2019). Variants with higher germline population frequencies which impact splicing are less likely than rare mutations having direct splicing effects to be involved in Mendelian diseases or cancer. The present study analyzes predicted splice-altering polymorphic variants in genotyped lymphoblastoid cell lines by q-RT-PCR, expression microarrays of samples of known SNP genotypes, and high throughput expression data corresponding to sequenced tumour exomes and genomes. The relatively high frequencies of these variants enables comparisons of expressed transcripts in multiple individuals and genotypes. Effects of these SNPs are confirmed by multiple methods, although the supporting evidence from these distinct approaches is often complementary, rather than entirely concordant.

An estimated 90 to 95% of all multi-exon genes are alternatively spliced (Wang *et al*. 2008; Pan *et al*. 2008; Baralle *et al*. 2017). The selection of splicing signals involves exon and intron sequences, complementarity with snRNAs, RNA secondary structure and competition between spliceosomal recognition sites (Berget 1995; Moore and Sharp 1993; Park *et al*. 2018). U1 snRNP interacts with the donor (or 5’) splice site (Séraphin *et al*. 1988; Zhuang *et al*. 1986) and U2 (and U6) snRNP with the acceptor and branch sites of pre-mRNA (Parker *et al*. 1987; Wu and Manley, 1989). The majority of human splice donors (5’) and acceptors (3’) base pair with the U1 and U2 RNAs in spliceosomes, but are generally not precisely complementary to these sequences (Rogan *et al*. 2003). Additional exonic and intronic cis-regulatory elements can promote or suppress splice site recognition through recruitment of trans-acting splicing factors. SR proteins are positive trans-acting splicing factors which contain RNA-recognition motifs (RRM) and a carboxy-terminal domain enriched in Arg/Ser dipeptides (SR domain; Birney *et al*. 1993). Binding of RRMs in pre-mRNA enhances exon recognition by promoting interactions with spliceosomal and other proteins (Fu and Maniatis 1992). SR proteins function in splice site communication by forming an intron bridge needed for exon recognition (Zuo and Maniatis 1996). Factors that negatively impact splicing include heterogeneous nuclear ribonucleoproteins (hnRNPs; Martinez-Contreras *et al*. 2007).

Splicing mutations affect normal exon recognition by altering the strengths of natural donor or acceptor sites and proximate cryptic sites, either independently or simultaneously. Weakened splice sites reduce of kinetics of mRNA processing, leading to an overall decrease in full length transcripts, increased exon skipping, cryptic splice site activation within exons or within adjacent introns, intron retention, and inclusion of cryptic, pseudo-exons (Talerico and Berget 1990; Carothers *et al*. 1993; Buratti 2006; Park *et al*. 2018). The kinetics of splicing at weaker cryptic sites is also slower than at natural sites (Domenjoud *et al*. 1993). Mutations strengthen cryptic sites either by increasing resemblance to “consensus sequences” (Nelson and Green 1990) or by modulating the levels of SR proteins contributing to splice site recognition (Mayeda and Krainer 1992; Cáceres *et al*. 1994). Mutations affecting splicing regulatory elements (Dietz *et al*. 1993; Richard and Beckmann 1995) disrupt trans-acting SR protein interactions (Staknis and Reed 1994) with distinct exonic and intronic cis-regulatory elements (Black 2003).

Information theory-based (IT-based) models of donor and acceptor mRNA splice sites reveal the effects of changes in strengths of individual sites (termed *R_i_*; Rogan *et al*. 1998; Rogan *et al*. 2003). This facilitates prediction of phenotypic severity (Rogan and Schneider 1995; von Kodolitsch *et al*. 1999; von Kodolitsch *et al*. 2006). The effects of splicing mutations can be predicted *in silico* by information theory (Rogan and Schneider 1995; Rogan *et al*. 1998; Kannabiran *et al*. 1998; Svojanovsky *et al*. 2000; Rogan *et al*. 2003; Viner *et al*. 2014; Dorman *et al*. 2014; Caminsky *et al*. 2014; Caminsky *et al*. 2016; Mucaki *et al*. 2016; Shirley *et al*. 2019), and these predictions can be confirmed by *in vitro* experimental studies (Vockley *et al*. 2000; Rogan *et al*. 2003; Lamba *et al*. 2003; Khan *et al*. 2004; Susani *et al*. 2004; Hobson *et al*. 2006; Caux-Moncoutier *et al*. 2009; Olsen *et al*. 2014; Vemula *et al*. 2014; Peterlongo *et al*. 2015). Strengths of one or more splice sites may be altered and, in some instances, concomitant with amino acid changes in coding sequences (Rogan *et al*. 1998; Peterlongo *et al*. 2015). Information analysis has been a successful approach for recognizing non-deleterious, sometimes polymorphic variants (Rogan and Schneider 1995; Colombo *et al*. 2013), and for distinguishing of milder from severe mutations (Rogan *et al*. 1998; von Kodolitsch *et al*. 1999; Lacroix *et al*. 2012).

Predicting the relative abundance of various transcripts by information analysis requires integration of the contributions of all pertinent cis-acting regulatory elements. We have applied quantitative methods to prioritize inferences as to which SNPs impact gene expression levels and transcript structure. Effects of mutations on combinations of splicing signals reveal changes in isoform structure and abundance (Mucaki *et al*. 2013; Caminsky *et al*. 2014). Multi-site information theory-based models have also been used to detect and analyze SNP effects on cis-acting promoter modules that contribute to establishing transcript levels (Vyhlidal *et al*. 2004; Bi and Rogan, 2004; Lu *et al*. 2017; Lu and Rogan 2019).

The robustness of this approach for predicting rare, deleterious splicing mutations justifies efforts to identify common SNPs that impact mRNA splicing. We previously described single nucleotide polymorphisms from dbSNP that affect splicing (Rogan *et al*. 1998; Nalla and Rogan 2005). Here, we explicitly predict and validate SNPs that influence mRNA structure and levels of expression of the genes containing them in immortalized lymphoblastoid cell lines and tumours. Since constitutive splicing mutations can arise at other locations within pre-mRNA sequences that elicit cryptic splicing, we examined whether more common genomic polymorphisms might frequently affect the abundance and structure of splice isoforms.

## Methods

### Information Analysis

The protein-nucleic acid interactions intrinsic to splicing can be analyzed using information theory, which comprehensively and quantitatively models functional sequence variation based on a thermodynamic framework (Schneider 1997). Donor and acceptor splice site strength can be predicted by the use of IT-based weight matrices derived from known functional sites (Rogan *et al*. 2003). The Automated Splice Site and Exon Definition server (ASSEDA) is an online resource based on the hg19 coordinate system to determine splice site information changes associated with genetic diseases (Mucaki *et al*. 2013). ASSEDA is now part of the MutationForecaster (http://www.mutationforecaster.com) variant interpretation system.

### Creation of Exon Array Database

Exon-level microarrays have been used to compare abnormal expression for different cellular states, which can then be confirmed by q-RT-PCR (Thorsen *et al*. 2008). We hypothesized that the predicted effect of SNPs on expression of the proximate exon would correspond to the expression of exon microarray probes of genotyped individuals in the HapMap cohort. We used the dose-dependent expression of the minor allele to qualify SNPs for subsequent information analysis consistent with alterations of mRNA splicing. Additional SNPs predicted by information analysis were also tested for effects on splicing (Nalla and Rogan, 2005).

Expression data were normalized using the PLIER (Probe Logarithmic Intensity Error) method on Affymetrix Human Exon 1.0 ST microarray data for 176 genotyped HapMap cell lines (Huang *et al*. 2007, Gene Expression Omnibus accession no. GSE 7792; Nembaware *et al*. 2008). Microarray probes which overlap SNPs, that were subsequently removed, were identified by intersecting dbSNP129 with probe coordinates (obtained from X:MAP [Yates *et al*. 2007]) using the Galaxy Browser [Giardine *et al*. 2005]). A MySQL database containing the PLIER normalized intensities and CEU [Utah residents with Northern and Western European ancestry] and YRI [Yoruba in Ibadan] genotypes for Phase I+II HapMap SNPs was created. Tables were derived to link SNPs to their nearest like-stranded probeset (to within 500nt), and to associate probesets to the exons they may overlap (transcript and exon tables from Ensembl version 51). A MySQL query was used to create a table containing the splicing index (SI; intensity of a probeset divided by the overall gene intensity) of each probeset for each HapMap individual.

The database was queried to identify significant SI changes of an exonic probeset based on the genotype of a SNP the probeset was associated with (SNP within natural donor/acceptor region of exon). Probesets displaying a step-wise change in mean SI (where the mean SI of the heterozygous group is in between the mean SI values of the two homozygous groups) were identified using a different program script (criteria: the mean SI of homozygous rare and heterozygous groups are < 90% of the homozygous common group). Splicing Index boxplots were created with R, where the x and y-axis are genotype and SI, respectively [Supplementary Figure 1]. These boxplots analyze the effect a SNP has on a particular probeset across all individuals.

SNPs with effects on splicing were validated by q-RT-PCR of lymphoblastoid cell lines. Where available, results were also compared to abnormal splicing patterns present in RNAseq data from tumors carrying these same SNPs (in the ValidSpliceMut database; Shirley *et al*. 2019). SNPs predicted to exhibit nominal effects on splicing (Δ*R_i_* < 1 bit) were included to determine minimal detectable changes by q-RT-PCR.

### Cell Culture & RNA Extraction

EBV-transformed lymphoblastoid cell lines of HapMap individuals with our SNPs of interest (homozygous common, heterozygous and homozygous rare when available) were ordered from the Coriell Cell Repositories (CEU: GM07000, GM07019, GM07022, GM07056, GM11992, GM11994, GM11995, GM12872; YRI: GM18855, GM18858, GM18859, GM18860, GM19092, GM19093, GM19094, GM19140, GM19159). Cells were grown in HyClone RPMI-1640 medium (15% FBS [HyClone], 1% L-Glutamine and 1% Penicillin:streptomycin [Invitrogen]; 37°C, 5% CO_2_). RNA was extracted with Trizol LS (Invitrogen) from 10^6^ cells and treated with DNAase (20mM MgCl_2_ [Invitrogen], 2mM DTT [Sigma-Aldrich], 0.4U/uL RNasin [Promega], 10µg/mL DNase [Worthington Biochemical] in 1x TE buffer) at 37°C for 15 minutes. The reaction was stopped with EDTA (0.05M; 2.5% v/v), and heated to 65°C for 20 minutes, followed by ethanol precipitation (resuspended in 0.1% v/v DEPC-treated 1x TE buffer). DNA was extracted using a Puregene Tissue Core Kit B (Qiagen).

### Design of Real-Time Expression Assay

Sequences were obtained from UCSC and Ensembl. DNA primers used to amplify a known splice form, or one predicted by information analysis, were designed using Primer Express (ABI). DNA primers [Supplementary Table 1] were obtained from IDT (Coralville, IA, USA), and dissolved to 200 uM. Primers were placed over junctions of interest to amplify a single splice form. T_m_ ranged from 58-65°C, and amplicon lengths varied from 69-136nt. BLASTn (Refseq_RNA database) was used to reduce possible cross-hybridization. Primers were designed to amplify the wildtype splice form, exon skipping (if a natural site is weakened), and cryptic site splice forms which were either previously reported (UCSC mRNA and EST tracks) or predicted by information analysis (where *R_i_* cryptic site ≥ *R_i_* weakened natural site).

Two types of reference amplicons were used to quantify allele specific splice forms. These consisted of intrinsic products derived from constitutively spliced exons with the same gene and external genes with high uniformity of expression among HapMap cell lines. Reference primers internal to the genes of interest were designed 1-4 exons adjacent from the affected exon (exons without any evidence of variation in the UCSC Genome Browser), placed upstream of the SNP of interest whenever possible. Two advantages to including an internal reference in the q-RT-PCR experiment include: potential detection of changes in total mRNA levels; and account for inter-individual variation of expression.

External reference genes (excluding the SNP of interest) were chosen based on consistent PLIER intensities with low coefficients of variation in expression among all 176 HapMap individuals. The following external controls were selected: exon 39 of *SI* (PLIER intensity 11.4 ± 1.7), exon 9 of *FRMPD1* (22 ± 2.81), exon 46 of *DNAH1* (78.5 ± 9.54), exon 3 of *CCDC137* (224 ± 25) and exon 25 of *VPS39* (497 ± 76). The external reference chosen for an experiment was matched to the intensity of the probeset within the exon of interest. This decreased potential errors in ΔΔC_T_ values and proved to be accurate and reproducible for most genes.

To control for interindividual variation in expression, we compared expression in HapMap individuals based on their SNP genotypes and familial relatedness. Families with all 3 possible genotypes were available (homozygous common, rare and heterozygous) for 12 of these SNPs (rs1805377, rs2243187, rs2070573, rs2835655, rs2835585, rs2072049, rs1893592, rs6003906, rs1018448, rs13076750, rs16802, and rs8130564). For those families in which all genotypes were not represented, samples from the same ethnic background (YRI or CEU populations) were compared for the missing genotype (N=8; rs17002806, rs2266988, rs1333973, rs743920, rs2285141, rs2838010, rs10190751, rs16994182; individuals with homozygous common and rare genotypes were from the same families for the latter two SNPs). Two SNPs were tested using homozygous individuals from different ethnic backgrounds: rs3747107 (*GUSBP11*) and rs2252576 (*BACE2*). While the splicing impact of rs3747107 was clearly observable by q-RT-PCR, either background or data noise did impact the interpretation of effects of rs2252576.

### PCR and Quantitative RT-PCR

M-MLV reverse transcriptase (Invitrogen) converted 1µg of DNase-treated RNA to cDNA with 20nt Oligo-dT (25µg/mL; IDT) and rRNAsin (Promega). Precipitated cDNA was resuspended in water at 20ng/µL of original RNA concentration. All designed primer sets were tested with conventional PCR to ensure a single product at the expected size. PCR reactions were prepared with 1.0M Betaine (Sigma-Aldrich), and were heated to 80°C before adding Taq Polymerase (Invitrogen). Optimal T_m_ for each primer set was determined to obtain maximum yield.

Quantitative PCR was performed on an Eppendorf Mastercycler ep Realplex 4, a Bio-Rad CFX96, as well as a Stratagene Mx3005P. SYBR Green assays were performed using the KAPA SYBR FAST qPCR kit (Kapa Biosystems) in 10µL reactions using 200µM of each primer and 24ng total of cDNA per reaction. For some tests, SsoFast Eva Green supermix (Bio-Rad) was used with 500µM of each primer instead.

When testing the effect of a SNP, amplification reactions with all primers designed to detect all relevant isoforms (as well as the gene internal reference and external reference) were run simultaneously, in triplicate. C_t_ values obtained from these experiments were normalized against the same external reference using the Relative Expression Software Tool (REST; http://www.gene-quantification.de/rest.html; Pfaffl *et al*. 2002).

### Taqman Assay

Two dual-labeled Taqman probes were designed to detect the two splice forms of *XRCC4* (detecting alternative forms of exon 8 either with or without a 6nt deletion at the 5’ end). Probes were placed over the sequence junction of interest where variation would be near the probe middle [Supplementary Table 1]. The assay was performed on an ABI StepOne Real-Time PCR system using ABI Genotyping Master Mix. Experiment was run in 25µL reactions (300nM each primer, 400nM probe [5’-FAM or TET fluorophore with a 3’ Black Hole quencher; IDT], and 80ng cDNA total). Probes were tested in separate reactions.

### RNAseq Analyses

The previous analyses were extended to include 24 additional, common SNPs for their potential influence on splicing. All SNVs present in ICGC (International Cancer Genome Consortium) patients (Shirley *et al*. 2019) were evaluated by the Shannon Pipeline (SP; Shirley *et al*. 2013) to identify those altering splice site strength. Common SNPs (average heterozygosity > 10% in dbSNP 150) predicted to decrease natural splice site strength by SP (where Δ*R_i_* < -1 bit) were selected. ICGC patients carrying these flagged SNPs were identified, and the expression of the corresponding SNP-containing region in RNAseq was visualized with IGV (Integrated Genome Viewer; https://igv.org). Similar RNAseq reads were grouped using IGV collapse and sort commands, which caused non-constitutive spliced reads to cosegregate to the top of the viewing window. IGV images which did not meet our gene expression criteria (exon affected by the SNP must have ≥5 RNAseq reads present) were eliminated. As this generated thousands of images, we report the analysis of two ICGC patients (DO47132 [Renal Cell Cancer] and DO52711 [Chronic Lymphocytic Leukemia]), chosen randomly, preselecting tissues to increase the likelihood of finding expression in these regions. Images were evaluated sequentially (in order of rsID value) and only concluded once the first 24 SNPs meeting these criteria were found. This type of analysis could not reveal a splicing event to be more abundant in these patients when compared to non-carriers. Nevertheless, splicing information changes resulting from SNPs corresponded to observed alternative and/or other novel splice isoforms. We then queried the ValidSpliceMut database for these SNPs, as abnormal splicing was only flagged in the database when the junction-read or read-abundance counts significantly exceeded corresponding evidence type in a large set of normal control samples (Shirley *et al*. 2019).

## Results

### Selection of Candidate SNPs Affecting Splicing

A publicly available exon microarray dataset was initially used to locate exons affected by SNPs altering splice site strength. A change in the mean SI of a diagnostic probeset in individuals of differing genotypes at the same variant can suggest altered splicing. The increase or decrease in SI is related to the expected impact of the SNP on splicing. For example, an exonic probe which detects a normally spliced mRNA will have decreased SI in the event of skipping. Mean SI may be increased when a probe detects the use of an intronic cryptic splice site. SNPs with strong impact on splicing will distinguish mean SI levels of individuals homozygous for the major versus minor alleles (and with heterozygous genotypes).

There were 9328 HapMap-annotated SNPs within donor/acceptor regions of known exons which contained at least one probeset. Of 987 SNPs that are associated to exonic probesets which differ in mean SI between the homozygous common and rare HapMap individuals, 573 caused a decrease in natural site *R_i_* value. Inactivating and leaky splicing variants (reduction in information content where final *R_i_* ≥ *R_i,minimum_* [minimum functional splice site strength]) both exhibit reduced SI values and were similarly abundant. Thus, both severe and moderate splicing mutations with reduced penetrance and milder molecular phenotypes were detected, consistent with Mendelian disorders (von Kodolitsch *et al*. 1999; von Kodolitsch *et al*. 2006).

Of the SNPs associated with significant changes in *R_i_* (termed Δ*R_i_*), 9328 occurred within the natural splice sites of exons detectable with microarray probesets. We initially focused on 21 SNPs on chromosome 21 (0.23% total, 18.8% of chr21) and 34 on chromosome 22 (0.36% of total, 14.5% of chr22) associated with step-wise decreases in probeset intensity at each genotype. Seven of the chr21 SNPs and 9 of the chr22 SNPs caused information changes with either natural splice site Δ*R_i_* ≥ 0.1 bits, or cryptic site(s) with an *R_i_* value comparable to a neighbouring natural site, and in which mRNA or EST data supported use of the cryptic site. These SNPs included: rs2075276 [*MGC16703*], rs2838010 [*FAM3B*], rs3747107 [*GUSBP11*], rs2070573 [*C21orf2*], rs17002806 [*WBP2NL*], rs3950176 [*EMID1*], rs1018448 [*ARFGAP3*], rs6003906 [*DERL3*], rs2266988 [*PRAME*], rs2072049 [*PRAME*], rs2285141 [*CYB5R3*], rs2252576 [*BACE2*], rs16802 [*BCR*], rs17357592 [*COL6A2*], rs16994182 [*CLDN14*], and rs8130564 [*TMPRSS3*].

The minimum information change for detecting a splicing effect by expression microarray is constrained by several factors. Detection of splice isoforms can be limited by genomic probeset coverage, which cannot distinguish alternative splicing events in close proximity (see Figure 1A). Even where genotype-directed SI changes are very distinct, some individuals with the common allele have equivalent SI values to individuals with the rare allele (rs2070573 [Figure 2] and rs1333973 [Figure 3]). In some cases, the number of individuals with a particular genotype is insufficient for statistical significance (rs2243187; Supplementary Figure 1.4). Although exon microarrays can be used to find potential alternate splicing and give support to our predictions, it became necessary to validate the microarray predictions by q-RT-PCR, TaqMan assays, and with RNAseq data from SNP carriers.

**Figure 1.**
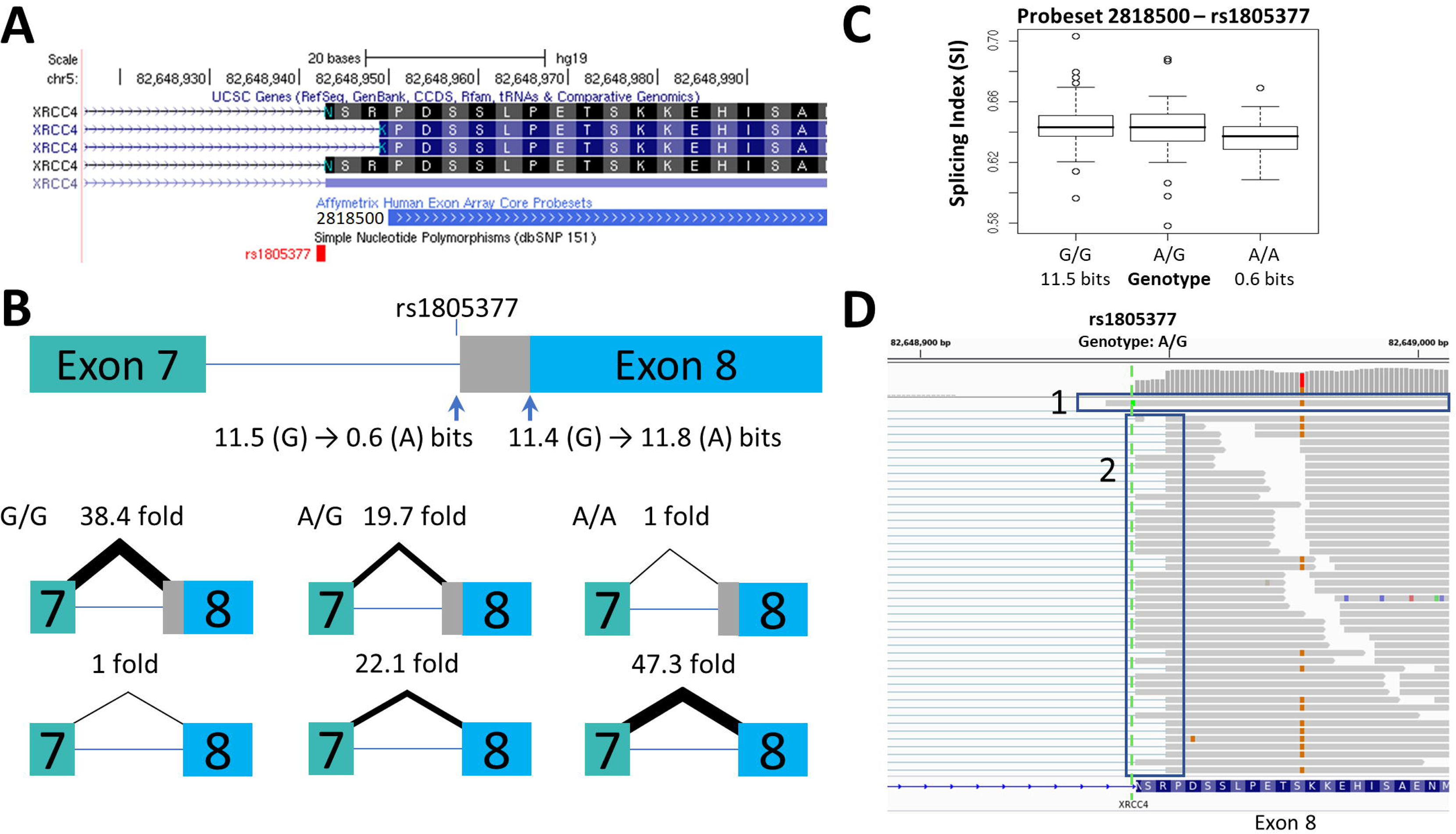
Splicing Impact of rs1805377 (*XRCC4*). The natural acceptor of *XRCC4* exon 8 is abolished by rs1805377 (11.5 -> 0.6 bits) while simultaneously strengthening a second exonic cryptic acceptor 6nt downstream (11.4 to 11.8 bits), resulting in a 6nt deletion in the mRNA. **(A)** Both of these acceptor sites have been validated in GenBank mRNAs, i.e. NM_022406 and NM_003401. **(B)** The relative abundance of the two splice forms was determined by q-RT-PCR. The weaker rs1805377 A/A allele (0.6 bit acceptor) was used ∼47 fold less frequently than the cryptic downstream acceptor (11.8 bits). **(C)** The two splice isoforms cannot be distinguished by the exon microarray as the upstream probeset (ID 2818500) does not overlap the variable region, though the average expression of the rs1805377 A/A allele is reduced. **(D)** ValidSpliceMut flagged this mutation for intron retention, which can be observed in the RNAseq of heterozygous ICGC patient DO27779 [Box 1]. Use of both acceptor sites is also evident [Box 2]. For more detail for this and all of the other SNPs analyzed, refer to Supplementary Figure 1.

**Figure 2.**
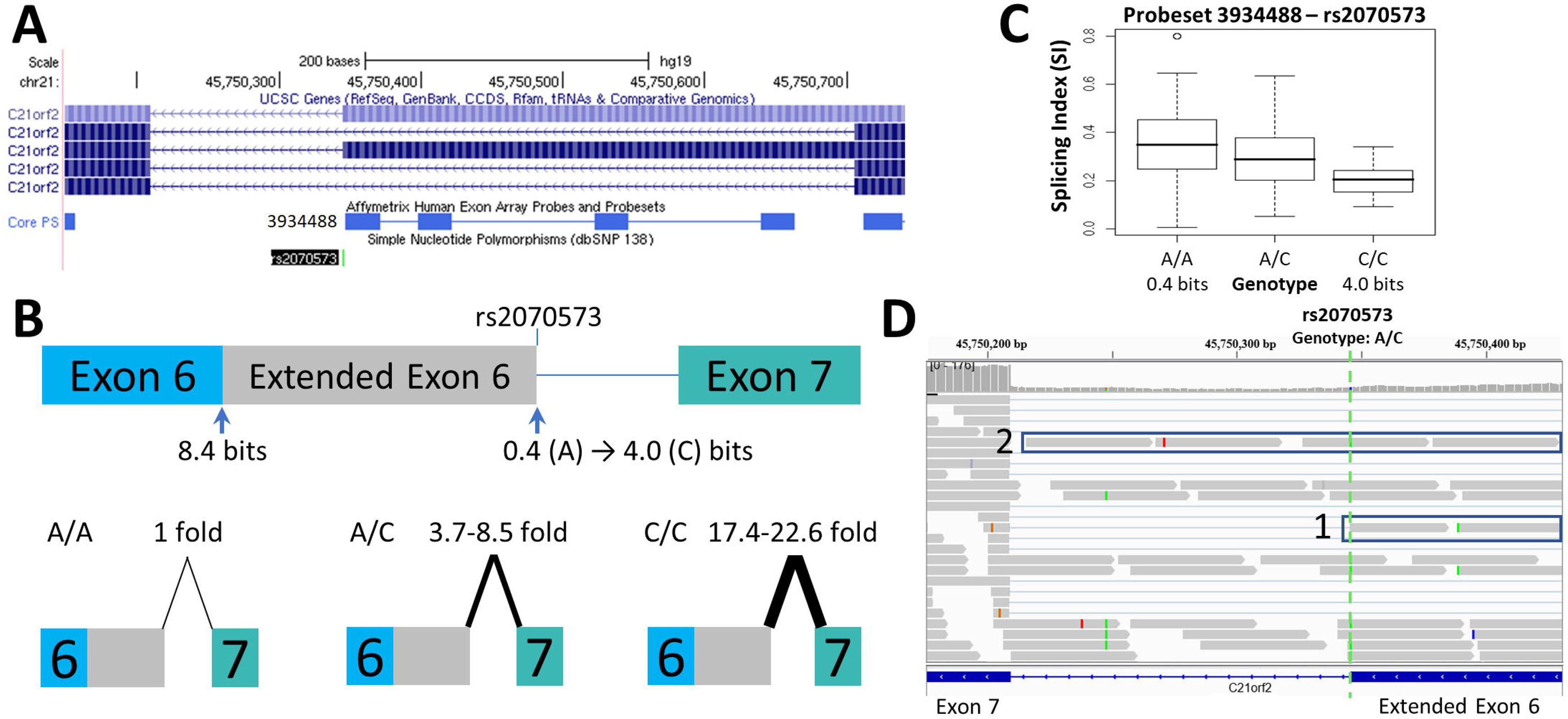
Splicing Impact of rs2070573 (*C21orf2*). **(A)** The SNP rs2070573 is a common polymorphism which alters the first nucleotide of the extended form of *C21orf2* exon 6. **(B)** The donor site is strengthened by the presence of the C allele (*R_i_* 0.4 to 4.0 bits; A>C) and its use extends the exon by 360 nt. Q-RT-PCR found a ∼4-9-fold and ∼17-23 fold increase in the extended exon 6 splice form in the A/C and C/C cell lines tested, respectively. **(C)** The exon microarray probeset which detects the extension (ID 3934488) shows a step-wise increase in SI with C-allele individuals which supports the q-RT-PCR result. **(D)** The variant was present in ValidSpliceMut, which associated the A-allele with an increase in total intron retention (6 patients flagged for total intron retention read abundance; p=0.019 [average over all patients]). This image displays sequence read distributions in the RNAseq data of TCGA BRCA patient, TCGA-BH-A0H0, who is heterozygous for rs2070573. The IGV panel indicates reads corresponding to total intron 6 retention [Box 1] and which extend beyond the constitutive donor splice site of exon 6 into the adjacent intron [Box 2]. All 4 reads which extend over the exon splice junction are derived from the G allele (strong binding site; not all visible in panel D).

**Figure 3.**
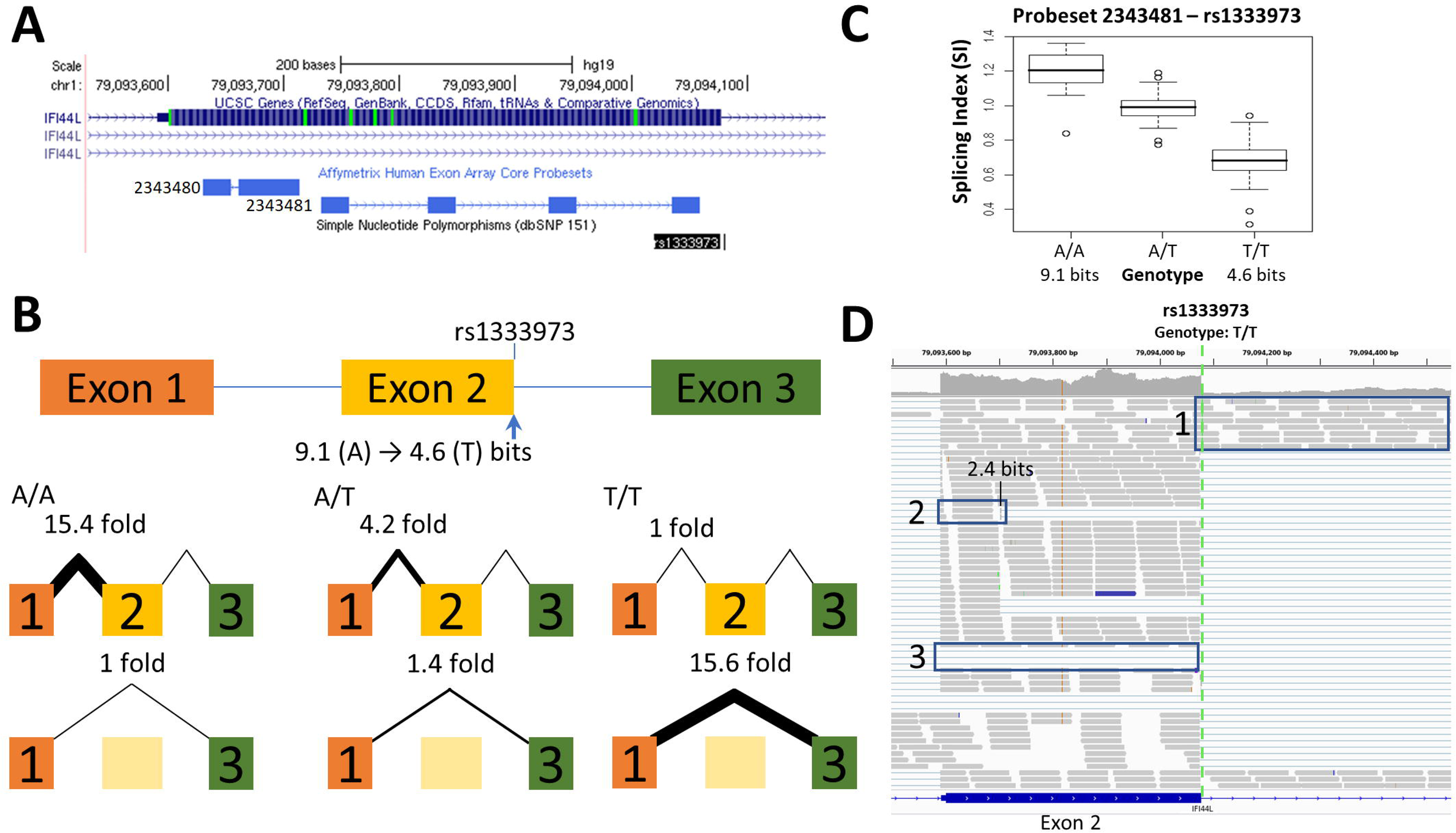
Splicing Impact of rs1333973 (*IFI44L*). **(A)** The natural donor of *IFI44L* exon 2 is weakened from 9.1 to 4.6 bits (A>T) by rs1333973, increasing the frequency of exon 2 skipping and other events. **(B)** By q-RT-PCR, skipping was found to be 15.6-fold higher while normal splicing was 15.4-fold lower in A homozygotes (relative to T homozygotes). **(C)** Exon microarray data strongly supports the q-RT-PCR findings. **(D)** ICGC patient DO6354 is homozygous for the T allele, which resulted in exon 2 skipping [Box 3], failure to recognize the exon 2 donor causing total retention of intron 2 [Box 1], and activation of an upstream exonic 2.4 bit cryptic donor 375nt from the affected site [Box 2].

We report q-RT-PCR validation studies for 13 of the 16 SNPs (q-RT-PCR primers could not be designed for rs16994182, rs2075276 and rs3950176), along with 9 other candidate SNPs from our previous information theory-based analyses (Nalla and Rogan, 2005): rs1805377 [*XRCC4*], rs2243187 [*IL19*], rs2835585 [*TTC3*], rs2865655 [*TTC3*], rs1893592 [*UBASH3A*], rs743920 [*EMID1*], rs13076750 [*LPP*], rs1333973 [*IFI44L*] and rs10190751 [*CFLAR*].

After amplification of known and predicted splice forms [Supplementary Table 1], 15 SNPs showed measurable changes in splicing consistent with information-theory predictions. Ten increase alternate splice site use (2 of which increased strength of cryptic site, 8 activated an unaffected pre-existing cryptic site), 6 affect exon inclusion (5 increased exon skipping), 3 increased activation of an alternative exon, and 4 decreased overall expression levels. Altered splicing could not be validated for 6 SNPs, however experimental analyses of 3 of the 5 SNPs where Δ*R_i_* < 1 bit were hampered by high inter-individual variability in expression.

Changes in splice site information were used to predict observed differences in splice isoform levels (Table 1). Figures 1–3 and Supplementary Figure 1 indicate the experimentally-determined splicing effects for each SNP, a UCSC Genome Browser image of the relevant region, boxplots showing exon microarray expression levels of each allele for the relevant probesets, and an IGV image of the RNAseq results for an individual tumour carrying the SNP. Abundance of the aberrant splice forms measured by q-RT-PCR (relative to an internal gene reference) are indicated in Table 2. Changes in predicted splice site strength were consistent with results measured by q-RT-PCR for 12 out of the 15 SNP (exceptions were rs2070573, rs17002806, and rs2835585). Variants predicted to reduce strength ≥ 100-fold were found to reduce expression by 38- to 58-fold, the variance falling within the margin of measurement error. Modest natural splice site affinity changes predicted to be < 8-fold (ΔR_i_ < 2.8) did not consistently result in detectable changes in splicing. In some instances, lower abundance splice forms were observed (i.e. rs2835585 altered exon skipping levels by up to 8.8-fold; nevertheless, the normal splice form predominated).

**Table 1:**
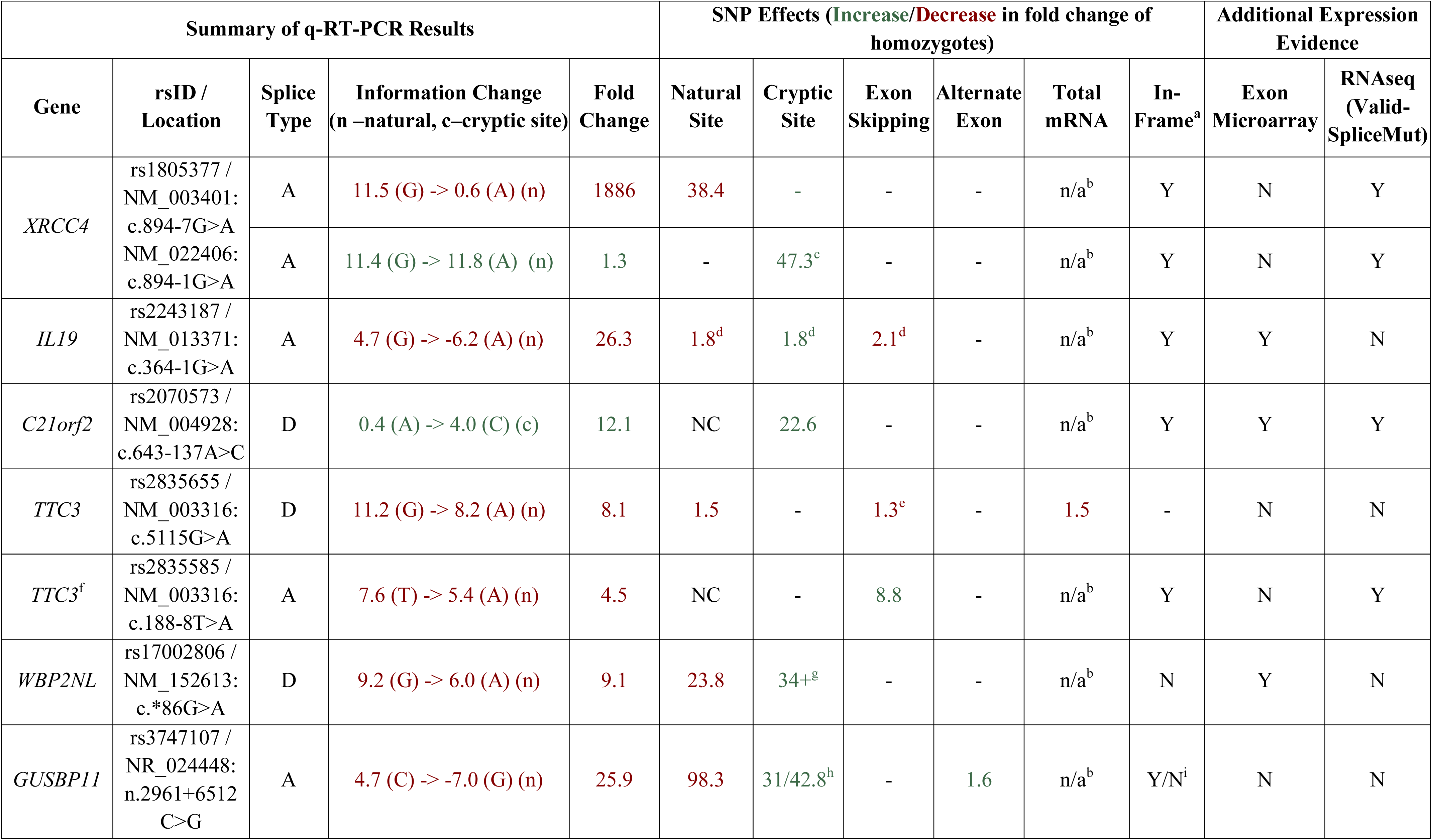
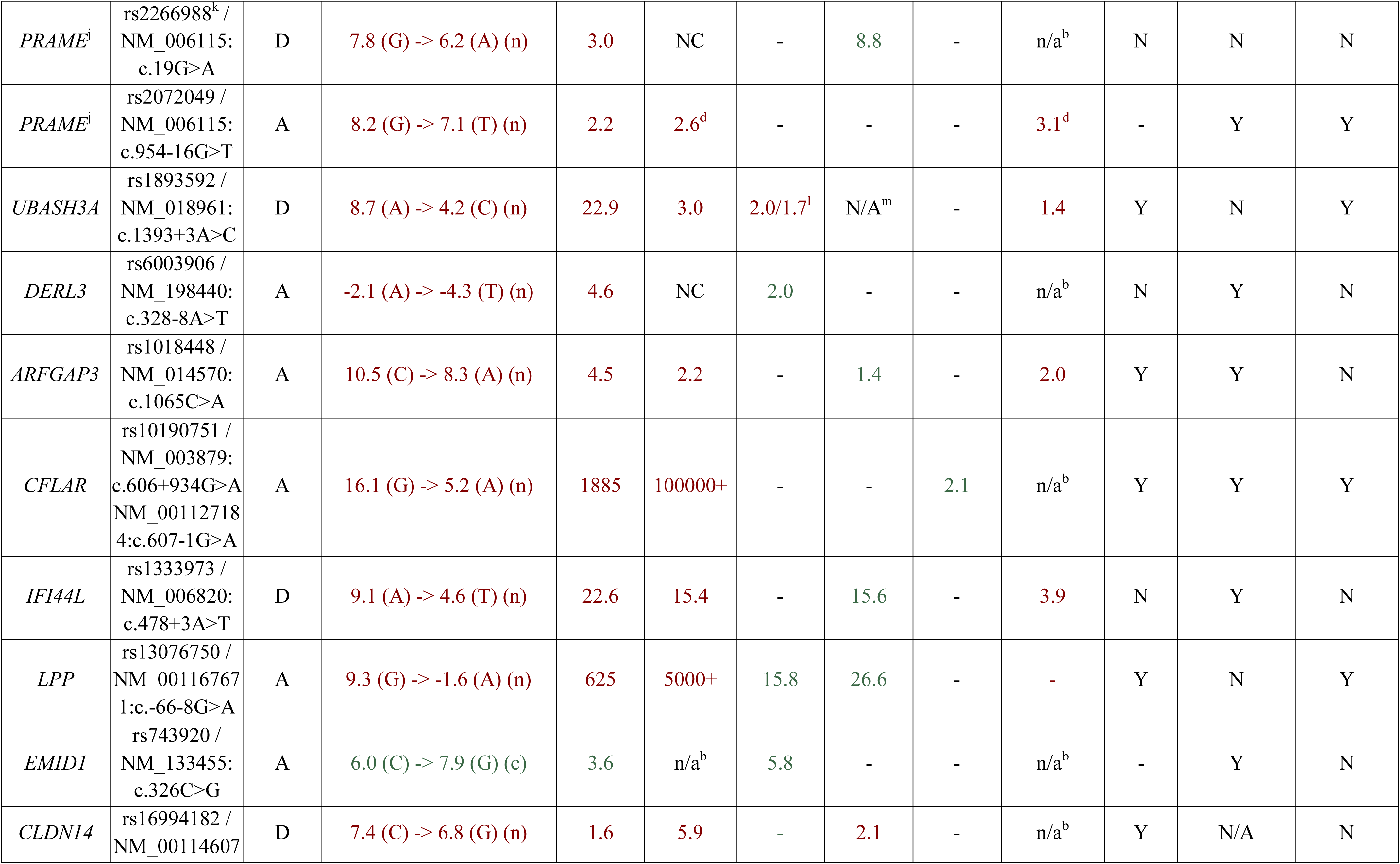
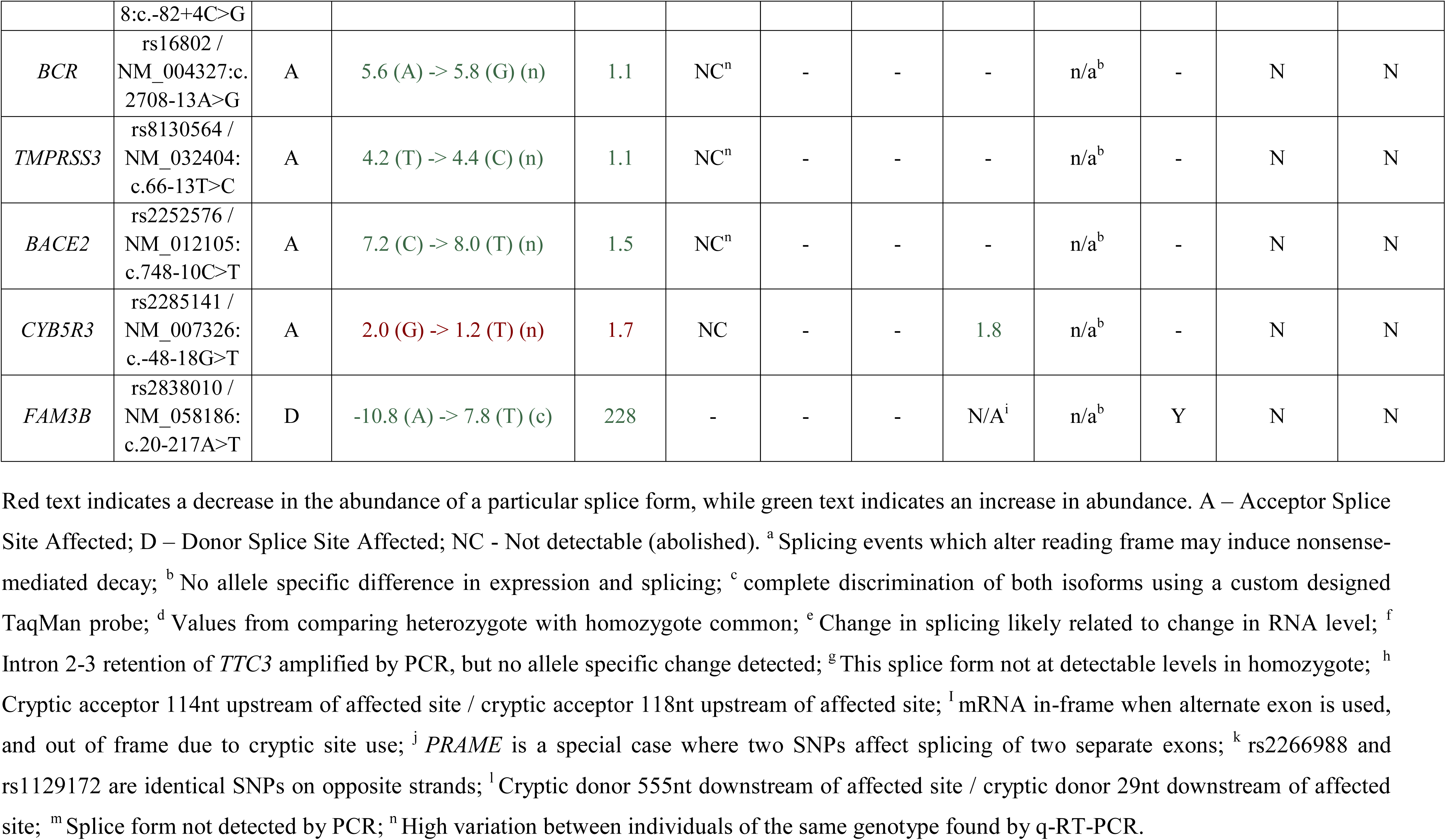
Summary of q-RT-PCR Results.

**Table 2:**
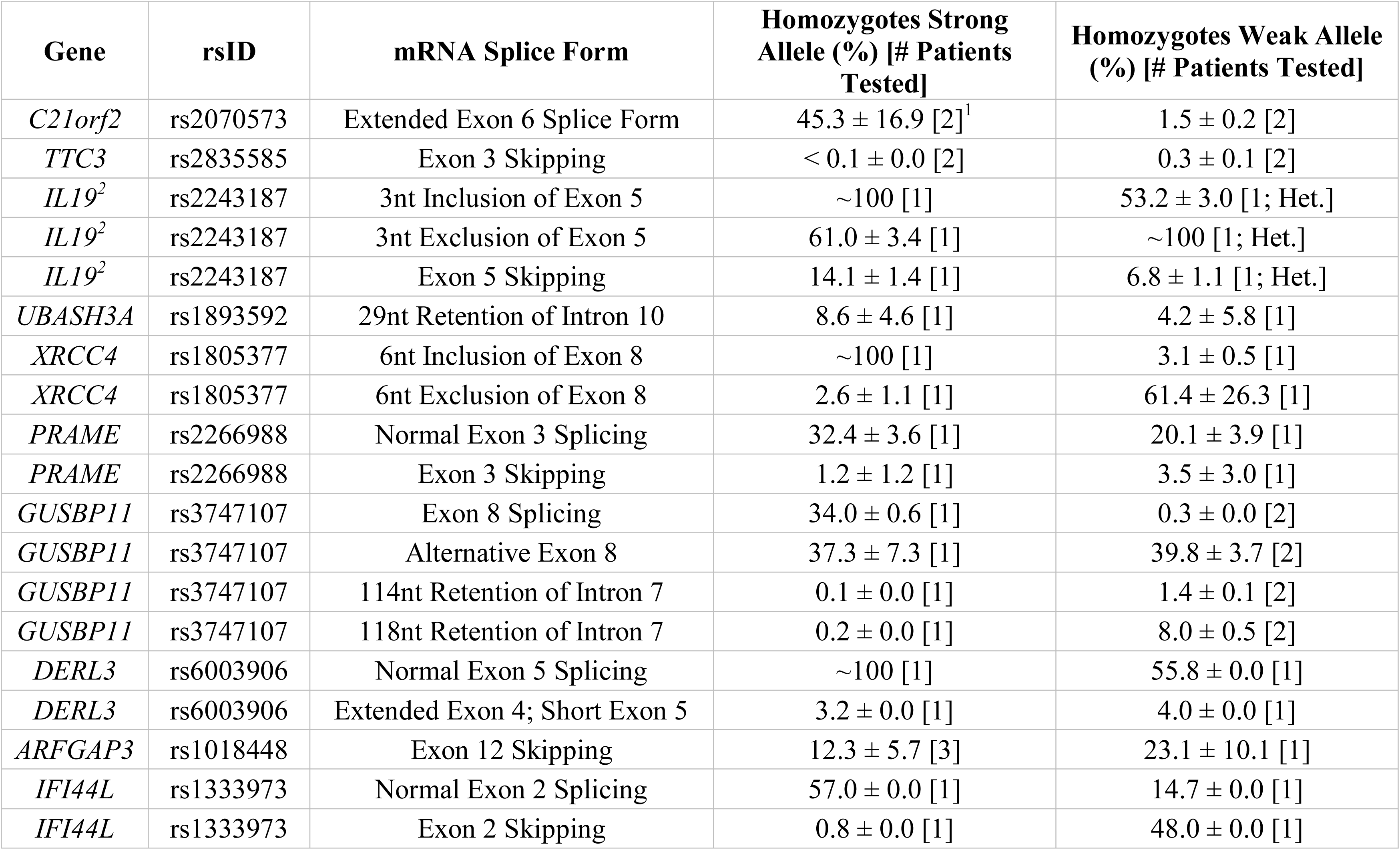
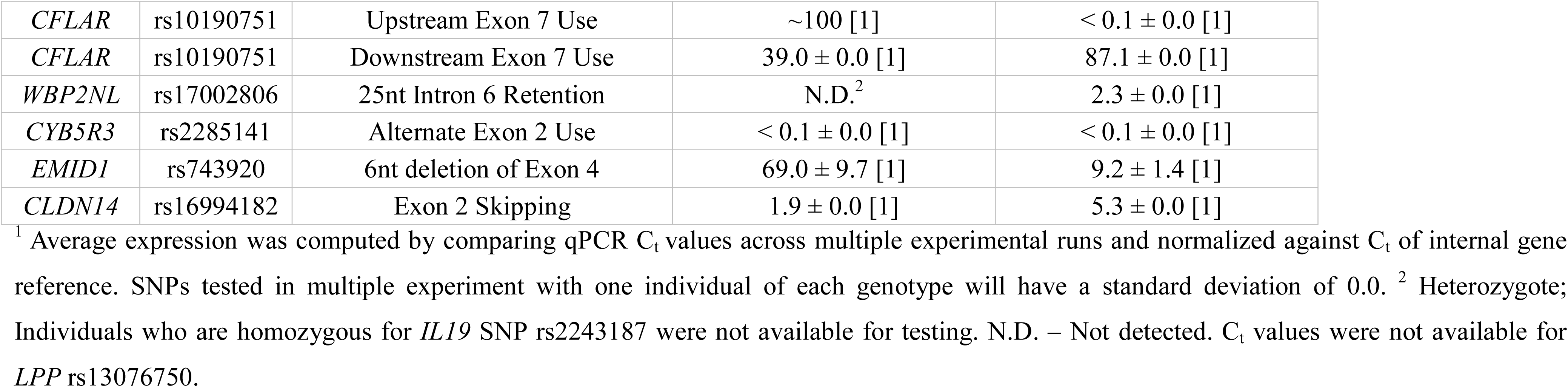
Abundance of mRNA Splice Forms Relative to Internal Gene Reference.

### SNPs Affecting Cryptic Site Strength and Activity

Increased cryptic site use coinciding with a decrease in natural site strength (Table 1) was validated for: rs1805377 (Figure 1); rs2243187 [Supplementary Figure 1.4]; rs3747107 [Supplementary Figure 1.8]; rs17002806 [Supplementary Figure 1.13]; rs6003906 [Supplementary Figure 1.15]; and rs13076750 [Supplementary Figure 1.12]. rs2070573 (Figure 2) and rs743920 [Supplementary Figure 1.7] strengthened cryptic splice sites resulting in increased use of these sites. Despite the difference in strength between the natural and cryptic sites affected by rs743920, the upstream 2.4 bit site was used more frequently (Table 2). Both *IL19* and *XRCC4* regions tested showed preference to the upstream acceptor as well, which is consistent with the processive mechanism documented to recognize acceptor splice sites (Robberson *et al*. 1990).

### SNPs Affecting Exon Inclusion

SNPs that reduced natural site strength (Δ*R_i_* from 1.6 to 10.9 bits) increased exon skipping from 2 to 27-fold for homozygotes of differing genotypes of rs2835585 [Supplementary Figure 1.21], rs1018448 [Supplementary Figure 1.3], rs1333973 (Figure 3), rs2266988 [Supplementary Figure 1.9], and rs13076750 [Supplementary Figure 1.12]. The exon microarray probesets for rs1018448 and rs1333973 detect decreased expression by genotype, which is consistent with increased exon skipping. Changes of average SI values did not correspond as well to specific genotypes for rs2835585 (*TTC3)*, rs2266988 (*PRAME)* and rs13076750 (*LPP)*, possibly due to increased cryptic site use (*LPP*) or large differences in the abundance of constitutive and skipped isoforms (*PRAME*, *TTC3*).

### SNPs Promoting Alternate Exon Use

SNP-related decreases in natural splice site strength may promote the use of alternative exons up or downstream of the affected exon. rs10190751 [Supplementary Figure 1.2] is known to modulate the presence of the shorter c-FLIP(S) splice form of *CFLAR* (Ueffing *et al*. 2009). The use of this exon differed by 2^17^ fold between the strong and weak homozygotes tested, which was reflected by the expression microarray result. By q-RT-PCR, the *CFLAR* (L) form using an alternate downstream exon was found to be 2.1-fold more abundant in the homozygote with the weaker splice site. rs3747107 [Supplementary Figure 1.8] and rs2285141 [Supplementary Figure 1.20] exhibit evidence of an increased preference by q-RT-PCR for activation of an alternate exon, though the microarray results for the corresponding genotypes for both SNPs were not significantly different.

### SNP-directed effects on mRNA levels

A change in the strength of a natural site of an exon can affect the quantity of the processed mRNA (Caminsky *et al*. 2014). This decrease in mRNA could be caused by nonsense mediated decay (NMD), which degrades aberrant transcripts that would result in premature protein truncation (Cartegni *et al*. 2002). Of the 22 SNPs tested, 2 showed a direct correlation between a decrease in natural splice site strength, reduced amplification of the internal reference by q-RT-PCR (of multiple individuals) and a decreasing trend in expression by genotype by microarray: rs2072049 [Supplementary Figure 1.9] and rs1018448 [Supplementary Figure 1.3], although these differences do not meet statistical significance.

### SNPs with pertinent splicing effects detected by RNAseq

Evidence for impact on splicing of the previously described SNPs was also assessed in TCGA and ICGC tumours by high throughput expression analyses. Splicing effects of these variants detected by q-RT-PCR and RNAseq were concordant in 80% of cases (N=16 of 20 SNPs), while impacts of 10% of SNPs (N=2) were partially concordant as a result of inconsistent activation of cryptic splice sites (Supplementary Figures 1.8D and 1.10D). Several isoforms predicted by information analysis of these SNPs were present in complete transcriptomes, but were undetectable by q-RT-PCR or expression microarrays. Examples include a 4.9 bit cryptic site activated by rs3747107 located 2 nucleotides from the natural splice site [Supplementary Figure 1.8D], exon skipping by rs1893592 [Supplementary Figure 1.10D], and a cryptic exon activated by rs2838010 [Supplementary Figure 1.16C]. Processed mRNAs that were not detected by q-RT-PCR may have arisen as a result of a lack of sensitivity of the assay, to NMD (which could mask detection of mis-splicing), to a deficiency of an undefine trans-acting splicing factor, or to design limitations in the experimental design. Another possibility is that the discordant splicing patterns of these two SNPs based could potentially be related to differences in tissue origin, since only the RNAseq findings were tumor-derived, whereas results obtained by the other approaches were generated from RNA extracted from lymphoblastoid cell lines. Cell culture conditions such as cell density and phosphorylation status can affect alternative splicing patterns (Li *et al*., 2006; Szafranski *et al*., 2014). These conditions, however, have not been studied in cases of allele-specific, sequence differences at splice sites or cis-acting regulatory sites that impact splice site selection. Considering the high level of concordance of splicing effects for the same SNPs in uncultured and cultured cells, it seems unlikely that culture conditions significantly impacts the majority of allele-specific, alternatively spliced isoforms. Our information theory-based analyses show that the dominant effect of SNP genotypes is to dictate common changes in splice site strength regardless of cell origin.

The results obtained from q-RT-PCR and RNAseq data for rs2070573, rs10190751, rs13076750, rs2072049, rs2835585, rs1893592 and rs1805377 were complementary to findings based on RNAseq [Supplementary Table 2]. RNAseq data can reveal potential allele-specific alternate splicing events that were not considered at the primer design phase of the study, while q-RT-PCR is more sensitive and can reveal less abundant alternative splice forms. A weak 0.4 bit splice site associated with rs2070573 was less abundant than the extended isoform (Figure 2) by both q-RT-PCR and exon microarray, however ValidSpliceMut also revealed increased total *C21orf2* intron 6 retention in 5 tumours with this allele. Similarly, rs10190751 was flagged for intron retention in 29 tumours, which was not evident by the other approaches. The long form of this transcript (c-FLIP[L]) in homozygous carriers of this SNP was twice as abundant by q-RT-PCR than the shorter allele, associated with the weak splice site. rs13076750 activates an alternate acceptor site for a rare exon that extends the original exon length by 7 nucleotides. The exon boundary can also extend into an adjacent exon, based on RNAseq of 8 tumors carrying this SNP. Expression was decreased in the presence of a 6.2 bit splice site derived from a rs2072049 allele that weakens the natural acceptor site of the terminal exon of *PRAME*. The actual cause of diminished expression is likely to have been related to NMD from intron retention. ValidSpliceMut showed intron retention to be increased in rs2835585, whereas increased exon skipping for the allele with the weaker splice site was demonstrated by q-RT-PCR. rs1893592 caused significant intron retention in all tumours (N=9), with exon skipping present in 3 diffuse large B-cell lymphoma patients, which was not detected by q-RT-PCR. Finally, rs1805377 was associated with the significant abundance of read sequences indicating *XRCC4* intron 7 retention by RNAseq (N=32), however this isoform could not be distinguished by the primers designed for q-RT-PCR and by TaqMan assay.

Alternative splicing events detected by RNAseq that were not evident in either q-RT-PCR or microarray studies included exon skipping induced by rs743920 [Supplementary Figure 1.7D], activation of a pre-existing cryptic splice site by rs1333973 (Figure 3), and intron retention by rs6003906 [Supplementary Figure 1.15D]. rs743920 creates an exonic hnRNP A1 site (*R_i_* = 2.8 bits) distant from the natural site which may compromise exon definition (Mucaki *et al*. 2013; Peterlongo *et al*. 2015) and may explain the SNP-associated increase in exon skipping. Exon definition analyses of total exon information (*R_i,total_*) also predicted the cryptic isoform arising from rs1333973 to be the most abundant (*R_i,total_* = 9.4 bits).

### Allele-specific mRNA splicing for other SNPs identified through RNAseq

A distinct set of 24 high population frequency SNPs were also evaluated for their potential impact on mRNA splicing by RNAseq analysis of ICGC patients. Those resulting in significantly decreased natural splice site strength (Δ*R*_i_ < -1 bit) were analyzed for SNP-derived alternative splicing events. SNPs fulfilling these criteria expressed at sufficient levels over the region of interest were: rs6467, rs36135, rs154290, rs166062, rs171632, rs232790, rs246391, rs324137, rs324726, rs448580, rs469074, rs518928, rs624105, rs653667, rs694180, rs722442, rs748767, rs751128, rs751552, rs752262, rs832567, rs909958, rs933208, and rs1018342 (Table 3). Splicing was predicted to be leaky for all natural splice sites affected by these SNPs (Rogan *et al*. 1998; *R_i,final_* ≥ 1.6 bits), where reduction in *R_i_* values ranged from 1.1 to 3.3 bits.

**Table 3:**
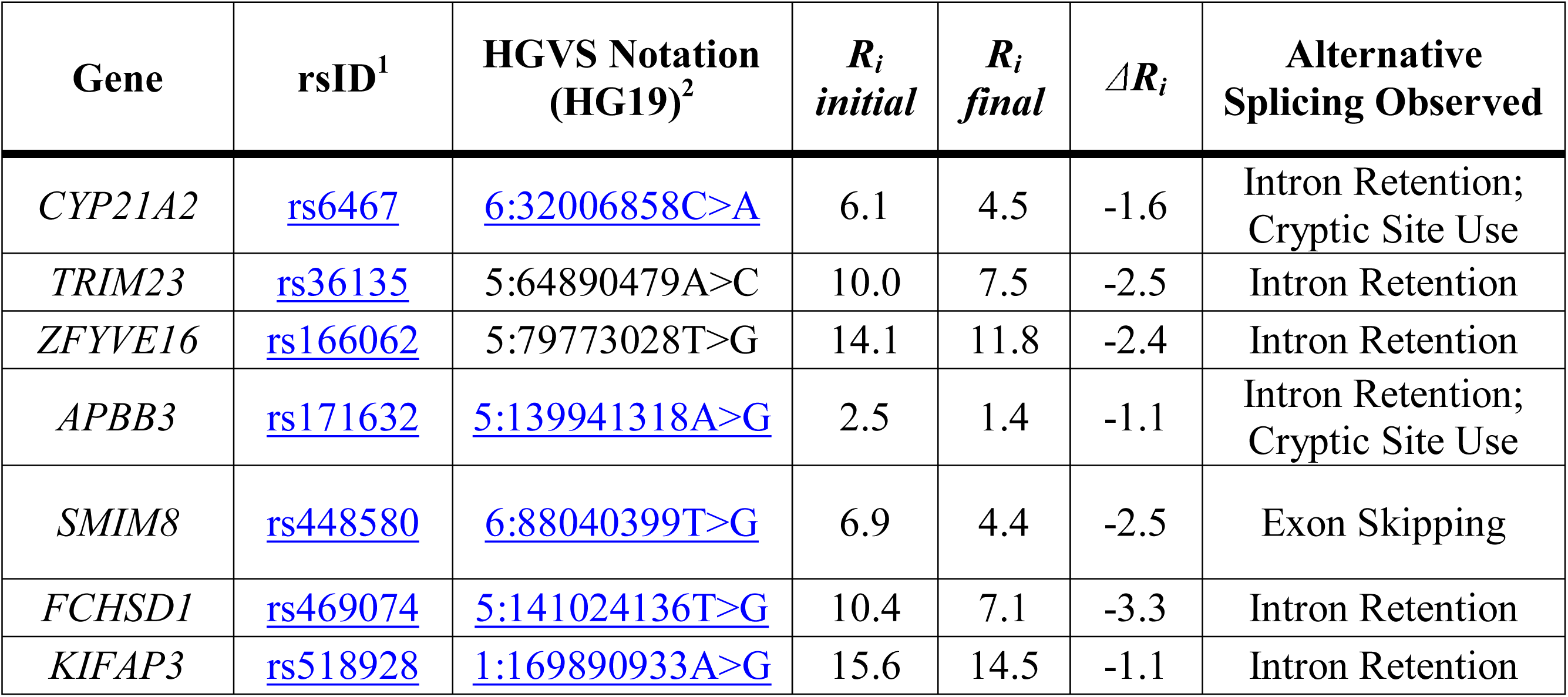
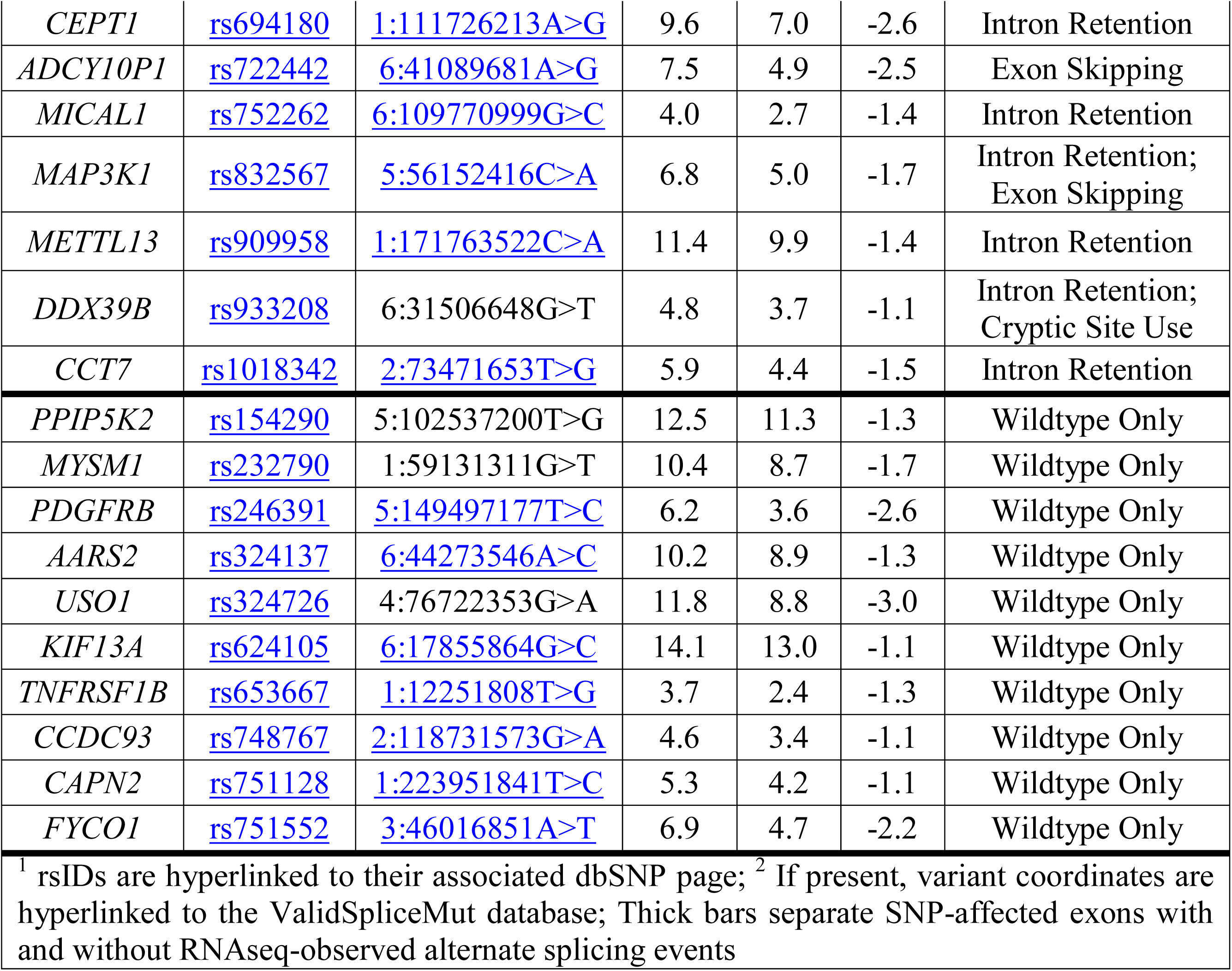
RNAseq Analysis of Natural Splice Sites Weakened by Common SNPs.

Alternative mRNA splicing was observed in 14 SNPs: rs6467, rs36135, rs166062, rs171632, rs448580, rs469074, rs518928, rs694180, rs722442, rs752262, rs832567, rs909958, rs933208, rs1018342; Table 3). Reads spanning these regions revealed intron retention (N=12), activation of cryptic splicing (N=4), and complete exon skipping (N=3). Eleven of these SNPs (79%) exhibited splicing patterns that significantly differed from the control alleles, and were therefore present in ValidSpliceMut. Interestingly, ValidSpliceMut contained entries for 7 of 10 SNPs where alternative splicing had not been found in the two patients reported in Table 3 (rs246391, rs324137, rs624105, rs653667, rs748767, rs751128, rs751552). The observed significant splicing differences for these SNPs occurred in distinct tumor types, consistent with tissue-specific effects of these SNPs on splicing.

### Limited corroboration of SNP-related predictions

Anticipated effects of the SNPs on splicing were not always confirmed by expression studies. Aside from incomplete or incorrect predictions, both design and execution of these studies as well as uncharacterized tissue specific effects could provide an explanation for these discrepancies. Furthermore, these undetected splicing events may have been targeted for NMD, however expression was not compared with mRNA levels from cells cultured with an inhibitor of protein translation. Stronger pre-existing cryptic sites were, in some instances, not recognized nor was isoform abundance changed. These include: rs1893592 (6.4 and 5.2 bit cryptic donor sites 29 and 555nt downstream of the affected donor); rs17002806 (a 5.7 bit site 67nt downstream of the natural); rs3747107 (creates a 4.9 bit cryptic site 2nt downstream [observed by RNAseq; Supplemental Figure 1.8 D]); and rs2835585 (5.8 and 5.9 bit cryptic sites 60 and 87nt upstream of the natural site). SNPs with modestly decreased natural site strength (0.2 to 4.5 bits) did not consistently result in exon skipping (for example, rs1893592, rs17002806 and rs2835655).

Six SNPs predicted to disrupt natural splice sites could not be confirmed. Splicing effects were not identified in the 4 SNPs where the information change was < 1 bit (< 2-fold). Genetic variability masked potential splicing effects of 3 of these SNPs, including rs16802, rs2252576 and rs8130564 (Table 1). PCR primer sets designed for *COL6A2* exon 21 (affected by rs17357592) did not produce the expected amplicon. Interpreting the results for rs16994182 (*CLDN14*) was complicated by the lack of a suitable internal reference. As *CLDN14* consists of 3 exons, any internal reference covering the affected second exon cannot parse whether differences in exon 2 expression were caused by the SNP or by general expression changes.

The T allele rs2838010 was predicted to activate a donor splice site of a rare exon in IVS1 of *FAM3B* (GenBank Accession AJ409094). The cryptic pseudoexon was neither detected by RT-PCR nor expression microarray [Supplemental Figure 1.16]. Interestingly, this exon is expressed in a malignant lymphoma patient who is a carrier for this genotype (ICGC ID: DO27769; [Supplemental Figure 1.16C]). Although the T allele is probably required to activate the pseudoexon, additional unknown splicing-related factors appear to be necessary.

## Discussion

Predicted SNP alleles that alter constitutive mRNA splicing are confirmed by expression data, and appear to be a common cause of alternative splicing. The preponderance of leaky splicing mutations and cryptic splice sites, which often produce both normal and mutant transcripts, is consistent with balancing selection (Nuzhdin *et al*. 2004) or possibly with mutant loci that contribute to multifactorial disease. Minor SNP alleles are often found in > 1% of populations (Janosíková *et al*. 2005). This would be consistent with a bias against finding mutations that abolish splice site recognition in dbSNP. Such mutations are more typical in rare Mendelian disorders (Rogan *et al*. 1998).

Exon-based expression microarrays and q-RT-PCR techniques were initially used to confirm the predicted impact of common and rare SNPs on splicing. Results were subsequently confirmed using RNAseq data for some of these SNPs (Viner *et al*. 2014; Dorman *et al*. 2014, Shirley *et al*. 2019). However, exon skipping due to rs1893592 was not consistently seen in all carriers. Although detected only in one type of tumor, this event may not be tissue specific, since 5 patients with the same genotype did not exhibit this isoform. Nevertheless, exon skipping was also observed in malignant lymphoma [Supplementary Figure 1.10D]). Intron retention in rs1805377 carriers was evident in only 22% of tumors. Increased total intron retention may be due to failure to recognize exons due to overlapping strong splice sites (Rogan *et al*. 2003; Vockley *et al*. 2000).

The splicing impacts of several of these SNPs have also been implicated in other studies. rs10190751 modulates the FLICE-inhibitory protein (c-FLIP) from its S-form to its R-form, with the latter having been linked to increased lymphoma risk (Ueffing *et al*. 2009). We observed the R-form to be twice as abundant for one of the rs10190751 alleles. Increased exon skipping attributed to rs1333973 has been reported in RNAseq analysis of *IFI44L* (Zhao *et al*. 2013 [a]), which has been implicated in reduced antibody response to measles vaccine (Haralambieva *et al*. 2017). The splicing impact of *XRCC4* rs1805377 has been noted previously (Nalla and Rogan 2005). This SNP has been implicated with an increased risk of gastric cancer (Chiu *et al*. 2008), pancreatic cancer (Ding and Li, 2015) and glioma (Zhao *et al*. 2013 [b]). Similarly, the potential impact of rs1893592 in *UBASH3A* has been recognized (Kim *et al*. 2015) and is associated with arthritis (Liu *et al*. 2017) and type 1 diabetes (Ge and Concannon, 2018). Hiller *et al*. (2006) described the 3nt deletion caused by rs2243187 in *IL19* but did not report increased exon skipping. rs743920 was associated with change in *EMID1* expression (Ge *et al*. 2005), however its splicing impact was not recognized. Conversely, studies linking *TMPRSS3* variants to hearing loss did not report rs8130564 to be significant (Lee *et al*. 2013; Chung *et al*. 2014). Interestingly, rs2252576 (in which we did not find a splicing alteration) has been associated to Alzheimer’s dementia in Down syndrome (Mok *et al*. 2014).

rs2835585 significantly increased exon skipping in *TTC3*, however normal expression levels at the affected exon junction were not significantly altered. This was most likely due to the large difference in abundance between the constitutive and skipped splice isoforms (Table 2). The skipped isoform does not disrupt the reading frame and the affected coding region has not been assigned to any known protein domain (Tsukahara *et al*. 1996, Suizu *et al*. 2009). It is unclear whether allele-specific, exon skipping in this instance would impact TTC3 protein function or activity.

Why are so few natural splice sites strengthened by SNP-induced information changes? Most such changes would be thought to be neutral mutations, which are ultimately lost by chance (Fisher, 1930). Those variants which are retained are more likely to confer a selective advantage (Li, 1967). Indeed, the minor allele in rs2266988, which strengthens a donor splice site by 1.6 bits at the 5’ end of the open reading frame in *PRAME* and occurs in 25% of the overall population (∼50% in Europeans). Several instances of modest changes in splice site strength that would be expected to have little or no impact, in fact, alter the degree of exon skipping.

Allele frequency can significantly vary across different populations, which can be indicative of gene flow and migration of a population (Cavalli-Sforza and Bodmer, 1971) as well as, in the case of splicing variants, genetic load and fitness in a population (Rogan and Mucaki, 2011). The frequencies of several of the variants presented here are significantly different between ethnic and geographically defined populations (Tables 1 and 3). We examined allele frequencies of these variants in sub-populations in both HapMap and dbSNP version 153. For example, representation of the alleles of the *XRCC4* variant rs1805377 (where its A-allele leads to a 6nt deletion of the gene’s terminal exon) differs between Caucasians and Asians (for the G- and A-alleles, respectively). Different linkage disequilibrium patterns of this variant occur in Han Chinese (CHB) and Utah residents with Northern and Western European ancestry (CEU) populations (Zhao *et al*. 2013). Similar differences in SNP population frequency include *EMID1* rs743920 (in HapMap: G-allele frequency is 47% in CHB, but only 7% in CEU and 10% in northern Swedish cohorts). This is consistent with dbSNP (version 153) where it is present in 72% in Vietnamese, but only 16% of a northern Sweden cohort). In *BACE2*, rs2252576, the T-allele is most prevalent at 84% in Yoruba in Ibadan, Nigeria populations (YRI), but only 8% in CHB. In *FCHSD1*, rs469074 the frequency of the G-allele is 37% in YRI and <1% in CHB. Some SNPs were exclusively present in a single population in the HapMap cohort (e.g. only the YRI population is polymorphic for *IL19* rs2243187, *WBP2NL* rs17002806, *DERL3* rs6003906, and *CLDN14* rs16994182).

Because of their effects on mRNA splicing, these differences in allele frequency would be expected to alter the relative abundance of certain protein isoforms in these populations. We speculate about whether isoform-specific representation among populations influences disease predisposition, other common phenotypic differences, or whether they are neutral. We suggest that SNPs decreasing constitutive splicing while increasing mRNA isoforms which alter the reading frame would be more likely to result in a distinct phenotype. q-RT-PCR experiments confirmed 5 SNPs which increased the fraction of mRNA splice forms causing a frameshift (Table 1), 3 of which simultaneously decrease constitutive splicing by ≥ 10-fold (*WBP2NL* rs17002806; *GUSBP11* rs3747107; and *IFI44L* rs1333973). rs3747107 and rs17002806 are much more common in YRI populations in HapMap (rs3747107” G-allele is present in 64% in YRI but only 23% in CEU; rs17002806 A-allele was not identified in any CHB or CEU individuals), while the A-allele of rs1333973 is much more common in CHB (76%, compared to 31% and 35% in CEU and YRI populations, respectively). These common variants are likely to change the function of these proteins and may influence individual phenotypes. A somewhat comprehensive catalog of DNA polymorphisms with splicing effects - confined or with increased prevalence in specific ethnic or geographically-identifiable groups - could be derived from combining ValidSpliceMut with population-specific SNP databases. Aside from those phenotypes described earlier, genes implicated by GWAS or other analyses for specific disorders represent reasonable candidates for further detailed or replication studies aimed at identification of the risk alleles in these cohorts.

The extent to which SNP-related sequence variation accounts for the heterogeneity in mRNA transcript structures has been somewhat unappreciated, given the relatively high proportion of genes that exhibit tissue-specific alternative splicing (Wang *et al*. 2008; Pan *et al*. 2008; Baralle *et al*. 2017). This and our previous study (Shirley *et al*. 2019) raise questions regarding the degree to which apparent alternative splicing is the result of genomic polymorphism rather than splicing regulation alone. Because much of the information required for splice site recognition resides within neighboring introns, it would be prudent to consider contributions from intronic and exonic polymorphism that produce structural mRNA variation, since these changes might be associated with disease or predisposition.

Individual information corresponds to a continuous molecular phenotypic measure that is well suited to the analysis of contributions of multiple, incompletely penetrant SNPs in different genes, as typically seen in genetically complex diseases (Cooper *et al*. 2013). Our protocol identifies low or non-penetrant allele-specific alternative splicing events through bioinformatic analysis, and either q-RT-PCR, exon microarrays or RNAseq data analysis. Allele-specific splicing can also be determined by full-length alternative isoform analysis of RNA (or FLAIR [Workman *et al*. 2019]). Differentiated splice forms are associated with specific alleles in heterozygotes with exonic SNPs. However, combining genome-base information with FLAIR may enable identification of intronic SNPs influencing splicing and low abundance alternative splice forms, which might otherwise be missed by FLAIR.

Targeted splicing analysis generally reproduces the results of our multi-genome-wide surveys of sequence variations affecting mRNA splicing. As splicing mutations and their effects were often observed in multiple tumor types, the impact of these mutations may be pleiotropic. Some events were only detected in q-RT-PCR data and not by RNAseq (and vice versa), highlighting the complementarity of these techniques for splicing mutation analyses. Results of this study increase confidence that the publicly (https://ValidSpliceMut.cytognomix.com) and commercially (https://MutationForecaster.com) available resources for information-theory based variant analysis and validation can distinguish mutations contributing to aberrant molecular phenotypes from allele-specific alternative splicing.

## Supporting information

Supp. Methods and Results

## Acknowledgements

P.K. Rogan acknowledges support from The Natural Sciences and Engineering Research Council of Canada (NSERC) [371758-09 and RGPIN-2015-06290], Canadian Foundation for Innovation, Canada Research Chairs, and CytoGnomix.

## Author Contributions

EJM designed and performed all q-RT-PCR experiments, processed and analyzed publicly exon microarray data, and performed all formal analysis. BCS performed data curation and software development. PKR conceptualized the project and was the project administrator. EJM and PKR prepared the original draft of the manuscript, while EJM, BCS and PKR reviewed and edited the document.

## BioRxiv

An earlier version this article is available from bioRxiv: https://www.biorxiv.org/content/10.1101/549089v1 (Mucaki *et al*. 2019)

