## Supplementary material for "Expression changes confirm genomic variants predicted to result in allele-specific, alternative mRNA splicing": Supp. Figure 1

**Supplementary Figure 1 Overview:**

The following section describes SNPs discussed in the main document in greater detail. Each SNP has an associated figure below the detailed description that is made up of four sections: The exon microarray splicing index (SI) data organized into a boxplot which separates individual by the genotype of the SNP of interest (except for rs16994182); the UCSC genome browser image of the region of interest which displays the gene (and its various splice forms), the location of the SNP of interest and the placement of the probeset of the boxplot; and for most SNPs, an RNAseq image from an individual with the SNP, and a visual representation of the splice form tested is given (each box represents an exon, horizontal lines are introns, and two diagonal lines indicate the natural/cryptic sites used to form the junction tested). The thickness of the diagonal lines connecting two exons or one exon and a cryptic splice site denotes the fold change in abundance of each splice isoform detected for that individual based on q-RT-PCR experiments. The UCSC genome browser images have been modified to eliminate unrelated SNPs.

**Supplementary Figure 1.1 – rs2070573 (*C21orf2* [*CFAP410*]):** (A) The SNP rs2070573 (AvHet: 0.441 +/- 0.162) is a common polymorphism which alters the first nucleotide of the extended form of *C21orf2* exon 6 (mRNA splice forms AB209578 and BC031300); (B) The donor site is strengthened by the presence of the C allele (*R_i_* 0.4 to 4.0 bits; A>C) and its use extends the length of exon 6 by 360 nt. Primer design was complicated due to the SNP laying within the junction. Therefore, two sets of primers were designed to amplify the extended form for each possible allele. Though the extended form was detected in the A/A individual, the splice form was detected 3.7-8.5-fold and 17.4-22.6 fold greater in the A/C and C/C individuals, respectively. A second set of individuals tested in the same manner was also increased in abundance. The detected increase exceeded the theoretical change predicted by information analysis; (C) The exon microarray probeset (ID 3934488) which detects the extension clearly shows a step-wise increase in SI with individuals with the C-allele, which supports the bioinformatic and q-RT-PCR results. The splice form which uses the unaffected natural donor site (8.4 bits) is more abundant, ranging from 2.8 to 91-fold more depending on genotype. The assay failed to show a significant difference in the quantity of this splice form between the individuals tested. The internal reference varied in abundance between cell lines, but this did not appear to be genotype related; (D) This variant was present in the ValidSpliceMut database, which showed a significant increase in intron retention within the region of the extended exon [Box 1; example IGV image from RNAseq data of heterozygous TCGA BRCA patient, TCGA-BH-A0H0].

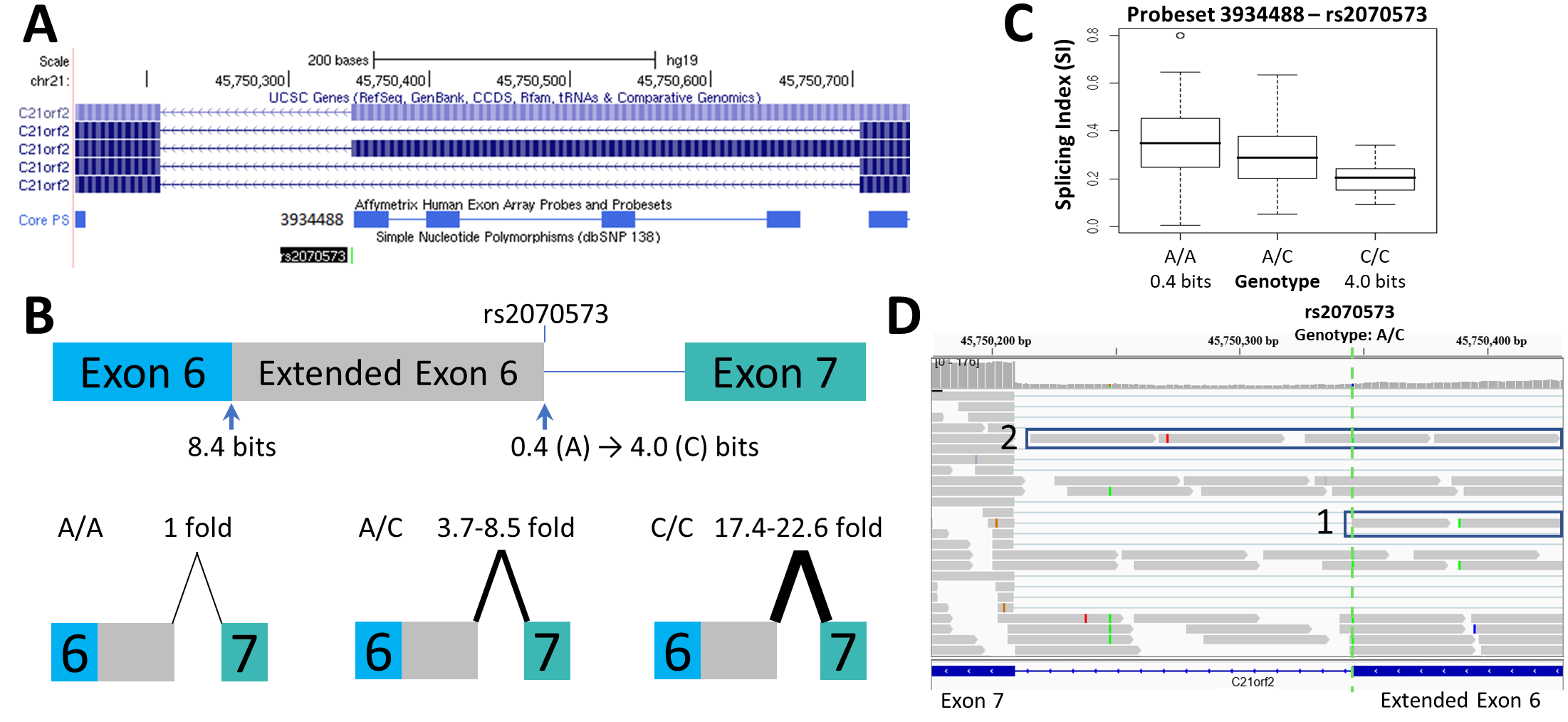

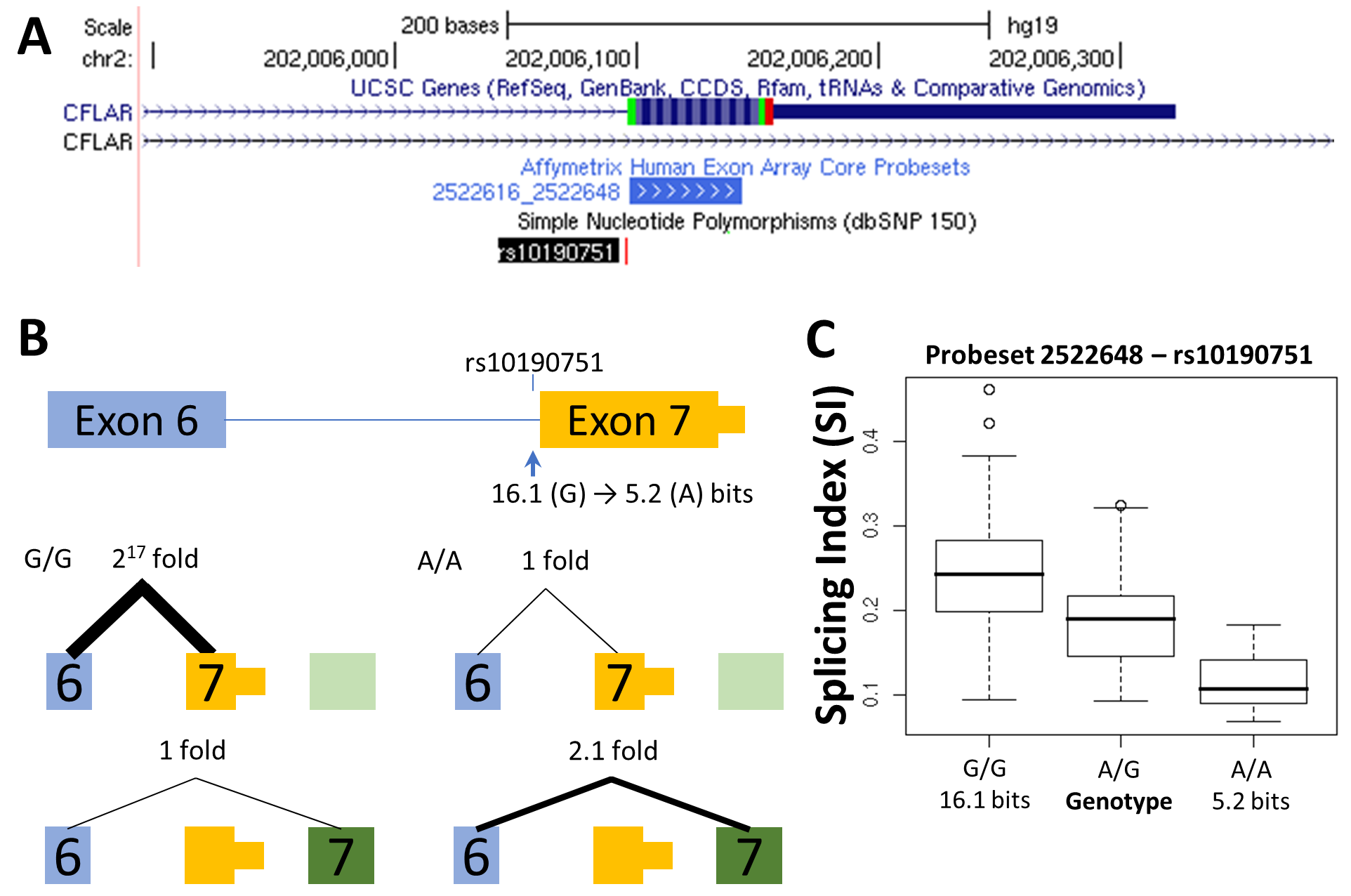
**Supplementary Figure 1.2 – rs10190751 (*CFLAR*): (**A) rs10190751 modulates the presence of the shorter c-FLIP(S) splice form (Ueffing *et al.* 2009) and serves as a positive control the prediction and detection of splicing effects by this SNP; (B) The G allele of rs10190751 activates the use of a 3’ terminal exon (A) by strengthening the acceptor site from 5.2 bits to 16.1 bits (A>G). Use of this exon was minimally detectable in the A/A cell line tested (~2^17^ less than G/G). While the A-allele acceptor is expected to be active (5.2 bits), site use was nearly undetectable. Presence of the longer splice form (c-FLIP (L)) was highest in the individual homozygous for the weak allele (A/A; 2.4 fold higher than G/G). The c-FLIP (R) splice form, which retains the entirety of intron 6, was not tested; (C) The exon microarray boxplot of the probeset which hybridizes to this exon (probeset ID 2522648) supports the q-RT-PCR result, reflecting the reduced inclusion of this exon in the presence of the A-allele; (D) The SNP was also present in the ValidSpliceMut web beacon, demonstrating a significant increase in intron retention of RNAseq reads within the region of the extended exon, which may be reflective of an increase in the c-FLIP (R) splice form. RNAseq from individuals with rs10190751 (IGV image from chronic lymphocytic leukemia [CLLE] ICGC patient DO6492 [heterozygote for SNP]) shows the presence of all 3 splice forms: c-FLIP(R) [Box 1], c-FLIP(S) [Box 2] and c-FLIP(L) [Box 3].

**Supplementary Figure 1.2 (cont.)**
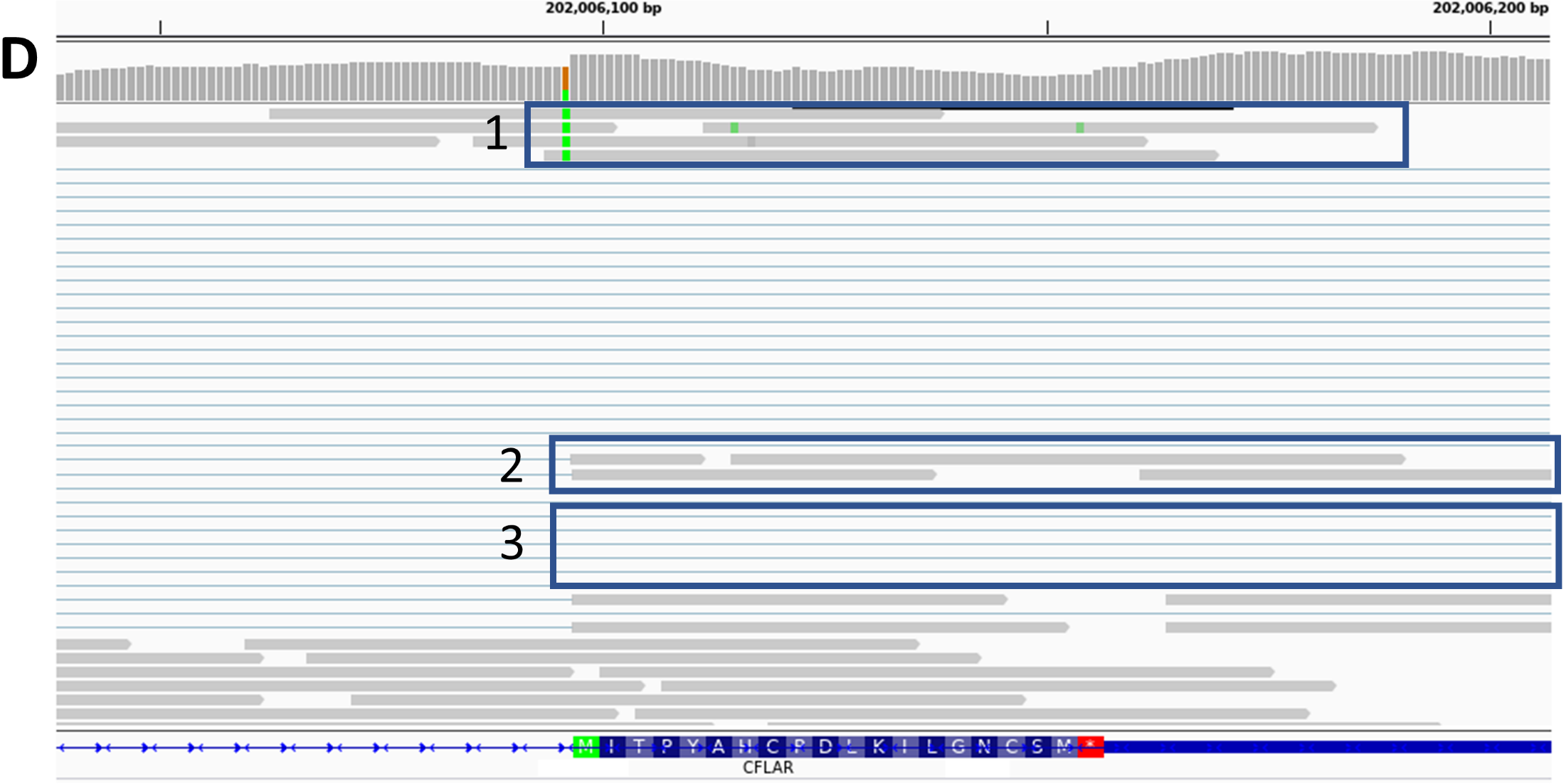

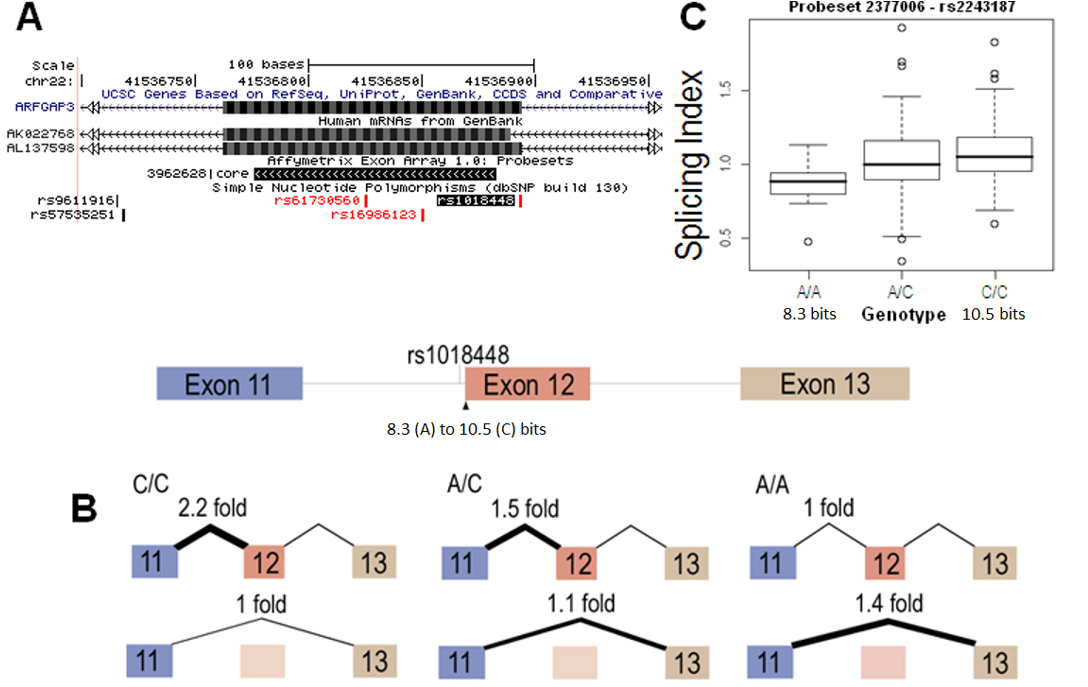
**Supplementary Figure 1.3 – rs1018448 (*ARFGAP3*):** rs1018448 strengthens the acceptor of exon 12 of *ARFGAP3* by 2.2 bits (*R_i_* 8.3 to 10.5 bits; A>C); (A) The UCSC genome browser (image from HG18 version of browser) displayed several mRNAs (AK022768, 10 others) which used an acceptor 5nt downstream of the natural acceptor site that was not recognized by information analysis. Primers designed for this region only cross-hybridized with the spliced product. Inspection of the sequence of these splice forms suggested this splice form might be a consequence of sequence alignment error; (B) The A/A individual tested was found to have 112-139% of the exon skipping of individuals carrying the C-allele, despite lower *ARFGAP3* mRNA levels. When normalizing against an internal reference gene, the A/A individual showed 153-257% of exon skipping compared to the A/C genotype and two C/C individuals tested; (C) An increase in the mean SI of the probeset in exon 12 for individuals with the C-allele suggested the possibility of exon skipping. Thus, the effect of rs1018448 on exon inclusion is reflected in the microarray boxplot. The presence of the C-allele appears to decrease exon skipping and increase *ARFGAP3* mRNA levels in the HapMap; (D) RNAseq data of individuals with this SNP (example from malignant lymphoma [MALY] patient DO27773 [ICGC]; which was heterozygous for SNP) show both normal splicing in the presence of the the mutation, limited intron retention [Box 1] and exon 12 skipping [Box 2].

**Supplementary Figure 1.3 (cont.)**

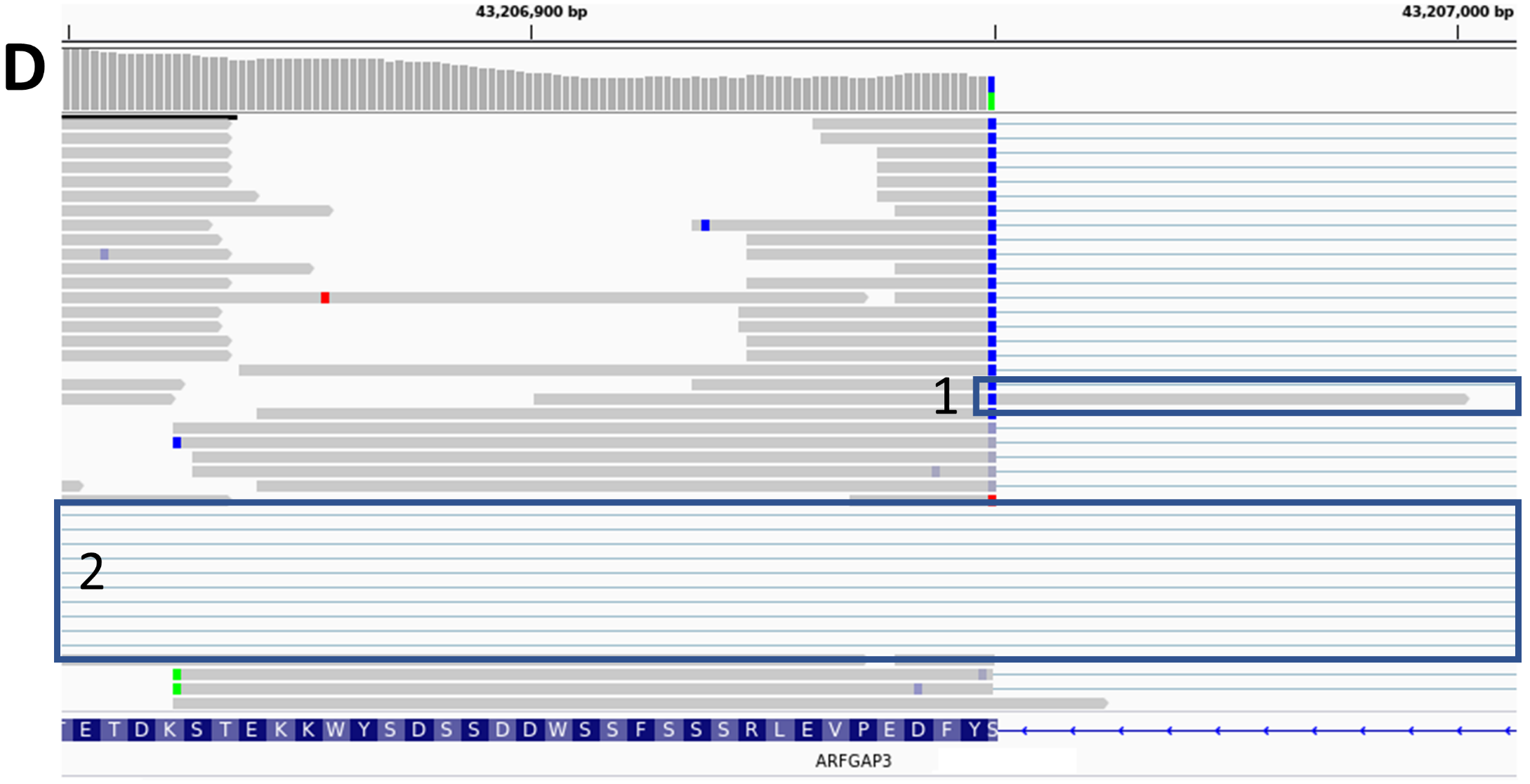

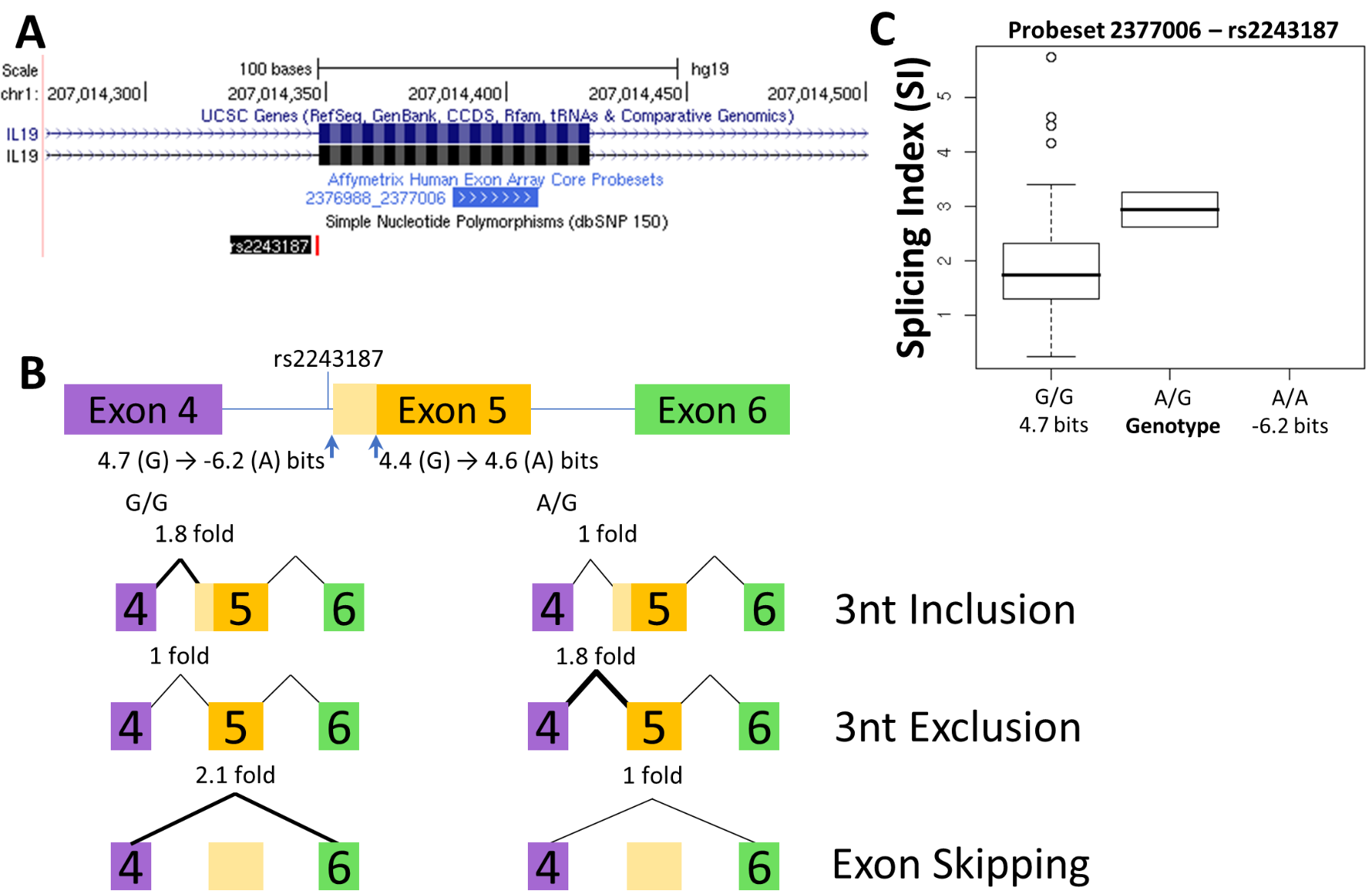
**Supplementary Figure 1.4 – rs2243187 (*IL19*):** (A) rs2243187 is an uncommon SNP within *IL19*. Homozygotes for the rare allele were not available in the HapMap or the Polymorphism Discovery Resource (Average Heterozygosity: 0.012 +/- 0.075); (B) The A-allele is predicted to abolish the acceptor site of exon 5 (*R_i_* = 4.7 to -6.2 bits; G>A), while a second cryptic site 3nt downstream is slightly strengthened (4.4 to 4.6 bits). Its use would lead to the deletion of a glutamine residue. Q-RT-PCR experiments detected 56.8% of the 3nt inclusion splice form in the heterozygote, which is the expected result if only one of two alleles are active. There seemed to be a slight preference for use of the upstream acceptor (168% more of the upstream acceptor isoform compared to the mRNA using the downstream acceptor in an individual with the G/G genotype by q-RT-PCR). The A/G genotyped individual expresses nearly twice the amount of (180%) the 3nt short form compared to the homozygote. There have been previous reports where two closely situated splice sites can disrupt wildtype splicing. This could explain why exon skipping events were less common in the heterozygote compared to the homozygote (205%; 7.5-14.7% of total *IL19*); (C) The exon microarray boxplot similarly shows a general trend of increased SI in heterozygotes (although there are some outlier homozygotes with higher SI values; probeset ID 2377006). The close proximity of the two acceptors seems to partially abrogate splicing in this region. Inactivation of one of the two splice forms eliminates this effect and reduces exon skipping. TCGA and ICGC individuals with the rs2243187 SNP did not express sufficient levels of *IL19* expression to perform this analysis.

**Supplementary Figure 1.5 – rs2835585 (*TTC3*):** The acceptor of *TTC3* exon 3 is weakened by rs2835585 (*R_i_* 7.6 to 5.4 bits; T>A). Two nearby cryptic acceptors are stronger than the natural site (*R_i_* 5.8 bit site 60nt upstream, 5.9 bit site 87nt upstream); (A) Evidence of their use or of exon skipping on the UCSC Genome Browser was not found. There was evidence of an intron retention splice form (IVS2) but q-RT-PCR did not detect a genotype-specific change, and neither cryptic site was amplified in any individual tested. It is clear that a strong splice site in close proximity to a natural splice site does not guarantee its use and reflects the requirement experimental evidence to confirm these predictions; (B) An increase in exon skipping was detected in individuals with the A-allele. Heterozygotes showed 2-6x additional exon skipping and A/A homozygotes showed 3-9x compared to homozygous T/T individuals. Regardless of genotype, exon skipping is a rare event (Table 2). This increased abundance of the rare transcript may not appreciably impact the overall levels of the major, constitutive splice form; (C) The microarray boxplot did not follow a stepwise pattern corresponding to genotype and did not show a decrease in homozygous weak individuals. This is understandable as total exon inclusion splice form was not significantly changed in abundance. The internal reference showed variable expression, which was independent from the levels exhibited by those carrying the rs2835585 A/A genotype; (D) The ValidSpliceMut web beacon flagged rs2835585, resulting from increased intron retention rather than increased exon skipping. This may support the use of either of the previously described intronic cryptic sites or it may suggest total retention of intron 2; RNAseq data of individuals with this SNP (representative IGV image from heterozygous ICGC MALY tumour DO27803) exhibited very limited exon skipping (reads shown in Box 1 do not indicate exon 3 skipping, rather they are reads from an alternate *TTC3* splice form [e.g. NM_001330681]), but as suggested by ValidSpliceMut, there is evidence of intron retention [Box 2].

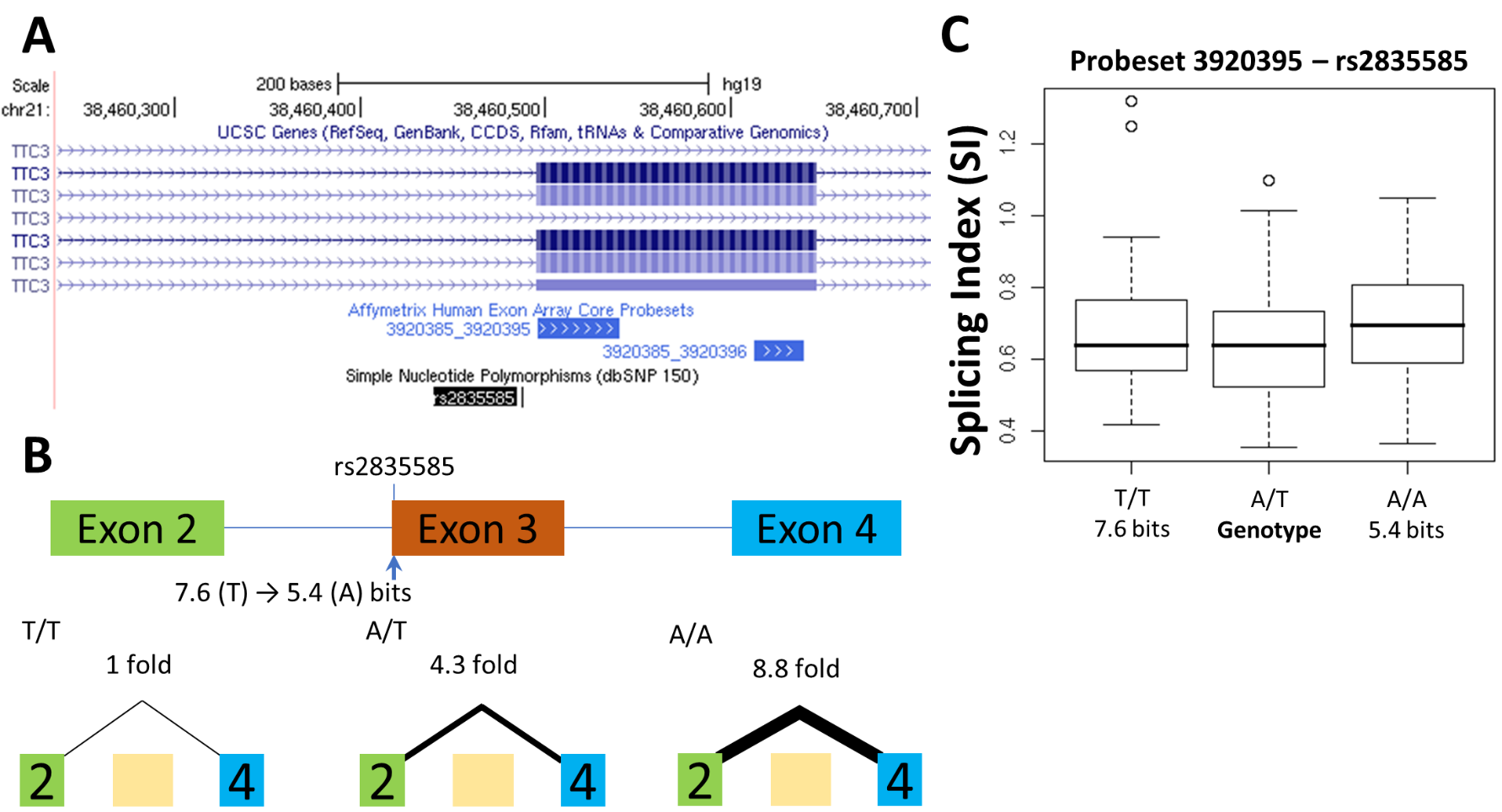

**Supplementary Figure 1.5 (cont.)**

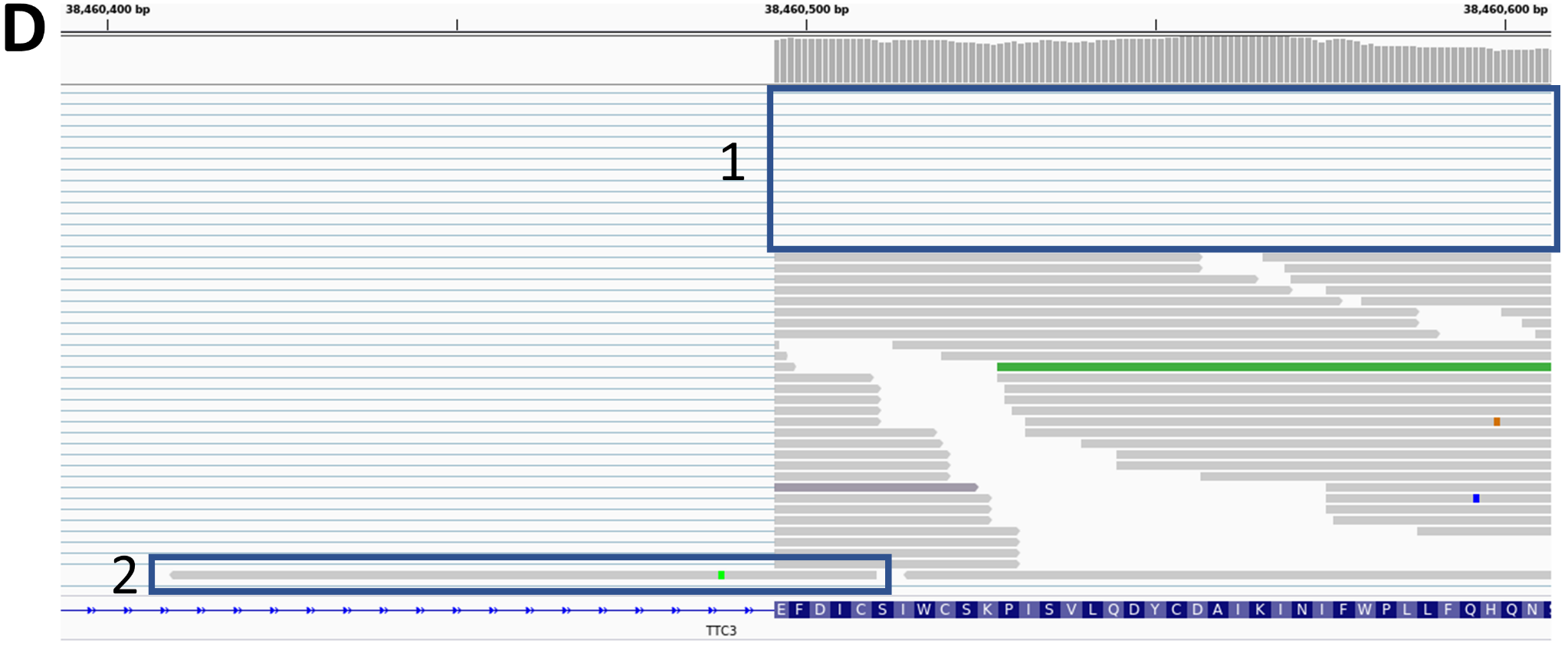

**Supplementary Figure 1.6 – rs1805377 (*XRCC4*):** The effect on splicing by rs1805377 (*XRCC4*) has been previously predicted (Nalla and Rogan, 2005). This SNP has been linked to bladder (Wu *et al*. 2006) and renal cell carcinomas (Margulis *et al.* 2008). The SNP is less common in European and most common in Asian populations (AvHet 0.492 +/- 0.063). It reduces the strength of the upstream acceptor of the exon 8 of *XRCC4* from 11.5 bits to 0.6 bits (G>A), while a second site 6nt downstream is slightly strengthened (*R_i_*: 11.4 -> 11.8 bits); (A) Transcripts which use either acceptor site has been previously described (mRNAs NM_022406, NM_003401); (B) Despite their similar strengths, results suggest a near exclusive preference of the upstream acceptor (G form) over the downstream acceptor (A form) when the G-allele is present. Additionally, dual-labelled probe experiment designed to discriminate between the G and A-forms completely discriminated the two splice forms between the two homozygotes (not shown). While it is unclear which assay is more accurate (probe-based assays avoid interference from non-specific amplification), the A-allele of rs1805377 causes a preference for the 6nt deletion splice form, which may explain its link to the aforementioned cancer types; (C) The 6nt deletion cannot be detected by the exon microarray as the upstream probeset (ID 2818500) does not overlap the variable region; (D) The SNP was flagged by ValidSpliceMut for intron retention [Box 1], rather than cryptic site activation, although this is clearly observed in available RNAseq data [Box 2; example IGV image from RNAseq of heterozygous ICGC MALY patient DO27779].

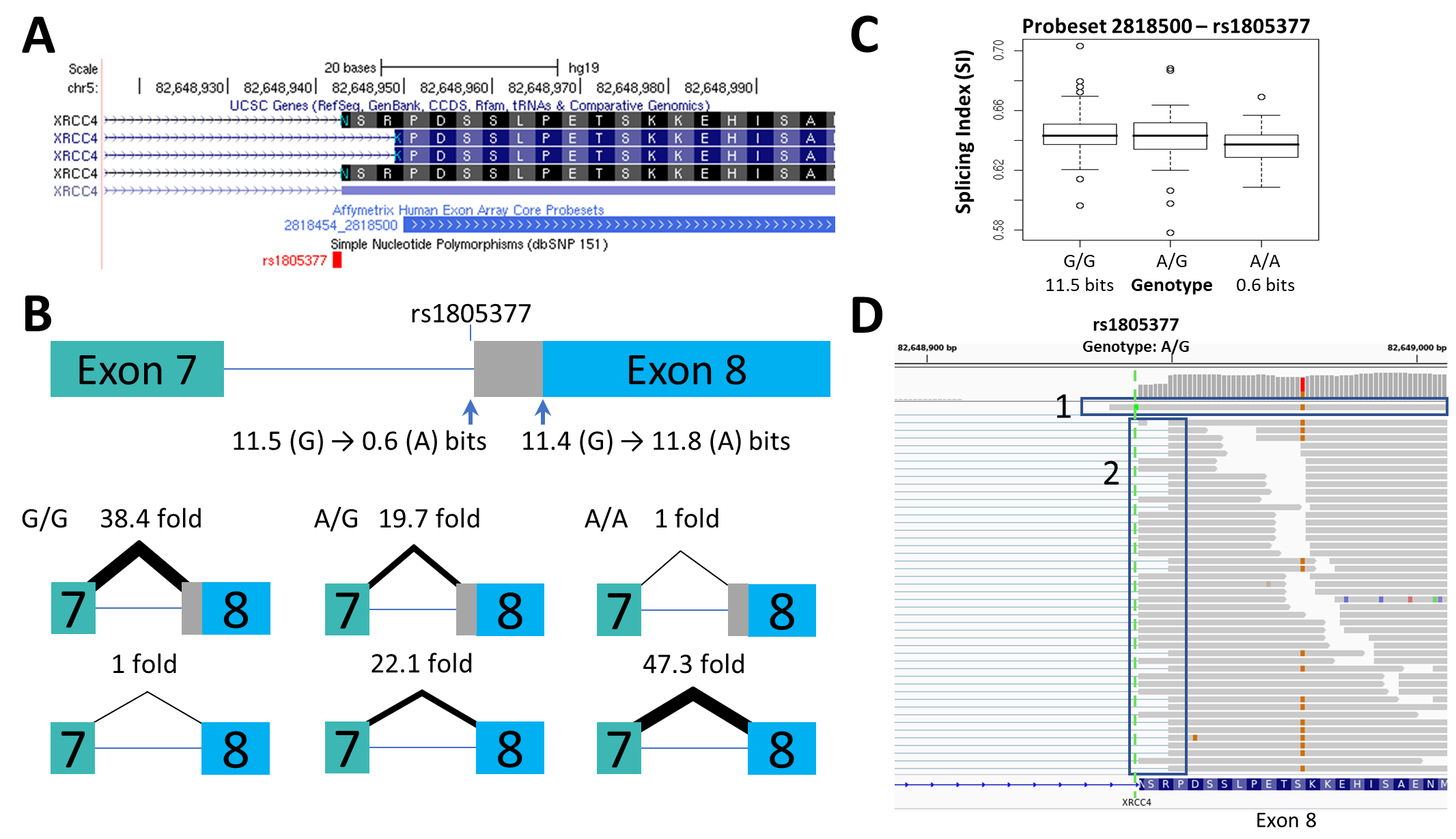

**Supplementary Figure 1.7 – rs743920 (*EMID1*): (**A) Exon 4 of the gene *EMID1* also contains a pair of neighboring acceptor splice sites separated by 6nt. Use of both sites has been reported previously (GenBank accession CU012965 uses the upstream site, AJ416090 uses the downstream); (B) The SNP strengthens the downstream acceptor site from 6.0 to 7.9 bits (C>G; 3.6-fold increase), while the weaker upstream acceptor (2.4 bits) is unaffected. Use of the downstream acceptor would result in a 6nt deletion of two amino acids from the coding region. Q-RT-PCR experiments clearly shows that the G-allele increases the use of the downstream site as the C/G and G/G individuals tested showed a 230% and 580% increase compared to the C/C individual, respectively. The splice form with a 6nt inclusion was 7.9-fold more abundant than the splice form lacking this sequence in an individual homozygous for the weaker allele (C/C; 1.8 fold more abundant in G/G individual). Despite the differences in strength, the splice form which uses the upstream 2.4 bit site generated the more abundant isoform. It is notable that other SNPs within both *IL19* and *XRCC4* that modulate the strengths of multiple acceptor sites showed preference for constitutional splicing using the upstream site. This pattern is consistent in *EMID1,* despite the upstream acceptor being significantly (17-64 fold) weaker. This observation is congruent with a processive mechanism of that recognizes functional acceptor splice sites; (C) This deletion does not overlap any exon microarray probeset, and therefore it cannot distinguish between the two splice isoforms; (D) RNAseq from patients with the SNPs (example from heterozygous ICGC MALY patient DO27769) showed partial use of the strengthened cryptic site in the presence of the SNP [Box 1] and an exon 4 skipping splicing event [Box 2] which was not originally considered as a possible outcome. It is possible that the alternative splice forms are influenced by rs743920, despite the natural site remaining intact, as it does create an exonic 2.8 bit hnRNP A1 site adjacent to the novel splice acceptor (not shown).

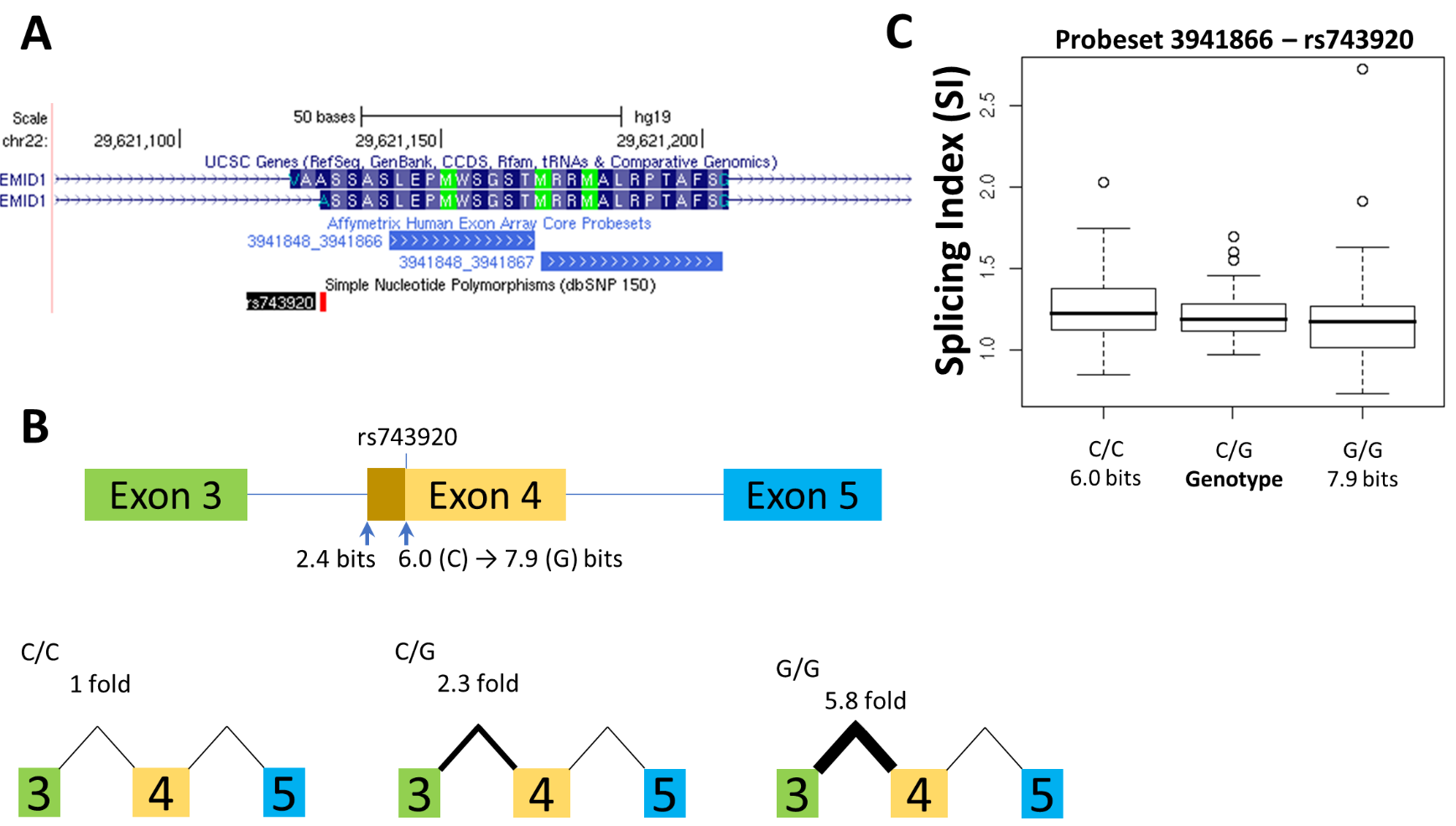

**Supplementary Figure 1.7 (cont.)**

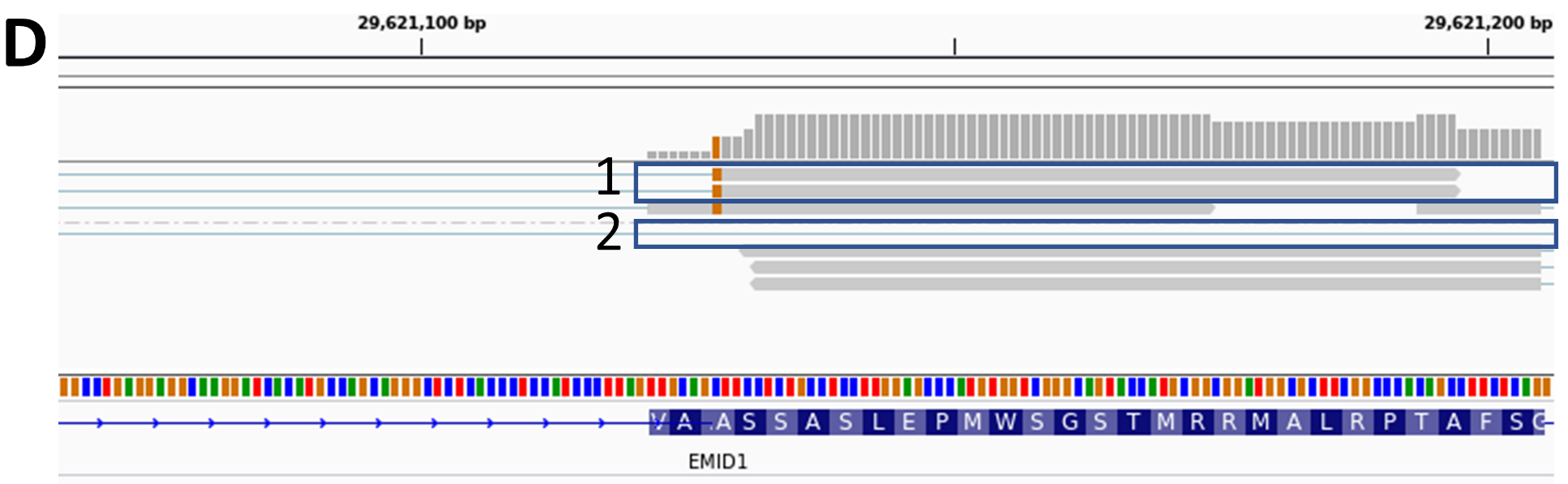

**Supplementary Figure 1.8 – rs3747107 (*GUSBP11*):** (A) The C>G variant of rs3747107 (AvHet 0.478 +/- 0.102) inactivates the natural acceptor site of a 3' terminal exon of *GUSBP11* (formerly *Fλ8;* GenBank mRNA accession BX538181; 4.7 to -7.0 bits). Cryptic acceptor sites 2nt downstream (*R_i_* -3.9 to 4.9 bits) were strengthened and both 114nt (6.5 bits) and 118nt (8.6 bits) upstream were also predicted. (B) Q-RT-PCR experiments demonstrated overall gene expression to vary slightly between individuals with different genotypes (C/G 134% of C/C amplification). The expression level of the homozygous G allele associated with the inactivated site was 1% of the homozygous normal allele. Use of the upstream cryptic sites was increased in the presence of the inactiving G-allele (Table 1), though they were less abundant relative to the internal reference (Table 2). *GUSBP11* alternatively splices to a downstream exon (Example GenBank accession BC063045, and 4 others). Primers were designed to detect changes in the use of this downstream exon. Expression levels of this exon increase in the presence of the G allele (116-133% in homozygous individuals); (C) Probeset 3954813 was expected to detect increased intron retention due to cryptic site use. While SI does increase, the change is slight and not statistically significant. Another mRNA (Accession AK024374) may also be detected by the same probeset, which could skew these results. Furthermore, a probeset which detects the alternative downstream exon was significant differ in mean SI values (not shown); (D) Interestingly, RNAseq data of patients with the SNP (example from heterozygous ICGC MALY patient DO27847) exhibit reads that demonstrate the cryptic site created by the SNP 2nt downstream is activated (Box 1). Additionally, reads support the use of the pre-existing cryptic site 118nt upstream of the affected acceptor site (Box 2).

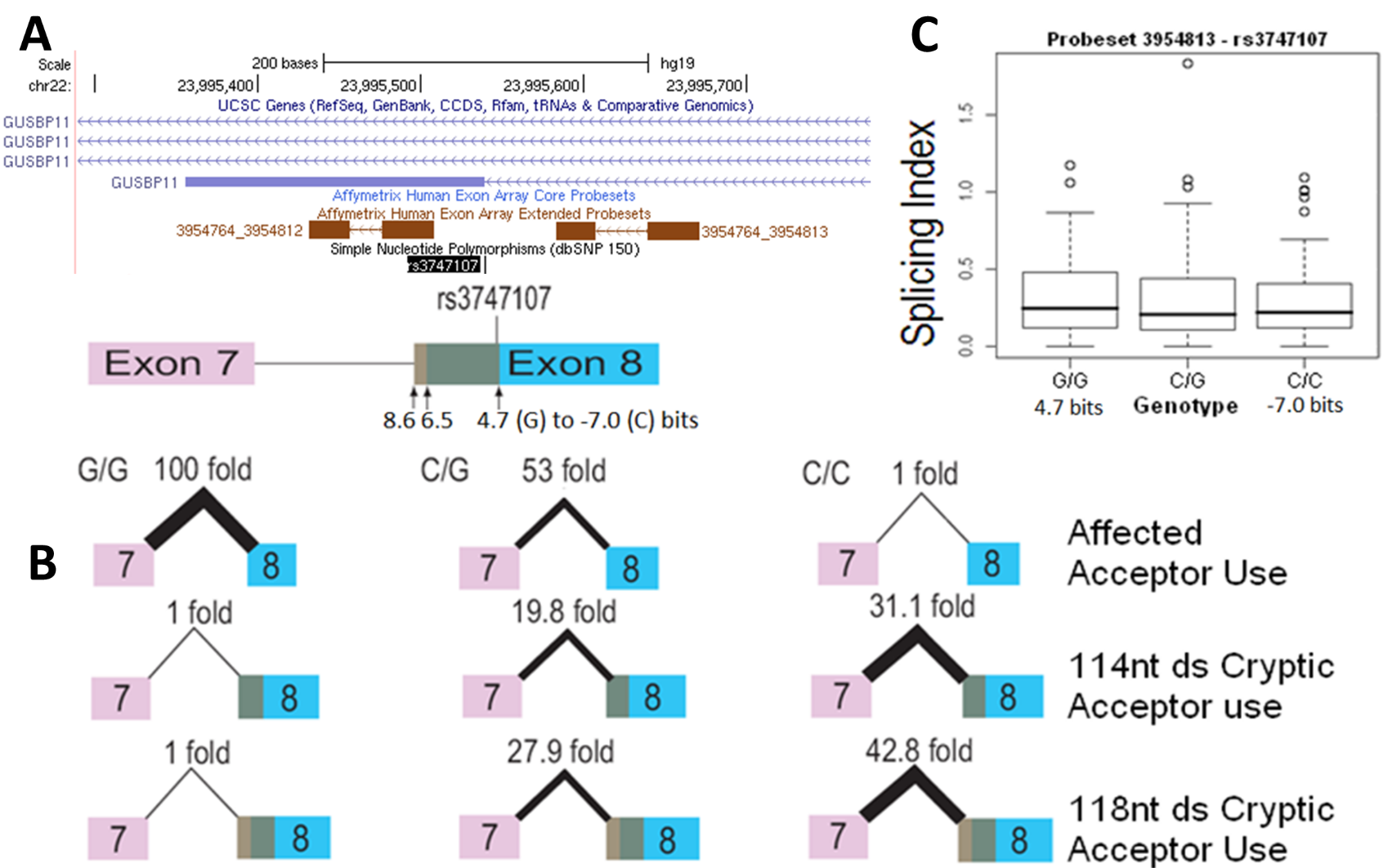

**Supplementary Figure 1.8 (cont.)**

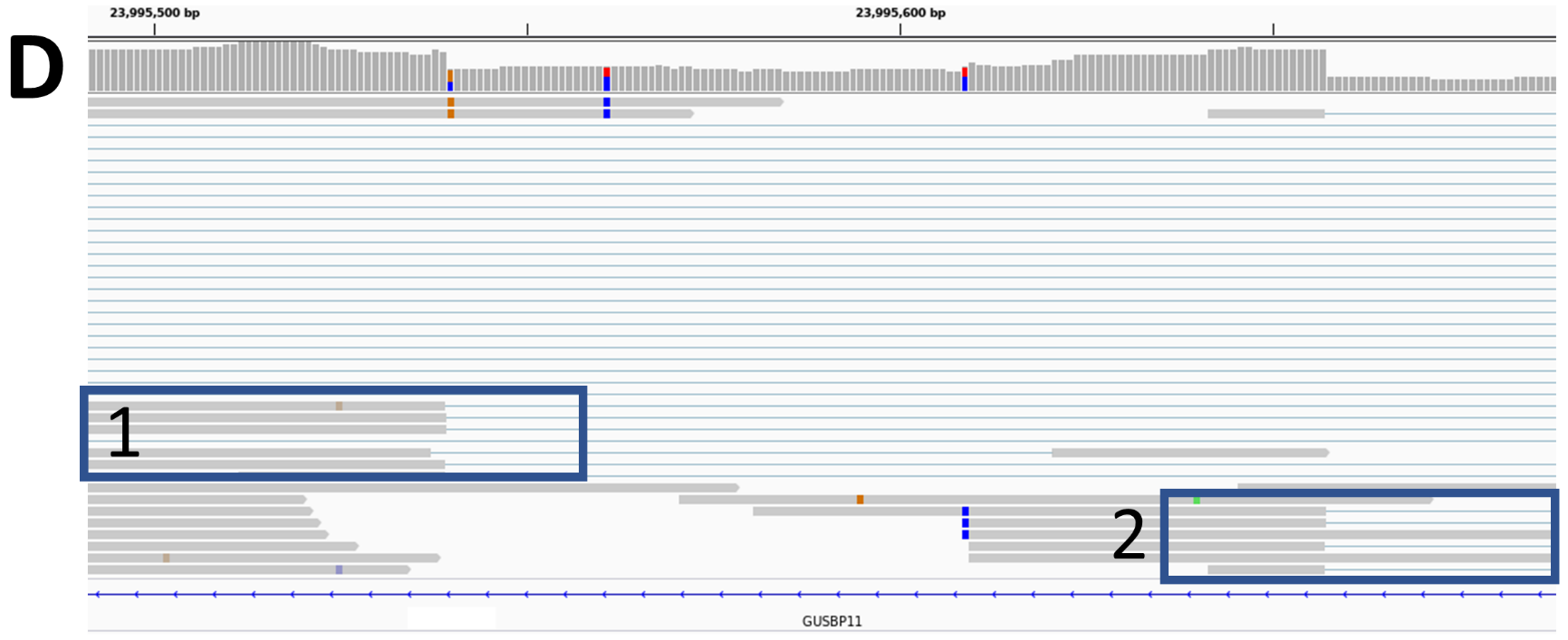

**Supplementary Figure 1.9 – rs2266988 (*PRAME*):** (A) rs2266988 weakens the natural donor site of exon 3 of *PRAME* by 1.6 bits or 3-fold (*R_i_* 7.8 to 6.2 bits; G>A). A different *PRAME* SNP rs2072049 affects the acceptor of exon 6, reducing its strength by 1.1 bits (8.2 to 7.1 bits; G>T). Detection of splicing effects of rs2266988 could be confounded by the rs2072049 genotype, as the latter SNP appears to affect total RNA levels (see Supplementary Figure 1.14). Therefore, only individuals homozygous common (G/G) for rs2072049 were considered in the evaluate rs2266988; (B) Low levels of skipping of exon 3 were found. Despite high standard errors, the increase in skipping detected in carriers of the weaker A-allele was statistically significant (460% and 876% greater in A/G and A/A compared to G/G, respectively); (C) Changes in splicing of exon 3 in the microarray data (probeset 3954440) were not detected, which was not unexpected, as skipping is significantly less common than constitutive splicing (see Table 2); (D) Skipping of exon 3 was observed in the RNAseq of individuals carrying the SNP (example IGV image from heterozygous ICGC MALY patient DO52652; Box 1]. Alternate splicing of the acceptor site of exon 3 is highlighted in Box 2, but this does not appear to be related to the rs2266988 genotype.

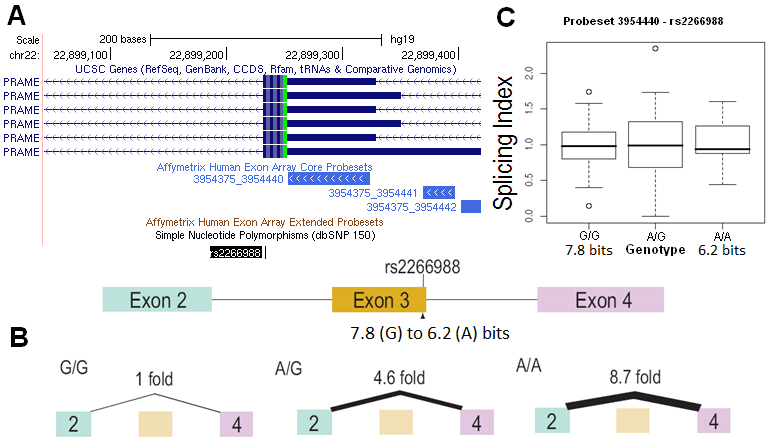

**Supplementary Figure 1.9 (cont.)**

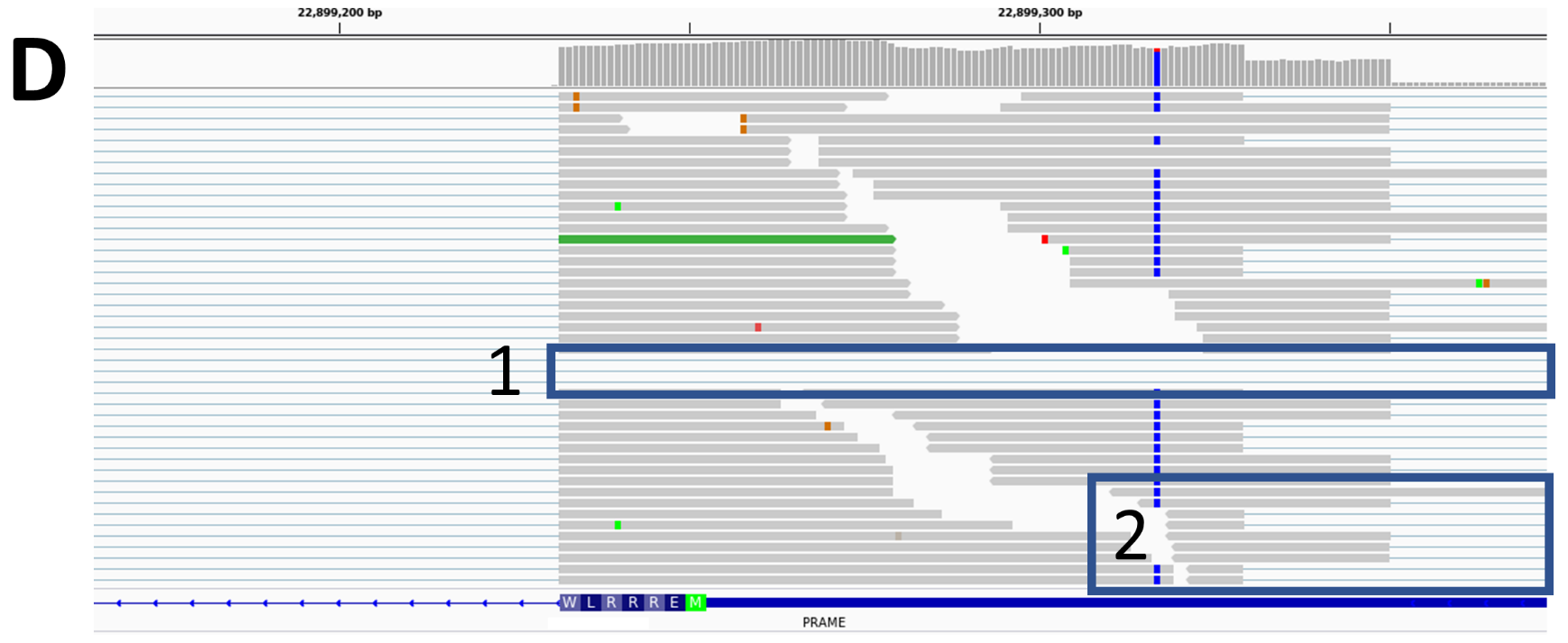

**Supplementary Figure 1.10 – rs1893592 (*UBASH3A*):** The natural donor site of *UBASH3A* exon 10 is weakened by 23-fold by rs1893592 (*R_i_* 8.7 to 4.2 bits; A>C). This SNP also strengthens a weak preexisting SRp40 binding site (*R_i_* 0.4 to 5.4 bits; A>C), which may enhance recognition of the 4.2 bit natural site. A 6.4 bit cryptic donor site is present 29nt downstream of the affected natural donor; (A) GenBank Accession BC028138 shows that a 5.2 bit donor 555nt downstream of the natural site can also be used (extending the length of exon 10 to 678nt), and activation of this site may occur in individuals carrying the 4.2 bit natural site; (B) The internal reference amplicon showed highly variable C_t_ values between individuals; this variability does not seem to be SNP-related (the heterozygote 8-fold greater amplification of expression compared to homozygotes). Expression in the two homozygous cell lines differed by 38% and upon adjusting for this source of variability, the affected splice junction was used 2.2-fold less in the C/C homozygotes (3-fold unadjusted); (C) Boxplots show a marginal increase of the extended 678nt exon splice form in C/C homozygotes. Exon skipping was evaluated, but was not detected; (D) This variant was flagged in ValidSpliceMut due to an increase in intron retention in 9 patients. Three of these patients also exhibited significant exon skipping, which had not been detected by q-RT-PCR. RNAseq data of individuals with the SNP (examples from heterozygous ICGC MALY patients DO27773 [top] and DO52673 [bottom]; D) show inclusion of intron 10 sequences [Box 1], use of the extended exon 10 splice form 555nt downstream [Box 1], exon 10 skipping [Box 2], and use of the cryptic site 29nt downstream of the affected donor site [Box 3].

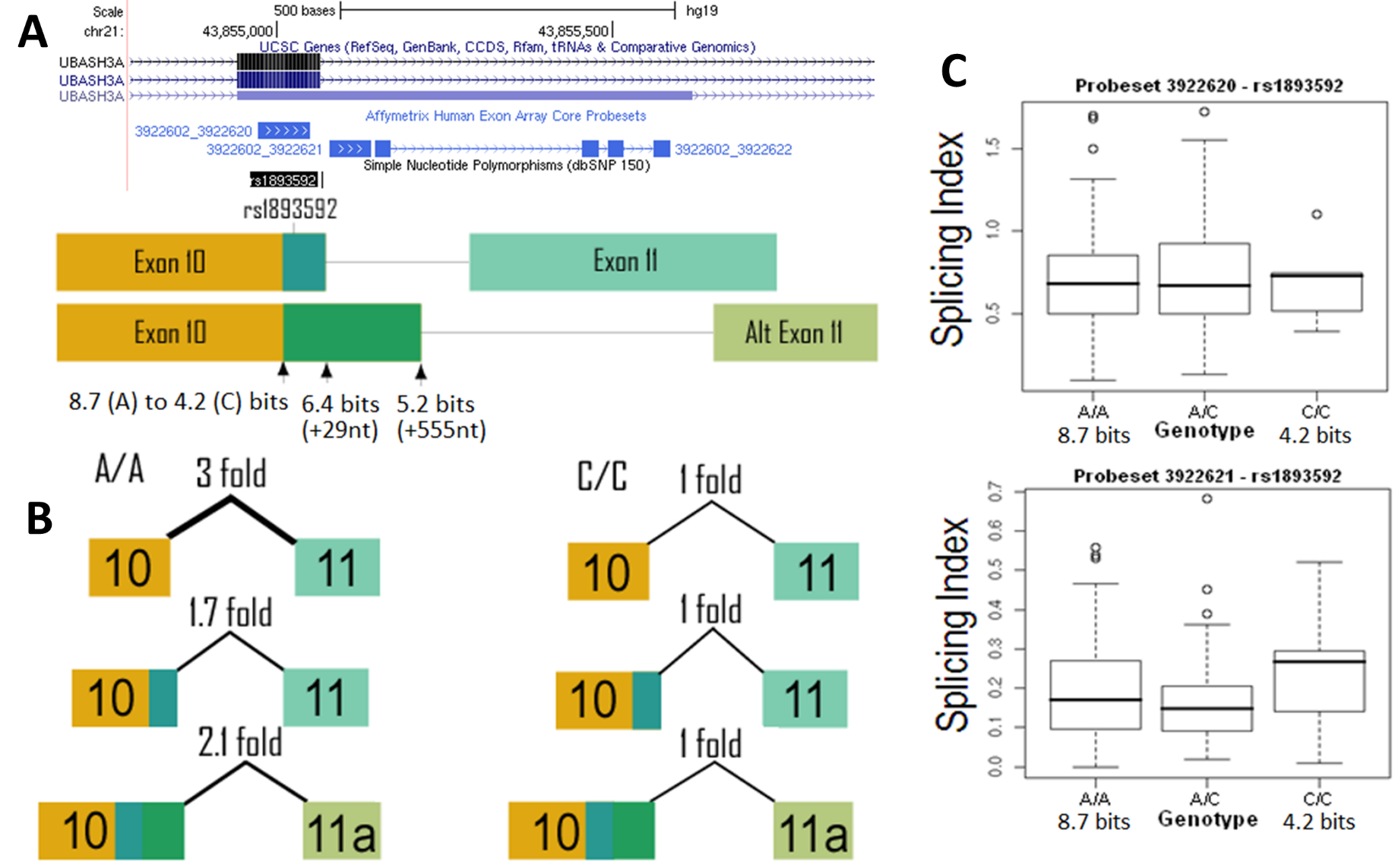

**Supplementary Figure 1.10 (cont.)**

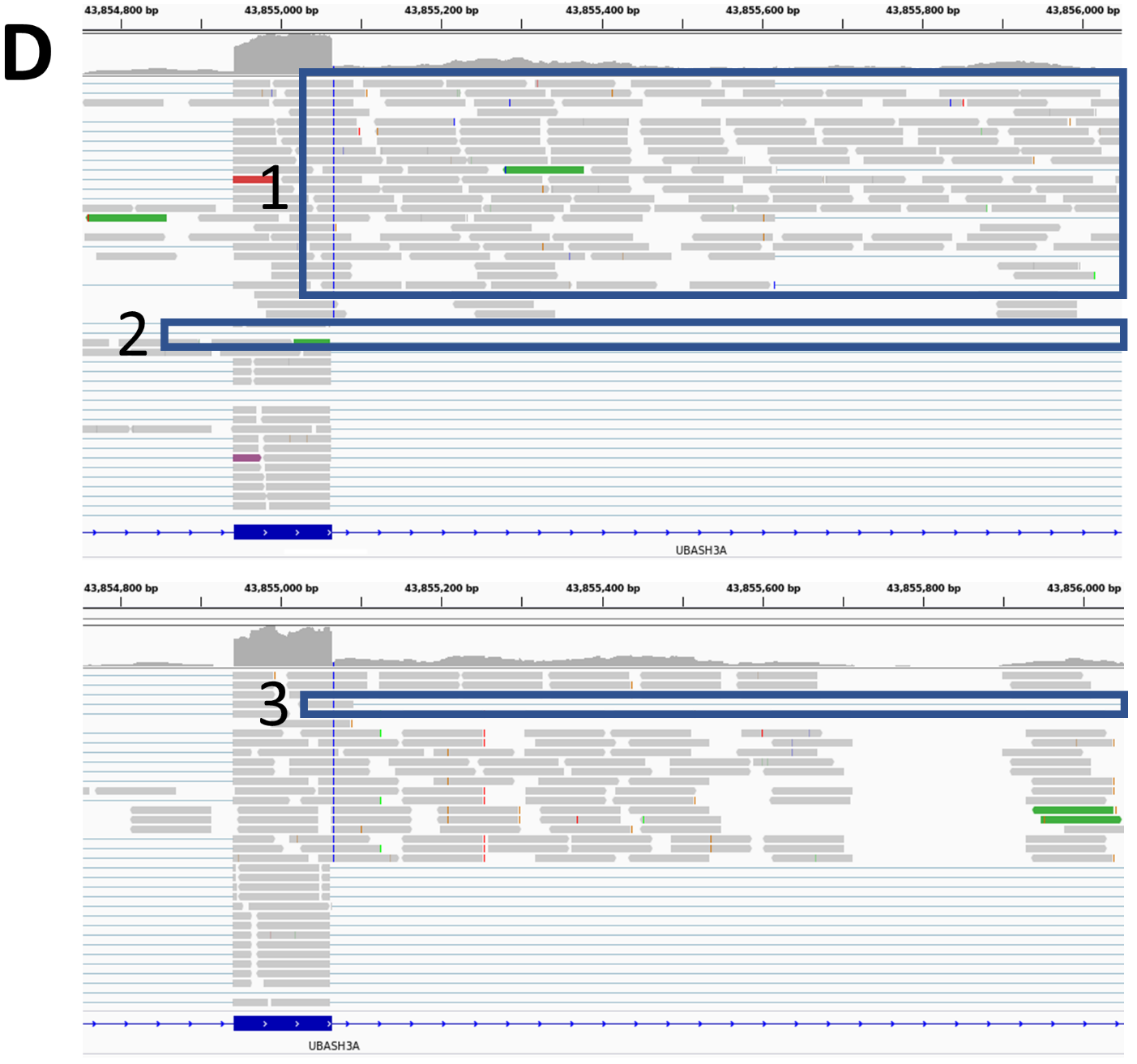

**Supplementary Figure 1.11 – rs1333973 (*IFI44L*):** (A) rs1333973 is a common SNP (AvHet - 0.500 ± 0.014) that weakens the natural donor site of exon 2 of *IFI44L* by 4.5 bits (*R_i_* 9.1 to 4.6 bits; A>T). Exon 2 is a large 488nt exon containing both coding and non-coding regions; (B) Q-RT-PCR experiments found a significant (15.6 fold) increase in exon skipping in T/T homozygous cell lines. Normal levels of splicing of exon 2 are reduced in the T/T homozygote by 15.4-fold (to 6.6%). The internal reference of *IFI44L* was also ~4-fold more abundant in the A/A homozygote with the stronger splice site. Assuming this difference is genotype-related, this change in overall abundance of *IFI44L* could influence interpretation of this analysis. Adjusting for this discrepancy, the difference in normal splicing between the homozygous genotypes would not be as pronounced, however but the difference in the extent of exon skipping would increase considerably in the T/T genotype; (C) The boxplot of the probeset which overlaps the affected exon (probeset ID 2343481) shows higher SI values in HapMap individuals carrying the stronger A allele; (D) RNAseq data of patients with the SNP (example is homozygous ICGC CLLE patient DO6354) reveals reads indicating total intron 2 inclusion [Box 1], the unanticipated use of a cryptic donor 375nt upstream of the affected donor [2.4 bits; Box 2], and skipping of exon 2 [Box 3].

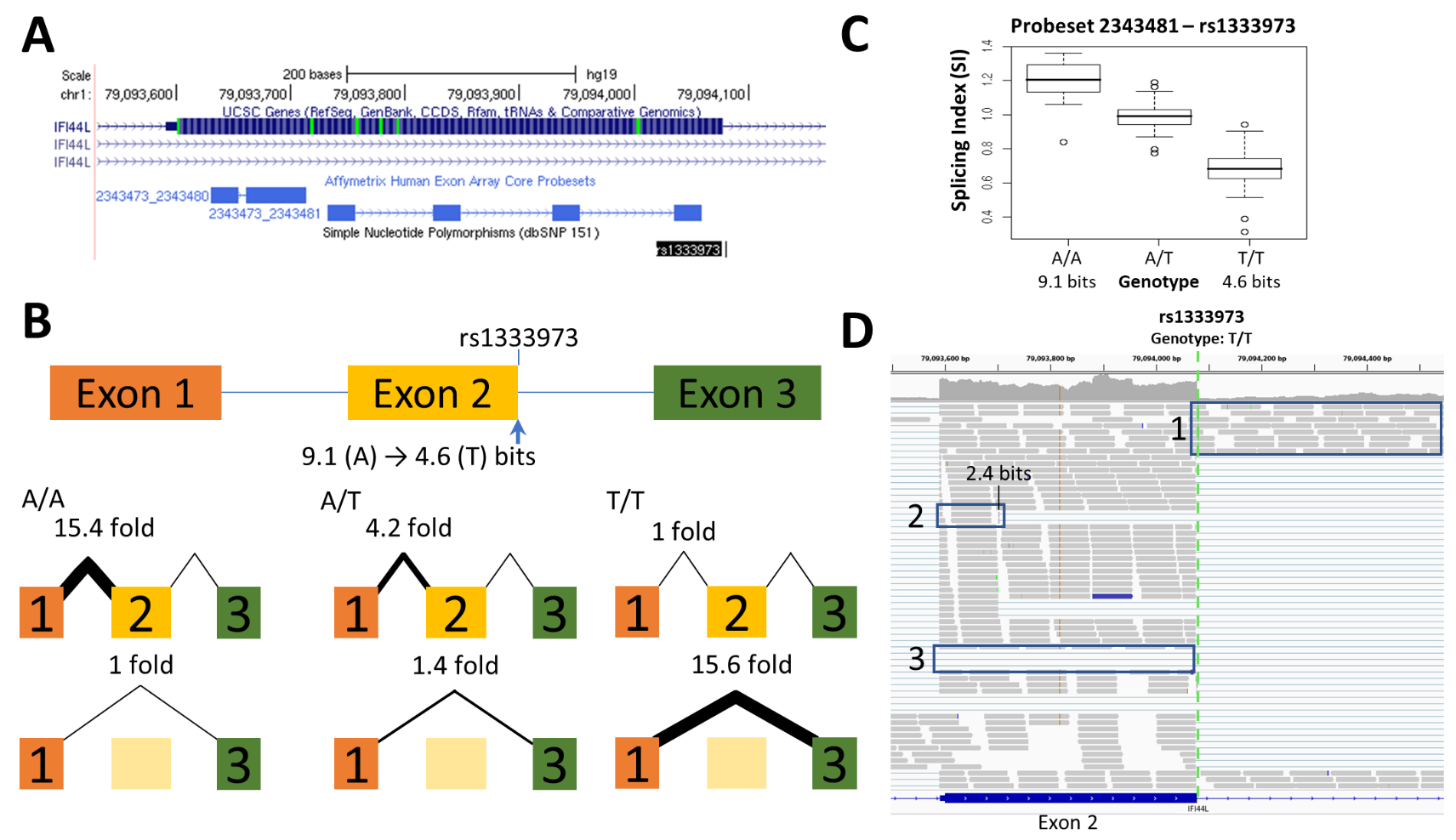

**Supplementary Figure 1.12 – rs13076750 (*LPP*):** (A) *LPP* codes is expressed at low levels in lymphoblastoid cell lines. The acceptor site of a rare non-coding exon within the first intron is abolished by rs13076750 (*R_i_* 9.3 to -1.6 bits; G>A). An alternate acceptor site for this exon is present 7nt downstream from the affected site (*R_i_*=2.0 bits); (B) Q-RT-PCR experiments confirmed that the affected natural site was inactivated by the A allele as it was not detectable in the individual homozygous for the weak allele (A/A), and the expression detected in the heterozygote reflected the expression of the single functional allele (56.3% of G/G individual). The 2.0 bit cryptic acceptor site is activated less frequently than the naturally spliced isoform, but is nearly 16 times more abundant in both individuals carrying the A allele. Exon skipping was also found to be increased by 10.3 and 26.6-fold in individuals heterozygous and the homozygous, respectively, for the weaker site (A allele); (C) The boxplot for this exon (probe ID 2657307) did not show a significant, consistent increase in individuals carrying G-allele. The SNPs effect may be masked by the increase in cryptic site use, as the probeset would not be able to distinguish between the two exon “1a” splice forms; (D) This variant flagged in the ValidSpliceMut web beacon for significantly increased abundance of intronic reads (relative to controls). RNAseq of individuals with the SNP (example shows CLLE patient DO6352 [ICGC]) exhibit reads indicating both use of the cryptic site 7nt downstream of the affected acceptor [Box 1], as well as total exon 1a skipping [Box 2].

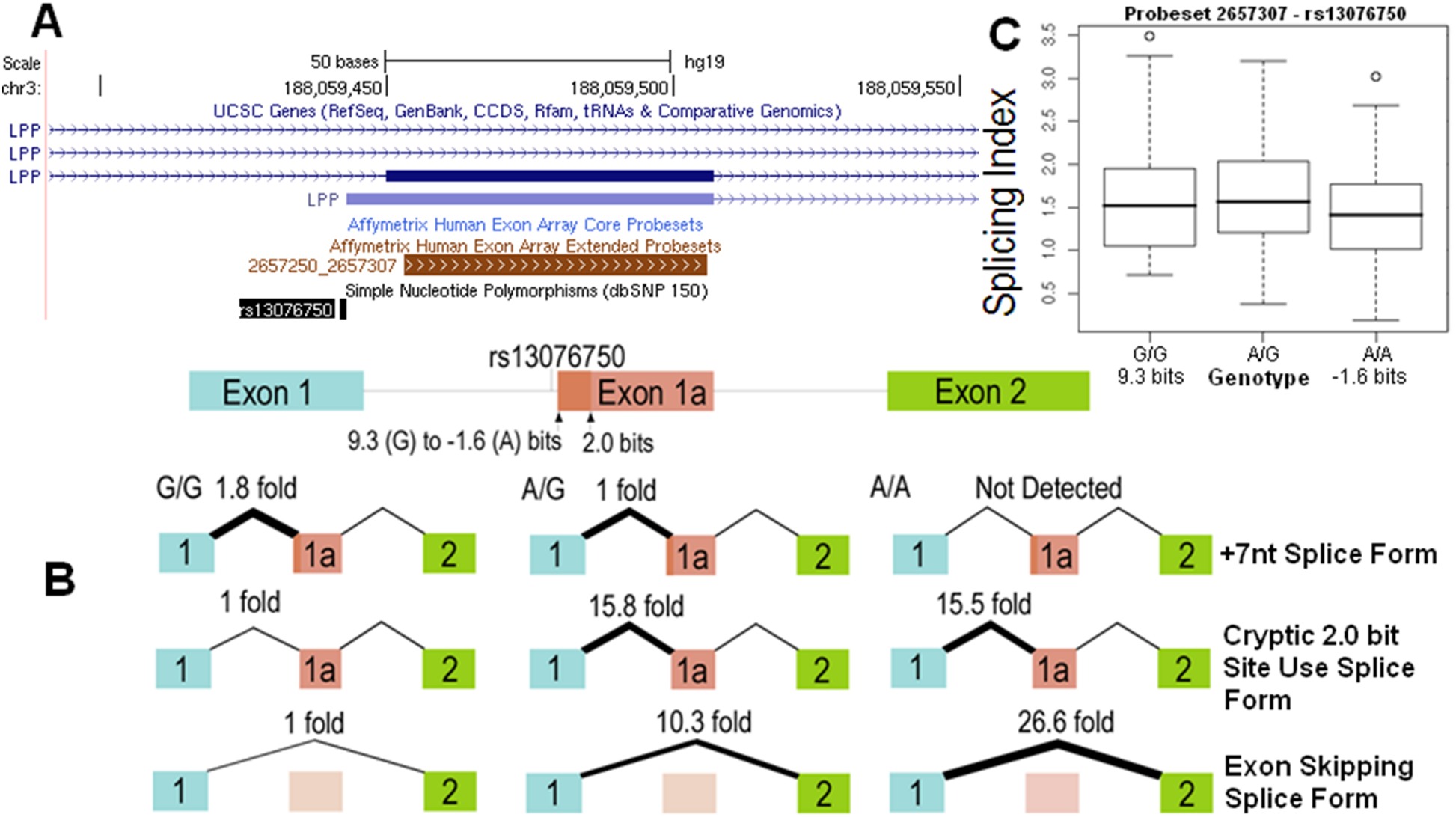

**Supplementary Figure 1.12 (cont.)**

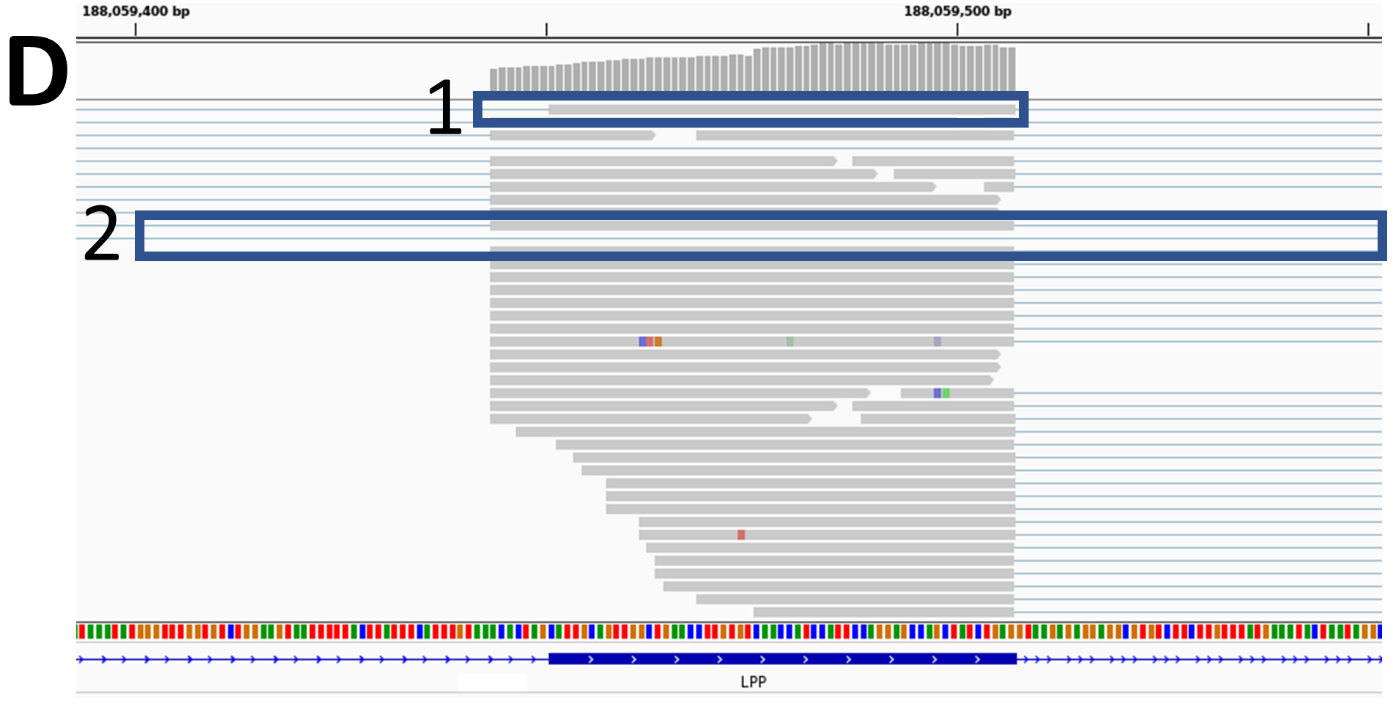

**Supplementary Figure 1.13 – rs17002806 (*WBP2NL*):** (A) rs17002806 (AvHet 0.062 +/- 0.165) weakens the donor splice site of exon 6 of *WBP2NL* from 9.2 to 6.0 bits (G>T); (B) Compared to G/G genotype, splicing from exons 6 to 7 dropped to 46.1% and 4.2%, respectively, for the G/T and T/T genotypes. The *WBP2NL* internal reference of the T/T and G/T individuals normalized to G/G individuals exhibited 67.5% and 95% expression, respectively. It is currently unclear if this difference is genotype related, as the internal reference will also amplify other splice isoforms of *WBP2NL*. A 5.8 bit site 25nt downstream was used with near equal abundance in the G/T and T/T individuals but was not detected in those with G/G genotype. This cryptic splice form is present at low levels relative to total *WBP2NL* levels (0.8 - 2.3% of internal reference). A second cryptic donor site 67nt downstream (5.7 bits) of the natural site was not detected; (C) Probeset 3947280 spans a region extending beyond the affected natural donor. The boxplot does not show a stepwise change in average expression for the three genotypes, but a general trend of increased intron retention in the presence of the T allele is evident, which could be consistent with the use of other downstream cryptic donor sites. Primers developed to detect the exon 5-6 junction by q-RT-PCR did not show significant differences between individuals, in contrast with those that amplified the junction between exons 6 and 7. Exon skipping was not detected. Therefore, this SNP impacts splicing of the affected exon 6 donor, but does not impact retention of exon 6; (D) RNAseq data of patients carrying the T allele with expressing sufficient levels of WBP2NL mRNA was rare. In patients that did meet this criteria, reads using the affected donor (or the 25nt intron 6 inclusion splice form) w not observed (example: RNAseq results from TCGA Thyroid Carcinoma [THCA] patient TCGA-DO-A1JZ).

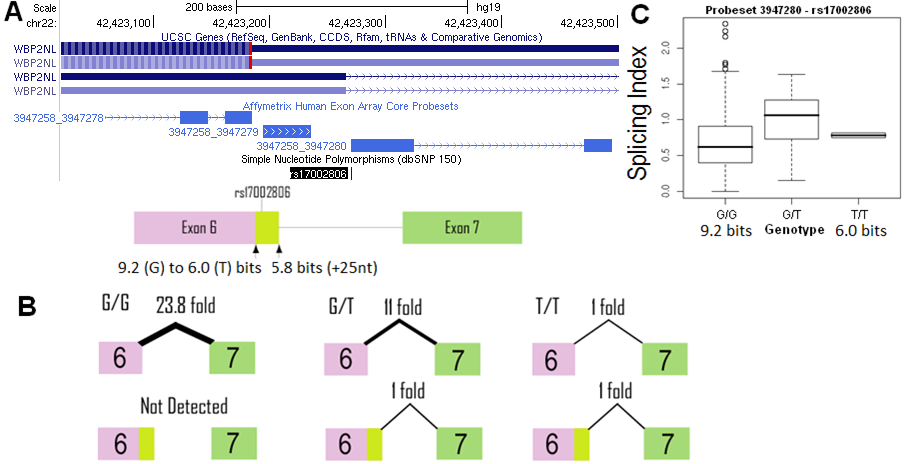

**Supplementary Figure 1.13 (cont.)**

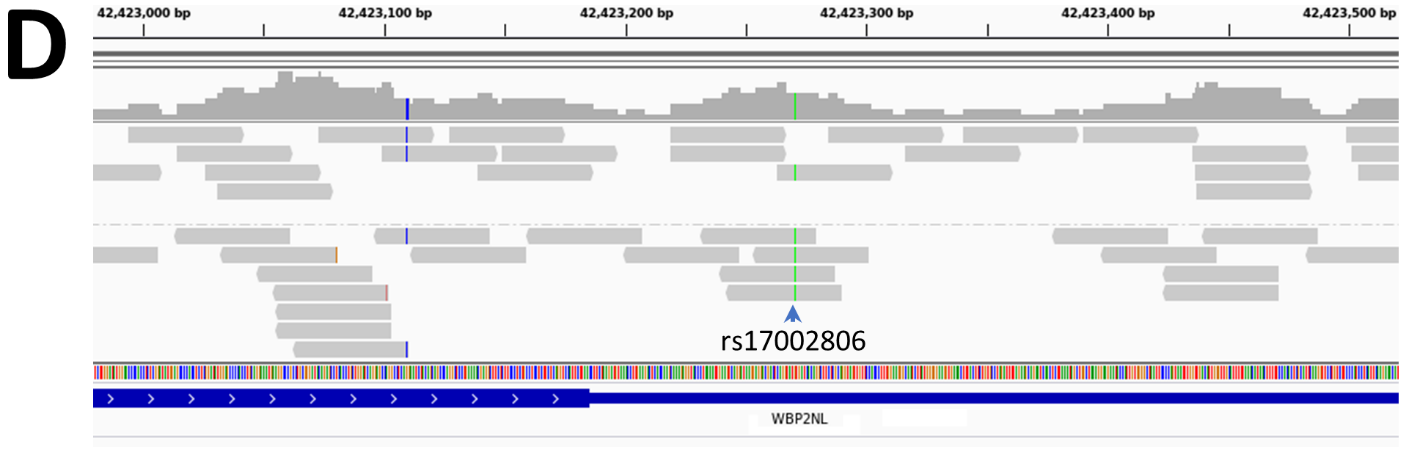

**Supplementary Figure 1.14 – rs2072049 (*PRAME*):** (A) rs2072049 reduces the strength of the acceptor site of the 3’ terminal exon by 1.1 bits (*R_i_* 8.2 > 7.1 bits; G>T). To control for potential confounding effects of rs2266988, individuals used to study rs2072049 were homozygous for the G/G genotype for rs2266988; (B) Individuals with the T-allele for rs2072049 showed decreased *PRAME* expression compared to the homozygous individuals with the 8.2 bit splice site when using q-RT-PCR primers that amplify the affected exon junction (exon 5-6) as well as the control exon 2-3 junction. Individuals carrying the 7.1 bit T-allele had 32.3-57% less *PRAME* expression compared to those with the 8.2 bit G/G genotype; (C) The boxplot measures total *PRAME* intensity (unlike all other boxplots which indicate SI, which is normalized by total gene expression) shows a small step-wise decrease in expression in individuals with the A-allele, congruent with predicted effects on splicing for these genotypes; (D) This variant is flagged in ValidSpliceMut based on an increase in intron retention which could not bet detected by q-RT-PCR. Abnormal splicing observed via RNAseq data in patients with the SNP was uncommon. A single read indicating intron retention [with the SNP; Box 1] is found, however normally spliced mRNA is the dominant isoform (example from heterozygous ICGC MALY patient DO52652).

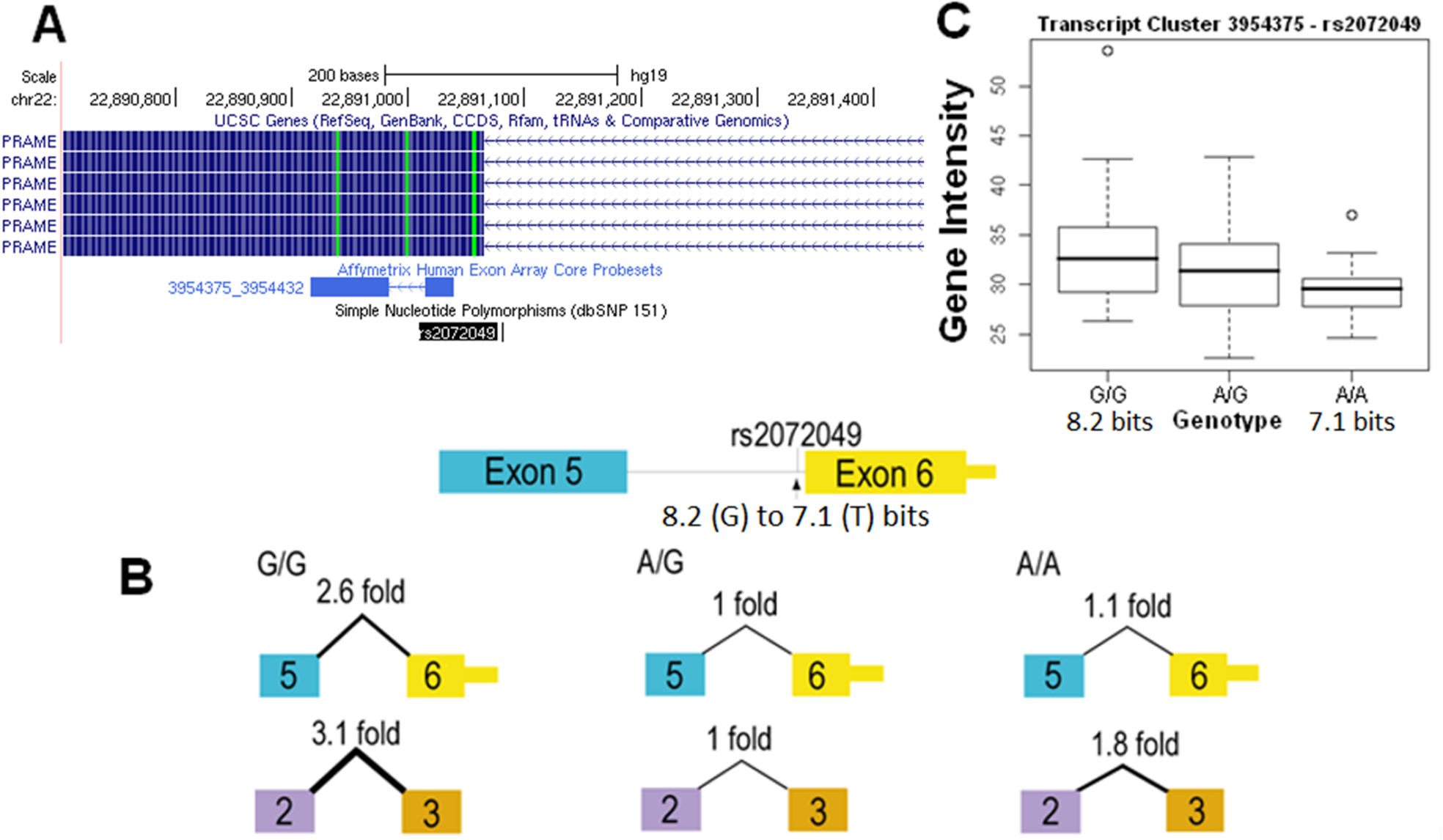

**Supplementary Figure 1.14 (cont.)**

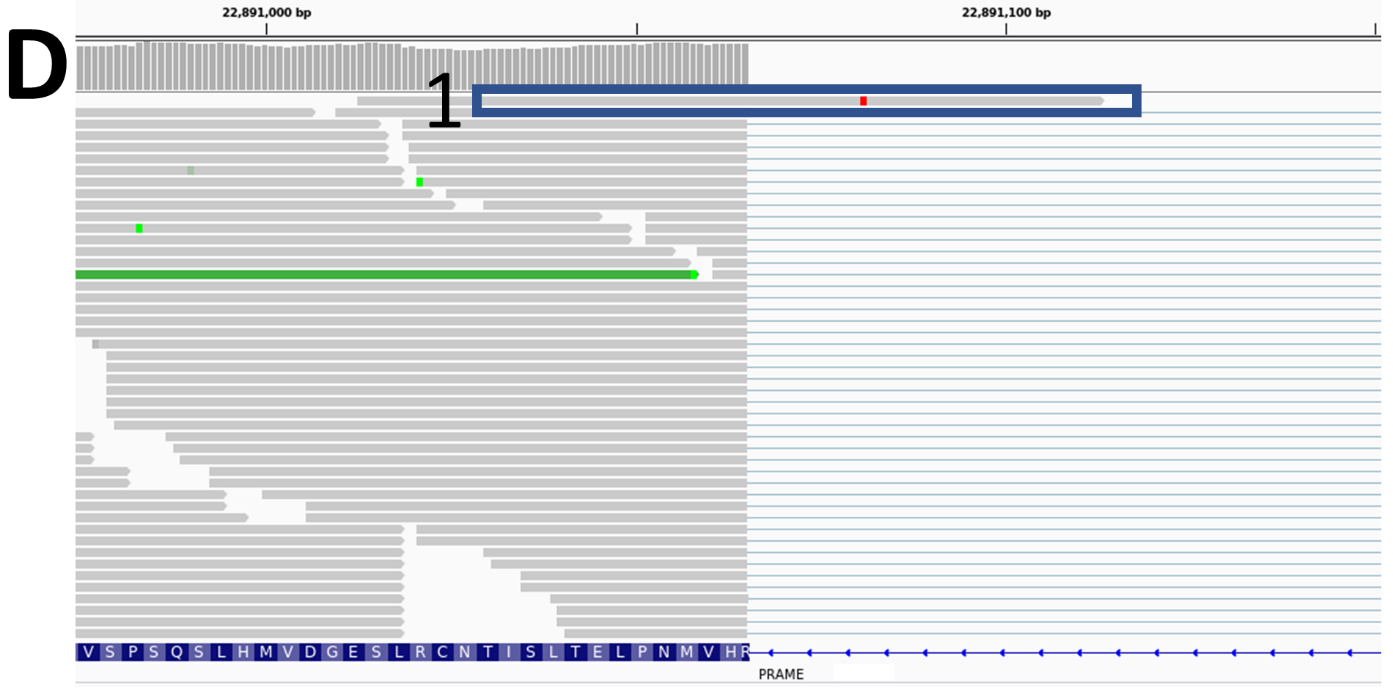

**Supplementary Figure 1.15 – rs6003906 (*DERL3*):** (A) Exon 5 of *DERL3* regularly uses a weak natural acceptor (*R_i_*: -2.1 bits, mRNA CR456372, CU013077, 3 others), while a strong 7.9 bit alternate acceptor exists downstream. Use of this site is well documented in RefSeq (third isoform in panel A), however this mRNA is associated with an alternative version of exon 4 that uses a downstream donor site (*R_i_*=8.0 bits) instead of the upstream donor associated with the affected exon 5 splice form (containing a -4.5 bit site). rs6003906 reduces the information content of the already weak -2.1 bit natural acceptor further by 4.6 fold (to -4.3 bits; A>T); (B) Despite its weak strength, the -4.3 bit acceptor site was still active. The q-RT-PCR assays did not reveal any significant change in the use of the weakened natural site, there was a doubling in the activation of the downstream acceptor site in the homozygous rare individual tested. It is notable that expression of the internal reference of this individual was also elevated to 163% of the wild type level, potentially skewing the results for the SNP-related splicing effects. Although the downstream acceptor is much stronger than the natural splice site, it was used less often (14-29 fold less depending on genotype). A splice form using the upstream exon 4 donor and downstream exon 5 acceptor was not detected; (C) The affected exon is covered by 4 different exon microarray probesets, 3 of which would be expected to detect a decreased SI were the downstream acceptor to be used instead (probeset IDs 3954926, 3954927, 3954928). Two of the three probesets exhibit a decreasing trend in SI (however, 3954927 is not stepwise by genotype), while the fourth probeset (3954925; would detect either splice form) does not change (C; top left). It appears that the T-allele increases the abundance of the alternate splice form from 3% (A/A) to 7% (T/T) of total *DERL3* expression). (D) In RNAseq data of individuals with the SNP (example from TCGA Stomach Adenocarcinoma [STAD] patient TCGA-VQ-A8PQ), the reads present indicate that the cryptic site of exon 5 is indeed used [Box 1] in addition to some intron 4 inclusion [Box 2].

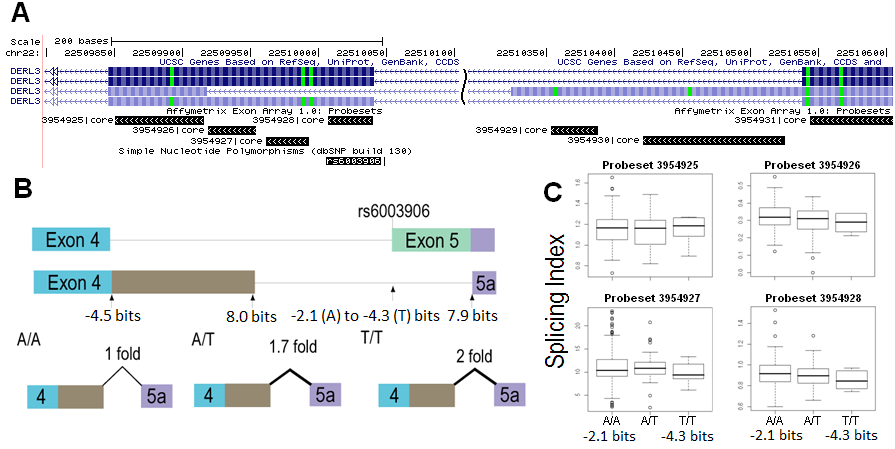

**Supplementary Figure 1.15 (cont.)**

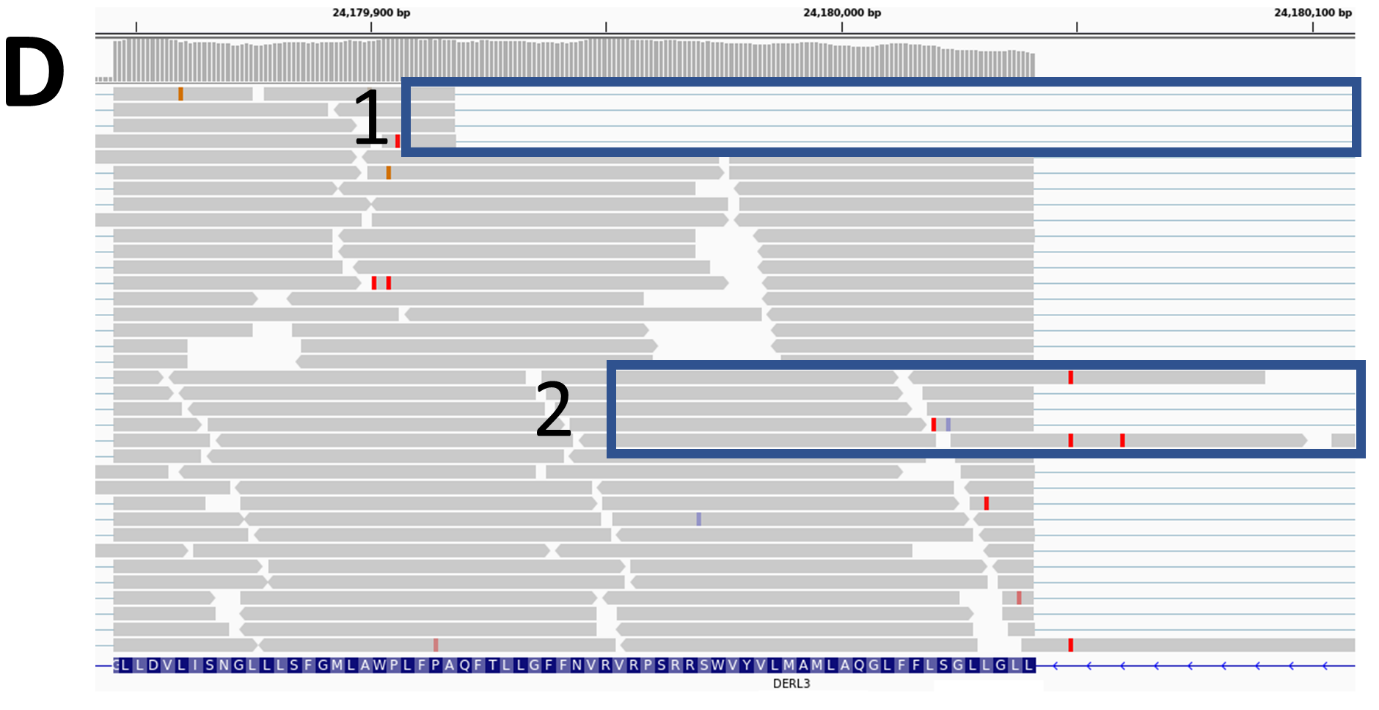

**Supplementary Figure 1.16 – rs2838010 (*FAM3B*):** (A) rs2838010 occurs within intron 1 of *FAM3B* that creates a strong donor site (*R_i_* -10.8 to 7.8 bits, A>T). This donor site occurs at the same coordinate of the donor of a cryptic exon (GenBank Accession AJ409094). The acceptor used by this rare exon is also detected by information analysis (8.2 bits). Both the q-RT-PCR and exon microarray results give inconsistent results according to the predicted effect of the SNP; (B) The single individual homozygous for the strong 7.8 bit allele did not express the rare exon (SI value of ~0). Most individuals tested did have very low SI values for this probe (151 of 176 individuals < 0.1), consistent with the possibility that other trans-acting factors may be required to activate expression of this exon; (C) Expression of the cryptic exon was observed in RNAseq data of heterozygous ICGC MALY patient DO27769 (Box 1), but not in other TCGA and ICGC patients with both *FAM3B* expression who carry the T-allele (i.e. ICGC MALY patient DO27847). Although it is very likely that the T allele is required for the inclusion of this exon, t other regulatory splice sites that either repress exon recognition or may be required for its expression.

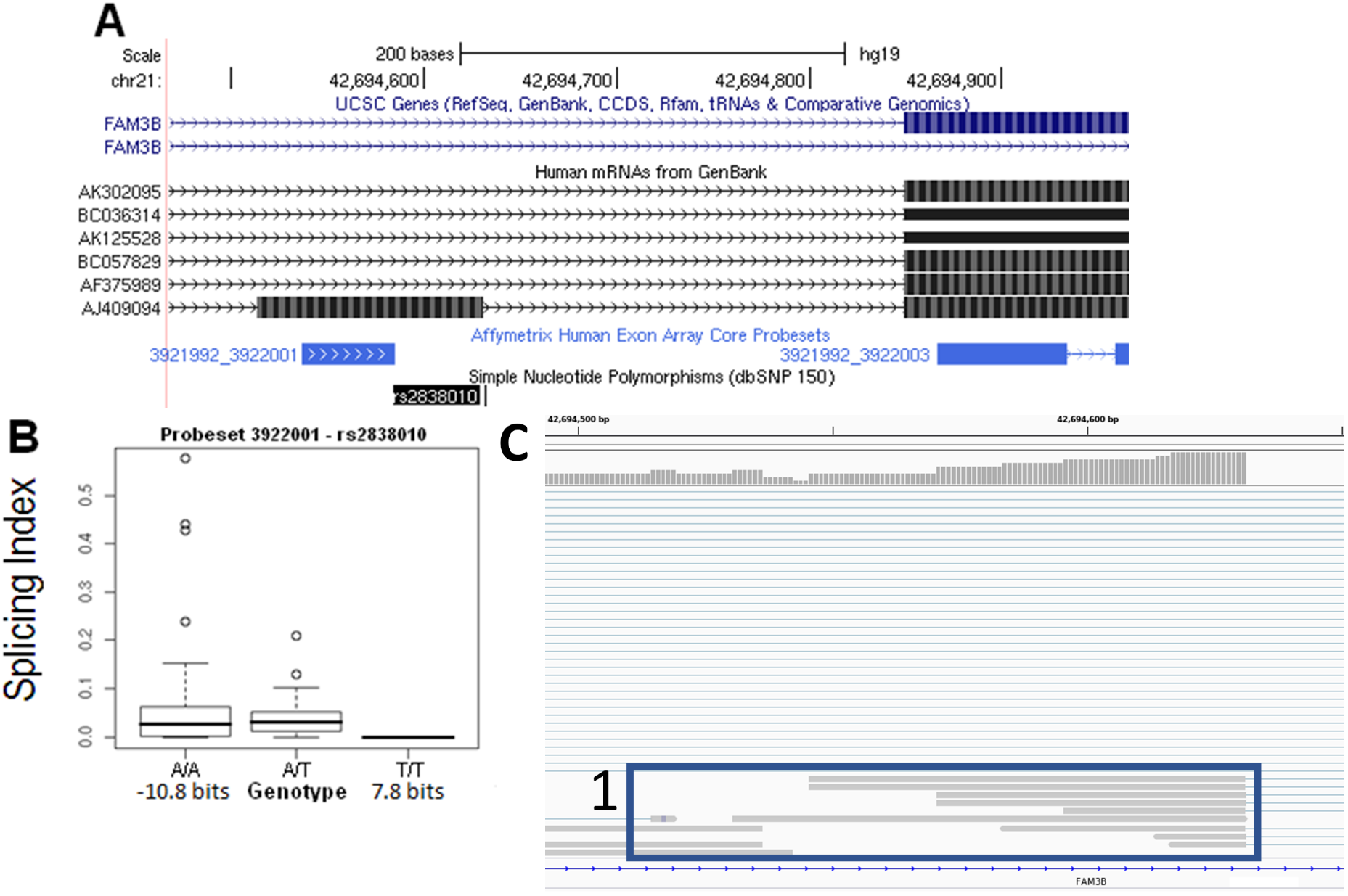

**Supplementary Figure 1.17 – rs16802 (*BCR*):** (A) rs16802 strengthens the acceptor site of exon 14 of *BCR* (*R_i_* 5.6 to 5.8 bits; A>G). The SNP was found by q-RT-PCR to vary 8.3-fold between two homozygous individuals with the stronger site (see Table 1); (B) The exon microarray boxplot is also uninformative, with all 3 genotypes have similar mean SI values; (C) RNAseq data (image from heterozygous ICGC MALY patient DO27769) has exhibits intron retention [Box 1] and exon 14 skipping [Box 2], however normal splicing is the predominant isoform. Variability in splicing regulatory factor abundance can potentially mask statistically significant splicing effects of SNPs such as rs16802.

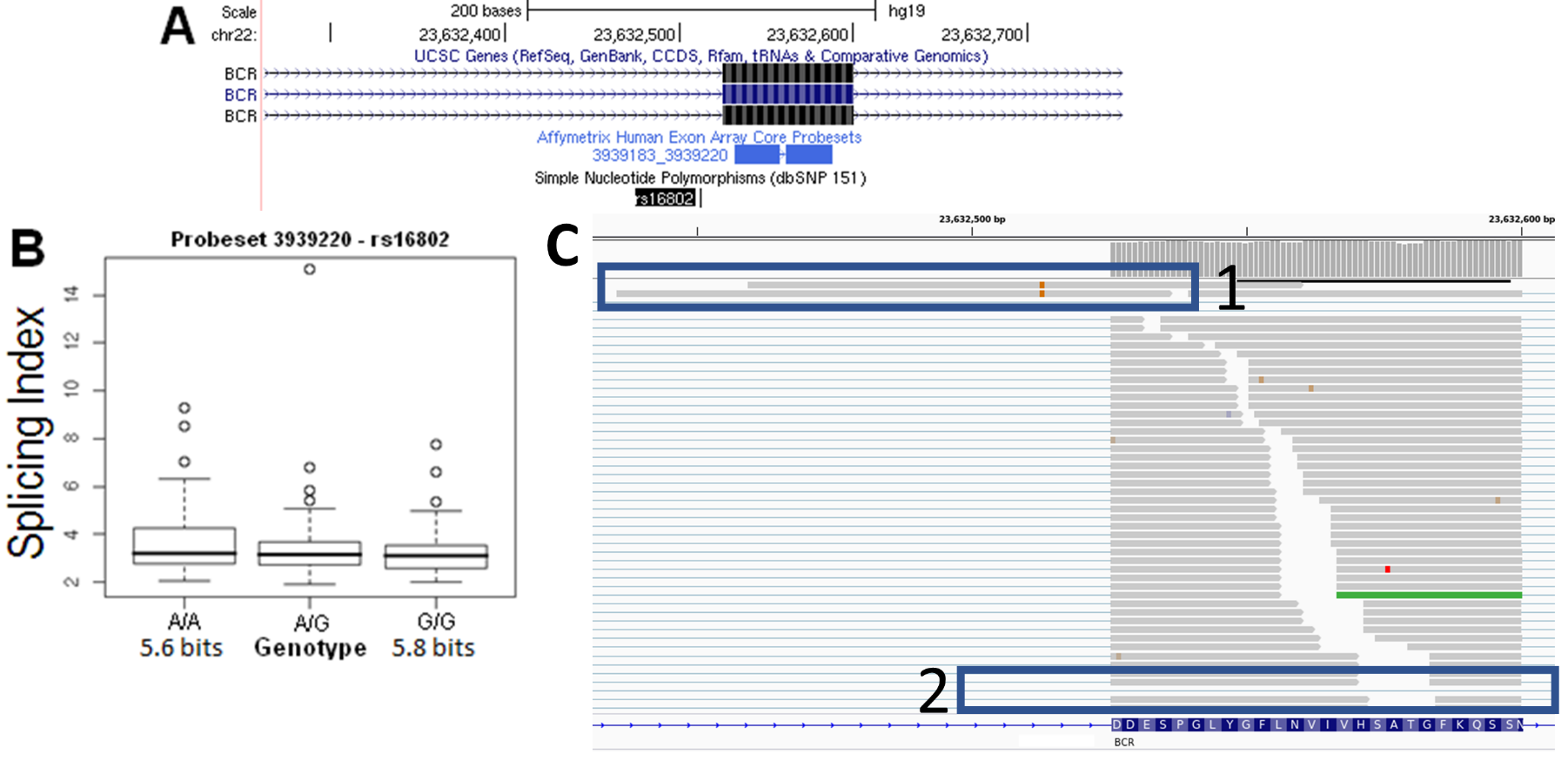

**Supplementary Figure 1.18 – rs2252576 (*BACE2*):** (A) rs2252576 strengthens the natural acceptor exon 5 of *BACE2* by 1.7 fold (*R_i_* 7.2 to 8.0 bits; C>T). However, expression is highly variable between individuals with the same genotype, as splicing across this junction differed by up to 12 fold for homozygous T/T individuals; (B) Individuals with different genotypes exhibited similar mean SI values; (C) RNAseq results indicate predominantly normal splicing for the T genotype, with occasional intron retention [Box 1; image from RNAseq of heterozygous ICGC MALY patient DO52658]. There is little evidence that the rs2252576 genotype influences splicing of this exon, which is consistent with the degree of change predicted by information analysis.

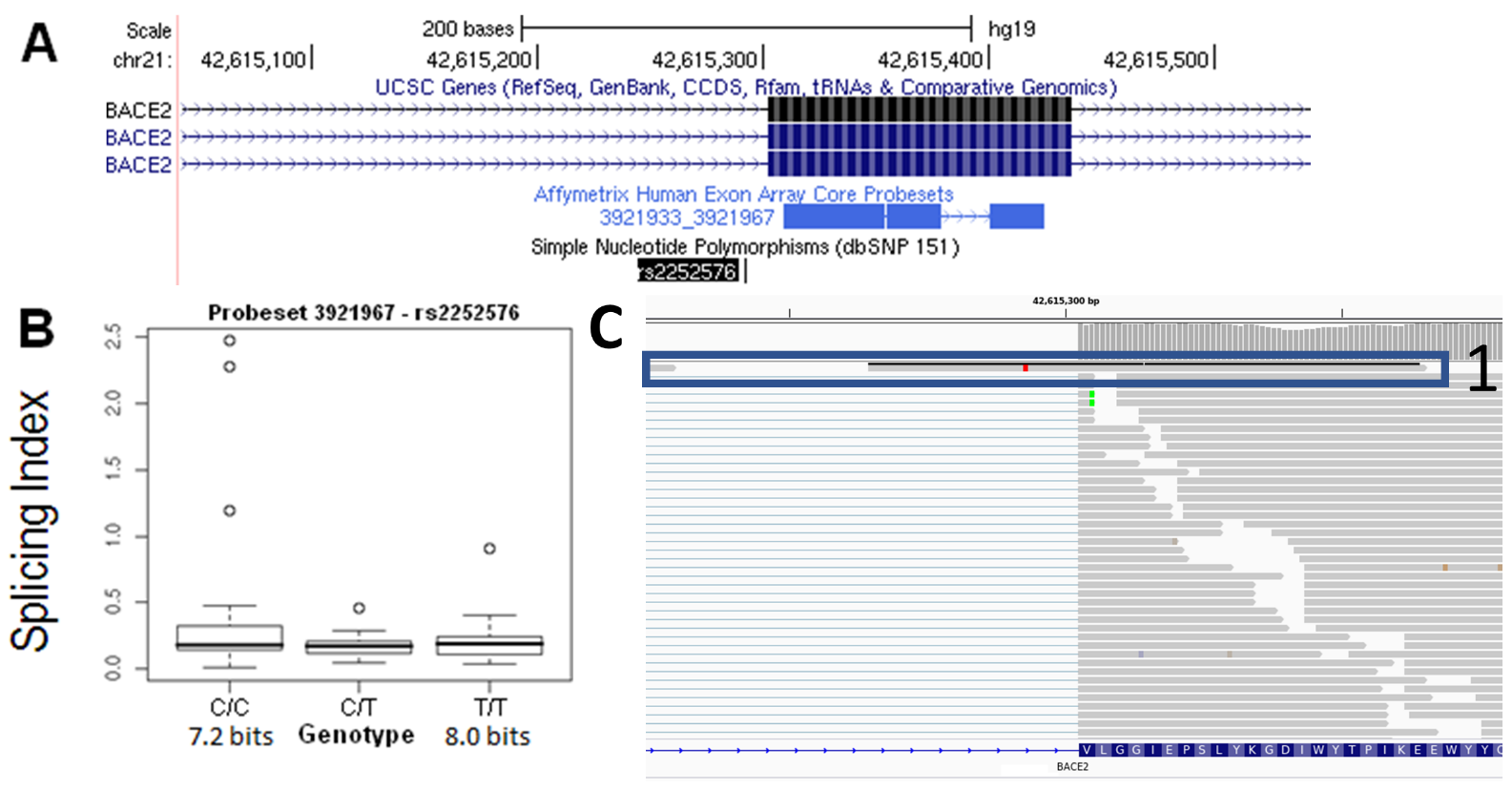

**Supplementary Figure 1.19 – rs8130564 (*TMPRSS3*):** (A) Splicing effects caused by rs8130564 in *TMPRSS3* (natural acceptor weakened from 4.4 to 4.2 bits; C>T) were also masked by variable expression (up to 112 fold difference between homozygous individuals with the stronger splice site); (B) One of the two boxplots (probeset ID 3933585; B; left panel) does indicate a slight, non-stepwise decrease in mean SI for carriers of the T allele, which may not be significant; (C) Normal splicing of this exon was observed in TCGA or ICGC patients carrying this SNP (IGV example from ICGC MALY patient DO52679).

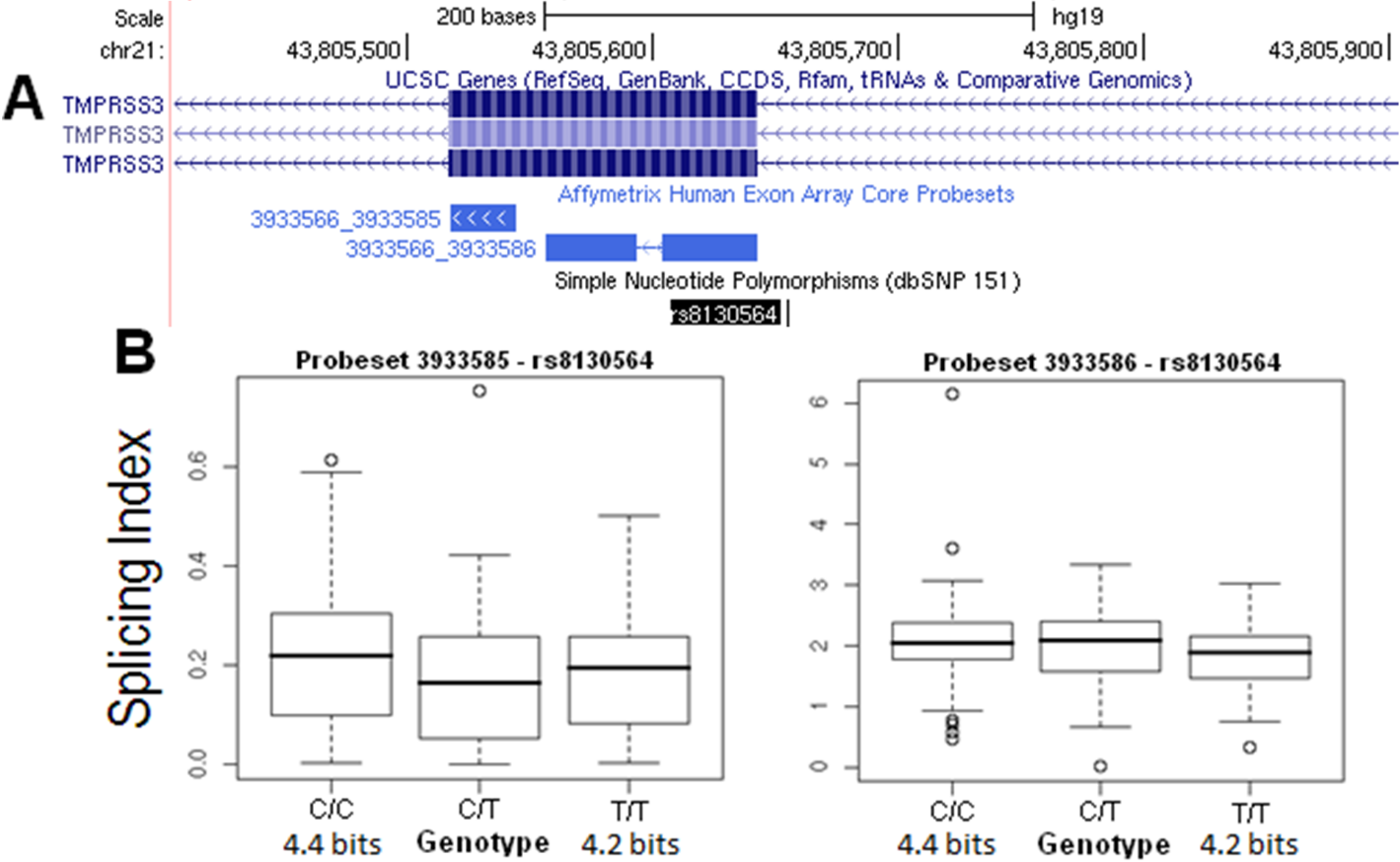

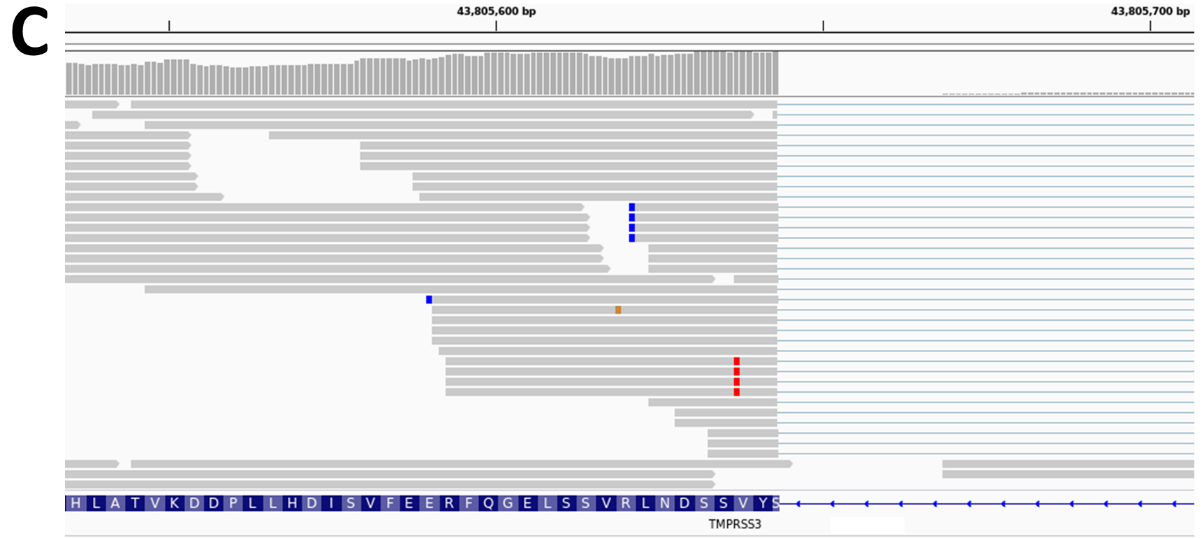

**Supplementary Figure 1.20 – rs2285141 (*CYB5R3*):** (A) rs2285141 decreases the strength of the natural acceptor site of exon 2 of *CYB5R3* (*R_i_* 2.0 to 1.2 bits; G>T); (B) Both q-RT-PCR and microarray boxplots (probeset 3962552) do not support any significant differences in the normal splicing pattern; however, there was an modest increase (1.8 fold) in the inclusion of an alternate exon 2 within intron 1 of *CYB5R3* in individuals with the weak allele; (C) RNAseq data showed no alternative splicing events in any of the patients tested with the T-allele (example of ICGC MALY patient DO52679).

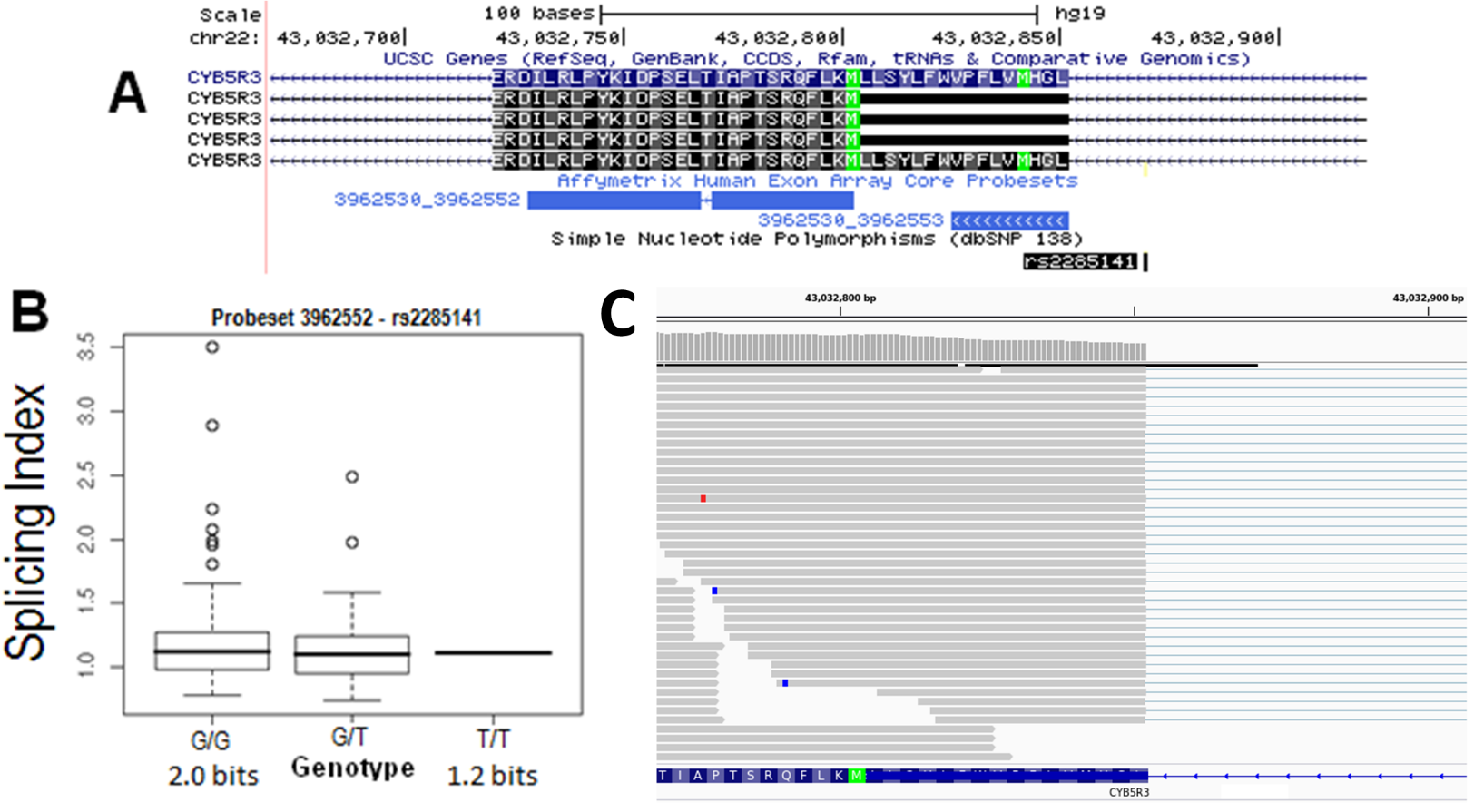

**Supplementary Figure 1.21 – rs2835655 (*TTC3*):** (A) The SNP rs2835655 weakens the natural donor site of *TTC3* exon 39 (*R_i_* 11.2 to 8.2 bits; G>A), reducing its strength by 8.1 fold. q-RT-PCR indicated that normal splicing and total mRNA levels were decreased by ~1.5 fold, which was not significant. Exon skipping was slightly decreased (~1.3 fold), likely due to a decrease in total expression; (B) the microarray data (probeset 3920463) do not support any significant differences in expression between the 3 genotypes; (C) ValidSpliceMut flagged this variant for intron retention (Box 1; example shows ICGC MALY patient DO27793 who is heterozygous for the SNP).

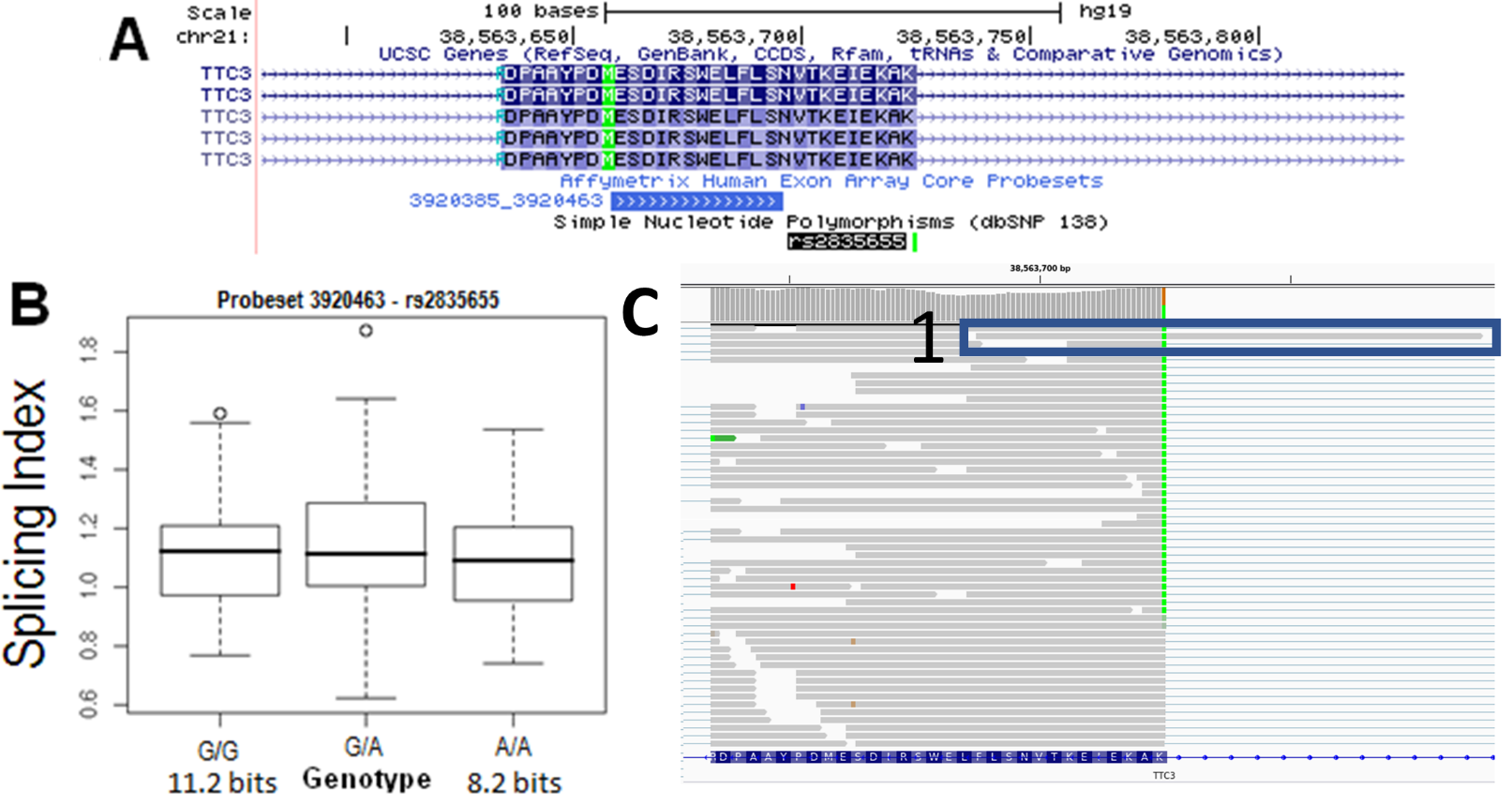

**Supplementary Figure 1.22 – rs16994182 (*CLDN14*):** (A) rs16994182 slightly weakens the natural donor site of exon 2 of *CLDN14* (*R_i_*: 7.4 to 6.8 bits; C>G; panel A). Although decreased levels of normal splicing were detected, this result could not be confirmed due to the lack of an internal reference to account for variation in expression between individuals (gene consists of 3 exons; an internal reference primer set to span the junction between the 6.8 bit exon 2 donor site could not be designed to unequivocally distinguished the expression levels of this isoform from the one generated by the 7.4 bit allele). Exon microarray boxplot and RNAseq data was not available due to a lack of *CLDN14* expression in patients with the G-allele.

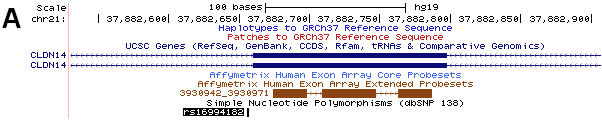
