## Supplementary material for "Expression changes confirm genomic variants predicted to result in allele-specific, alternative mRNA splicing": Supp. Table 1

| **Supplementary Table 1: q-RT-PCR Primers to analyze Splice Forms** | | | | | | |
| --- | --- | --- | --- | --- | --- | --- |
| Splice Form Detection | | Size (nt) | Tm  (ºC) |  | Sequence | Detected |
| External Reference | | | | | | |
| *CCDC137* | Exon 2-3  Junction | 100 nt | 60 | F | AGATTATGAGGAGCCGCCAAG | Yes |
|  |  |  |  | R | GCTTCCTTCTCCAATGTCTTTCTG |  |
| *DNAH1* | Exon 43-44  Junction | 104 nt | 60 | F | CTGATGCGTCACTTCAACTACCTG | Yes |
|  |  |  |  | R | CCAAGGAGTCCAT**CCAACCA** |  |
| *FRMPD1* | Exon 8-9 Junction | 106 nt | 59 | L | GAAGTATCGAGTACTTTGCACTGGC | Yes |
|  |  |  |  | R | TCCTCCCTTTCTACCAC**CTGC** |  |
| *VSP39* | Exon 2-3 Junction | 103 nt | 60 | L | CTCAGTTCCATCAAAACTCG**GTTT** | Yes |
|  |  |  |  | R | GAAGCCAAGACACACAGTGCTG |  |
| Splice Form Detection Primers | | | | | | |
| *XRCC4* rs1805377 (SYBR Green) | | | | |  | |
| **11.5/3.9 bit upstream acc. (6nt included exon 8)** | | 106 nt | 59 | L | AAAAGGAAAA**TTCTAGGCCTGATTCT** | Yes |
|  |  |  |  | R | TCTGGGCTGCTGTTTCTCAGA |  |
| **11.3/11.4 bit downstream acc. (6nt deleted exon 8)** | | 106 nt | 59 | L | TTCAAGAAAAGGAAAA**GCCTGATT** | Yes |
|  |  |  |  | R | TCTGGGCTGCTGTTTCTCAGA |  |
| ***XRCC4* Internal Reference (exon 6-7)** | | 106 nt | 59 | L | GGAAAGTGAAAACCAAACTGATCTCT | Yes |
|  |  |  |  | R | TCTACTTGGTGCAATATCAGTGACATC |  |
| *XRCC4* rs1805377 (dual labelled probe test) | | | | |  | |
| **Primers For Both Probes** | | - | - | L | GAAATCTTGGGACAGAACCTAAA | Yes |
|  |  |  |  | R | TCAGAGTTTCTAAAGACATGTTTTCA |  |
| **Probe- upstream acc. (6nt included exon 8)** | | 141 nt | 61 | P | AGCTTCAAGAAAAGGAAAA**TTCTAGGCCTG** | Yes |
| **Probe - downstream acc. (6nt deleted exon 8)** | | 135 nt | 61 | P | CTTCAAGAAAAGGAAAA**GCCTGATTCTTCA** | Yes |
| *UBASH3A* rs1893592 (SYBR Green) | | | | |  | |
| **9.1/4.3 bit natural donor (exon 10)** | | 107 nt | 60 | L | TCCAGTCCAGAATTGCAG**GG** | Yes |
|  |  |  |  | R | CAGGATGAGTTTGGCCGTCT |  |
| **6.1 bit cryptic donor**  **(Exon 10 + 555nt Extension)** | | 107 nt | 60 | L | ACTCAGAG**CCCTCCGCTGT** | Yes |
|  |  |  |  | R | TCCAGGTTCCACTCGTATCTTGA |  |
| **7.0 bit cryptic donor**  **(Exon 10+29nt extension)** | | 107 nt | 60 | L | TTTGAGGACTGTCTAGTAGGAAAG**GG** | Yes |
|  |  |  |  | R | GAGTTTGGCCGTCTGCACA |  |
| **Exon 10 Skipping** | | 119 nt | 65 | L | CCTTGCAG**GCTACCGTTGCAAG** | No |
|  |  |  |  | R | TAGCGCGTC**CCCATCAGGAGT** |  |
| ***UBASH3A* Internal Reference (Exon 8-9)** | | 107 nt | 60 | L | TACAGGCCTTGCAG**GCTACC** | Yes |
|  |  |  |  | R | AGTGGAGCATTGCTGCAGC |  |
| *C21ORF2* rs2070573 (SYBR Green) | | | | |  | |
| **10.4 bit natural donor (exon 6)** | | 96 nt | 62 | L | CCCAGGATGAACGTGGCCT | Yes |
|  |  |  |  | R | GGACG**TTCCTGCCCCTGTG** |  |
| **5.7/2.4 bit cryptic donor (Exon6+360nt Extension)**  **G version^1^** | | 96 nt | 62 | L | GGGCAACAGGAGTCACGTGG | Yes |
|  |  |  |  | R | GCAGTCAGGACGTT**CGGCA** |  |
| **5.7/2.4 bit cryptic donor (Exon6+360nt Extension)**  **C version^1^** | | 106 nt | 62 | L | TGAGGGCAACAGGAGTCACG | Yes |
|  |  |  |  | R | CAGGATGGCAGTCAGGACGTT**A** |  |
| ***C21ORF2* Internal Reference (Exon 4-5)** | | 96 nt | 62 | L | TGGACAACCAGG**CTGTGACG** | Yes |
|  |  |  |  | R | CGTGGCCTGTGCCCTCTCT |  |
| *EMID1* rs743920 (SYBR Green) | | | | |  | |
| **6.4 bit upstream acc. (6nt included exon 4)** | | 108 nt | 59 | L | GAGCTGCGAGGAAG**TTGCAG** | Yes |
|  |  |  |  | R | TTGAGACAACCTGAGAAGGCTGT |  |
| **10.5/12.4 bit downstream acc. (6nt deleted exon 4)**  **C version^1^** | | 85 nt | 59 | L | GTACAAGATAGTGACCGCCCGT | Yes |
|  |  |  |  | R | CAAGGAGGCAGAGGAAC**CTTC** |  |
| **10.5/12.4 bit downstream acc. (6nt deleted exon)**  **G version^1^** | | 85 nt | 59 | L | GTACAAGATAGTGACCGCCCGT | Yes |
|  |  |  |  | R | CAAGGAGGCAGAGGAAG**CTTC** |  |
| ***EMID1* Internal Reference 1 (exon 4-5) (shorter amplicon)** | | 85 nt | 59 | L | CTTCTCAG**GTTGTCTCAACTGCAG** | Yes |
|  |  |  |  | R | AGTCAGCATGGTCAT**CTTGGC** |  |
| ***EMID1* Internal Reference 2 (exon 4-5) (longer amplicon)** | | 111 nt | 59 | L | CAGCCTTCTCAG**GTTGTCTCAAC** | Yes |
|  |  |  |  | R | GAGGTACTGGCTGCTCTATGACAG |  |
| *IL19* rs2243187 (SYBR Green) | | | | |  | |
| **7.3/-0.3 bit upstream acc. (3nt included exon 5)** | | 101 nt | 63 | L | AGCCAAACCCCAAAATCTTGAGAA | Yes |
|  |  |  |  | R | GTGACACTGCCTCTGTTCCTG**ACAT** |  |
| **7.6/7.5 bit downstream acc. (3nt excluded exon 5)** | | 101 nt | 63 | L | CAGGAGCCAAACCCCAAAATCTT | Yes |
|  |  |  |  | R | TGACACTGCCTCTGTTC**ACATTGC** |  |
| **Exon 5 Skipping** | | 104 nt | 63 | L | CAAGGATCATCAGGAGCCAAACC | Yes |
|  |  |  |  | R | GGACCTCCAG**ACATTGCCGC** |  |
| ***IL19* Internal Reference (Exon 2-4)** | | 102 nt | 63 | L | AAAAGAGCCATC**CAAGCTAAGGACAC** | Yes |
|  |  |  |  | R | GGTCACGCAGCACACATCTAAGG |  |
| *PRAME* rs2266988 (SYBR Green) | | | | |  | |
| **8.9/7.3 bit natural donor (exon 3)** | | 101 nt | 65 | L | TGAGACCTAGAAATCCAAGCGTTGGA | Yes |
|  |  |  |  | R | GATGTATCGGCTCTGAATGGAACC**C** |  |
| **Exon 3 Skipping** | | 95 nt | 65 | L | TGGTGAACTCTCTGAGGAAAAAC**GGT** | Yes |
|  |  |  |  | R | CCCTGCCAGCTCCACAAGTCTC |  |
| ***PRAME* Internal Reference (Exon 5-6)** | | 101 nt | 65 | L | TCAGTTGCTCAG**GCACGTGATGA** | Yes |
|  |  |  |  | R | CGCTGGGACTCTGGGACAGATG |  |
| *TTC3* rs2835585 (SYBR Green) | | | | |  | |
| **6.4/4.4 bit natural acc. (exon 3)** | | 102 nt | 64 | L | TTATGTTCGTGTGACTCAGCTTTACTGTGA | Yes |
|  |  |  |  | R | CATATACTGCAGATGTCAAATT**CCAAATTCC** |  |
| **Exon 3 Skipping** | | 102 nt | 64 | L | ATGATTATGTTCGTGTGACTCAGCTTTACTG | Yes |
|  |  |  |  | R | TCAGAAGCACGTGAATTG**CCAAA** |  |
| **Intron 2 (IVS2) Inclusion** | | 101 nt | 60 | L | GGGTG**GGTGTGCAATATAAAGATT** | Yes |
|  |  |  |  | R | CCTCCCTTTGTAGTCTTTCCATAAATTAA |  |
| **6.9 bit cryptic acceptor (exon 3 + 60 nt)** | | 105 nt | 64 | L | TTATGTTCGTGTGACTCAGCTTTACTGTGA | No |
|  |  |  |  | R | GCCTCATTTTTTTAATGAATCAACA**CCAA** |  |
| **7.2 bit cryptic acceptor (exon 3 + 87 nt)** | | 107 nt | 64 | L | TCGTGTGACTCAGCTTTACTGTGATGG | No |
|  |  |  |  | R | TCAACACTTAATAGTAAAAAGGCAAAAATGGAT**C** |  |
| ***TTC3* Internal Reference (Exon 4-6)** | | 102 nt | 64 | L | GATGGAAGATATTGTGGATTTGGCAAAG | Yes |
|  |  |  |  | R | GCCAAGATTTT**ATTTTCTATTTTACAACCAATTCTC** |  |
| *TTC3* rs2835655 (SYBR Green) | | | | |  | |
| **Natural Exon 38-39 Junction (unaffected)** | | 113 nt | 64 | L | GTACTTGAAAACTGGAAGGAGAGTGAAGTGTATAAG | Yes |
|  |  |  |  | R | GGATATGCTGCAGGATCA**CGGCTA** |  |
| **Exon 39 Skipping** | | 113 nt | 65 | L | AAAACTGGAAGGAGAGTGAAGTGTATAAGCTACAG | Yes |
|  |  |  |  | R | CCTTAATTTGTTCTTCAAACTGAGA**CGGC** |  |
| ***TTC* Internal Reference (Exon 40-41)** | | 113 nt | 65 | L | AGAACAAATTAAGGCAATTAAAAATGGTTCTCG | Yes |
|  |  |  |  | R | CAGGGAGTAACTCGGGATGAAC**CGT** |  |
| *FAM3B* rs2838010 (SYBR Green) | | | | |  | |
| **Natural Exon 1-2 Junction (unaffected)** | | 95 nt | 59 | L | CTGGTG**GCCTGCTCAAGGT** | Yes |
|  |  |  |  | R | GGGTGCATCTGGAATGAGCT |  |
| **1.4/9.2 bit donor of cryptic exon within IVS1** | | 93 nt | 59 | L | CCATTGGCTGGTG**GTTCAC** | No |
|  |  |  |  | R | TTAACTTAACCAACAAACGTCCTTGA |  |
| ***FAM3B* Internal Reference (exon 3-5)** | | 96 nt | 59 | L | GCCCATCTGACACCTATGCC | Yes |
|  |  |  |  | R | GTTCTCCCATAAGT**AGGTTATCCTCAA** |  |
| *WBP2NL* rs17002806 (SYBR Green) | | | | |  | |
| **10/6.5 bit natural donor (exon 6)** | | 90 nt | 60 | L | AGGATAAGGAGGACGACTCAG**CTT** | Yes |
|  |  |  |  | R | CTGTCCTTGAGAACATGGGAGC |  |
| **Exon 5-6 Junction (unaffected)** | | 109 nt | 60 | L | TTTCCACTTAGAACCTTAAATGACTGGT | Yes |
|  |  |  |  | R | CATAGACAATAA**CTGAACAAGGCATCTG** |  |
| **Exon 6 Skipping** | | 101 nt | 60 | L | AAATGACTGGTTCAGCTCTATGGG | No |
|  |  |  |  | R | AGATCAATCAATGACTCTAAG**CTGAACA** |  |
| **6.6 bit cryptic donor (exon 3 + 25 nt)** | | 101 nt | 59 | L | AGACTCACCAAGCAAAGAGGTACC | Yes |
|  |  |  |  | R | CAATCAATGACTCTAAG**CTGCGAG** |  |
| **5.7 bit cryptic donor (exon 3 + 67 nt)** | | 101 nt | 60 | L | GGAGGACGACTCAGGTATGTGATC | No |
|  |  |  |  | R | GATCAATCAATGACTCTAAG**CTTCCC** |  |
| ***WBP2NL* Internal Reference (exon 4-5)** | | 106 nt | 60 | L | TGGTGAAAGCTGCCTCTGCT | Yes |
|  |  |  |  | R | CACATATTCCCTTCCCCAGTAATTAC |  |
| *IFI44L* rs1333973 (SYBR Green) | | | | |  | |
| **9.5/5.0 bit natural acceptor (exon 2)** | | 94 nt | 59 | L | CCGTGGCTGCTCGATAAATC | Yes |
|  |  |  |  | R | TTGTCACTTCCATTGTTCTATAT**CTGTTT** |  |
| **Exon 2 Skipping** | | 94 nt | 59 | L | AACCGTGGCTGCTCGATAAA | Yes |
|  |  |  |  | R | CGTCTAGGTTATCCTTAATTC**CTGTTTC** |  |
| ***IFI44L* Internal Reference (exon 5-6)** | | 94 nt | 59 | L | TGTATGCCAGACAGATATCAG**TTTAATTC** | Yes |
|  |  |  |  | R | GAATCCTGTCCTTCAGAGATGGAG |  |
| *CFLAR* rs10190751 (SYBR Green) | | | | |  | |
| **17.4/9.9 bit acceptor of upstream exon 7 (s form)** | | 93 nt | 59 | L | AGCAGGGACAAGTTACAGGAATGT | Yes |
|  |  |  |  | R | GCATAGGGTGTTATCAT**CCTGAAGT** |  |
| **6.9 bit acceptor of downstream exon 7** | | 94 nt | 59 | L | CAAGGAGCAGGGACAAGTTACAG | Yes |
|  |  |  |  | R | TCCCATTATGGAG**CCTGAAGTT** |  |
| ***CFLAR* Internal Reference (exon 3-4)** | | 91 nt | 59 | L | AATCTGATGTGTCCTCATTAATTTTCC | Yes |
|  |  |  |  | R | ACCACAAGGTCCAAGAAACT**CTTC** |  |
| *LPP* rs13076750 (SYBR Green) | | | | |  | |
| **9.3/-1.6 bit upstream acc. (7nt included exon 1a)** | | 91 nt | 60 | L | TTCCCTGTGTTCTGCTTTTTTCAT | Yes |
|  |  |  |  | R | CCAACTGCAATGCTAGTGT**CAATT** |  |
| **2.0 bit downstream acc. (7nt excluded exon 1a)** | | 89 nt | 60 | L | TGATTCCCTGTGTTCTGCTTTTTT | Yes |
|  |  |  |  | R | TCAGCCAACTGCAATG**CAAT** |  |
| **Exon 1a Skipping (cryptic exon in IVS1)** | | 89 nt | 60 | L | TCCCTGTGTTCTGCTTTTTTCAT | Yes |
|  |  |  |  | R | GGGTGAGACATTGTTGGAAT**CAAT** |  |
| ***LPP* Internal Reference (exon 3-4)** | | 90 nt | 60 | L | CTGGAGGTGAGG**GTGATTTTCTT** | Yes |
|  |  |  |  | R | GAGGAGGAAAGTTTCCAGAGATAGATG |  |
| *GUSBP11* rs3747107 (SYBR Green) | | | | |  | |
| **Upstream Exon 12 (8.7/1.4 bit acc.)** | | 71 nt | 59 | L | TCATGACTAACCAGT**AGTGGGTGC** | Yes |
|  |  |  |  | R | CTGGTCGTAGGCTGGATGTGT |  |
| **1.6/7.5 bit upstream cryptic Acc. (-2 nt)** | | 69 nt | 59 | L | TCATGACTAACCAGT**TGGGTGC** | No |
|  |  |  |  | R | CTGGTCGTAGGCTGGATGTGT |  |
| **8.3 bit upstream cryptic acc. (-114 nt)** | | 76 nt | 58 | L | CCGATTTCATGACTAACCAGT**GAT** | Yes |
|  |  |  |  | R | AGAGTCTCCTTGGGAAACTTACCA |  |
| **10.9 bit upstream cryptic acc. (-118 nt)** | | 76 nt | 59 | L | TTTCATGACTAACCAGT**TTAGGATGG** | Yes |
|  |  |  |  | R | AGAGTCTCCTTGGGAAACTTACCA |  |
| **6.0 bit upstream cryptic acc. (-156 nt)** | | 69 nt | 59 | L | GCCGATTTCATGACTAACCAGT**GT** | No |
|  |  |  |  | R | CCATCCTAACTGGGAAGACAAAAAG |  |
| **Downstream Exon 12 (8.9 bit acc.)** | | 70 nt | 59 | L | CCGATTTCATGACTAACCAGT**GTC** | Yes |
|  |  |  |  | R | TCACAGGACGGCAGGAACA |  |
| ***GUSBP11*** **Internal Reference (exon 10-11)** | | 69 nt | 59 | L | GTGGACATTGACCCCACTGG | Yes |
|  |  |  |  | R | GAACATCAGAGGTGGATC**CTGC** |  |
| *DERL* 3 rs6003906 (SYBR Green) | | | | |  | |
| **2.2/0.3 bit natural acceptor (exon 5)** | | 92 nt | 59 | L | ATGCTGGAAGAGGGCTCCTT | Yes |
|  |  |  |  | R | AGGAGTCCCAGCAG**GGTCA** |  |
| **11.3 bit downstream alt. acc. (exon 5 – 123nt) Uses extended exon 4** | | 95 nt | 59 | L | GAGGGCCCACTCTGTGCTC | Yes |
|  |  |  |  | R | ACGGTGC**CTGGCTCTTTTG** |  |
| **11.3 bit downstream alt. acc. (exon 5 – 123nt) Uses short exon 4** | | 89 nt | 59 | L | ATGCTGGAAGAGGGCTCCTT | No |
|  |  |  |  | R | AGGAACGGTGC**GGTCATAAG** |  |
| ***DERL3* Internal Reference (exon 2-3)** | | 77 nt | 59 | L | GGAGCTCCTCAGCCCCTTT | Yes |
|  |  |  |  | R | TGACGAGCCTCCAGAC**CTG** |  |
| *ARFGAP3* rs1018448 (SYBR Green) | | | | |  | |
| **10.6/12.8 bit natural acceptor (exon 12)** | | 100 nt | 59 | L | AATCACCCATTATGGCAAAACC | Yes |
|  |  |  |  | R | AACTCCACTGGCTCGTCAAAGT |  |
| **5nt upstream acc. (exon 12)**  *** Potentially misaligned** | | 101 nt | 59 | L | CAGGAATCACCCATTATGGCA | No |
|  |  |  |  | R | TTAACTCCACTGGCTCGTCAA**CT** |  |
| **Exon 12 Skipping** | | 100 nt | 60 | L | ACCATAGAGCAGGAATCACCCAT | Yes |
|  |  |  |  | R | GAGCAGTAGGT**CTTGAGCTGGAAG** |  |
| ***ARFGAP3* Internal Reference (exon 13-14)** | | 100 nt | 58/60 | L | ATACAGATGAGGCCCAGAAGAAGTT | Yes |
|  |  |  |  | R | CGGGCCCTGGTCTCATA**AT** |  |
| *BCR* rs16802 (SYBR Green) | | | | |  | |
| **8.8/9.4 bit natural acceptor (exon 14)** | | 91 nt | 59 | L | GCTGCAGATGCTGACCAACTC | Yes |
|  |  |  |  | R | CCCCGGAGACTCATCAT**CTTC** |  |
| ***BCR* Internal Reference (exon 10-11)** | | 93 nt | 59 | L | ATCCAGAGAGAGAAG**AGGGCG** | Yes |
|  |  |  |  | R | CATAAGCAGCAGCAGTGACTCC |  |
| *BACE2* rs2252576 (SYBR Green) | | | | |  | |
| **9.0/9.6 natural acceptor (exon 5)** | | 97 nt | 60 | L | CTTAATCTGGACTGCAGAGAG**TATAACG** | Yes |
|  |  |  |  | R | CCACCGCATCAAACACCTTC |  |
| ***BACE* Internal Reference (exon 3-4)** | | 97 nt | 60 | L | TTAAATGGAATGGAATACTTGGCCT | Yes |
|  |  |  |  | R | TTTGCTTGTGTCACCAGGGA |  |
| *TMPRSS3* rs8130564 (SYBR Green) | | | | |  | |
| **6.8/6.3 natural acceptor (exon 6)** | | 103 nt | 59 | L | CATGTGCTCCGATGACTGGA | Yes |
|  |  |  |  | R | CAGCGAGCTCACTCTGAGGTTAT |  |
| ***TMPRSS3* Internal Reference (exon 7-8)** | | 103 nt | 59 | L | GTGCACAG**CCTGTGGTCATAGA** | Yes |
|  |  |  |  | R | CTGGAACTGAAGGCTGGCC |  |
| *CLDN14* rs16994182(SYBR Green) | | | | |  | |
| **Natural Exon 2-3 Junction** | | 94 nt | 63 | L | CTTTCAGATATAGCACTGGACTTGGCTG | Yes |
|  |  |  |  | R | ACAGTTCTTGACTTCTTGGCTTGTGTC |  |
| **Exon 2 Skipping** | | 95 nt | 63 | L | CTGCCAAGAGAGGGAGTAAGATGTTCA | Yes |
|  |  |  |  | R | GGAGCC**CGTGCTGCTGTGT** |  |

**Supplementary Table 1: q-RT-PCR Primers to analyze Splice Forms**

**Supplementary Table 1: q-RT-PCR Primers to analyze Splice Forms**

This table lists all primers for all SYBR Green q-RT-PCR experiments described in this paper. The top four primer sets were simultaneously run with test primers to normalize sample-to-sample variation. Each group of primer sets which tested the effects of a specific mutation were designed to be similar in amplicon size and T_m_, though this was dependant on sequence. Primers designed to detect a specific splice form were placed over an exon junction (unless sequence of junction is detrimental to primer design, i.e. contains a G quartet, palindromic sequence). The tested splice form is described in the left-most column, while whether or not that splice form was detected is in the right-most column. Internal references are designed to amplify regions of the gene not expected to be affected by the SNP and is used to detect potential variation in the gene between individuals tested. The bolded regions of the primer sequence indicate the region of the primer that crosses the exon junction. Legend: F – Forward primer, R – Reverse primer, Acc. – Acceptor splice site. **^1^** When the mutation is located on the +1 or +2 positions of the splice site (within the exon), primers placed over the junction would lay on the polymorphism and therefore two primer sets are designed to account for this, i.e. *C21orf2* 5.7/2.4 bit cryptic donor (Exon 6 + 360nt Extension) G version with 100% match when tested mutation is a G. Note in Table 1 (q-RT-PCR tables) that primers that mismatch will often still accurately measure the change in splicing.
