## Supplementary material for "Expression changes confirm genomic variants predicted to result in allele-specific, alternative mRNA splicing": Supp. Table 2

| **Supplementary Table 2: SNPs in ValidSpliceMut Splicing Mutation Database** | | | | |
| --- | --- | --- | --- | --- |
| **Gene** | **rsID** | **Variant^1^** | **# Matches to ValidSpliceMut** | **Veridical Evidence Type** |
| *XRCC4* | rs1805377 | [5:82648943G>A](https://validsplicemut.cytognomix.com/view.php?targets=907750+898346+898347+898348+898349+907749+933253+907751+933252+898344+933254+933255+933256+933257+898345+819965+898343+191626+78198+191621+191622+191623+191624+191625+191627+898342+813246+813247+813248+819964+898341+933258) | 32 | Patients flagged for intron inclusion RNAseq reads |
| *IL19* | rs2243187 | 1:207014348G>A | - | - |
| *C21orf2* | rs2070573 | [21:45750346C>A](https://validsplicemut.cytognomix.com/view.php?targets=191610+812565+191609+79880+191608) | 5 | Patients flagged for junction-spanning and total intron inclusion RNAseq reads |
| *TTC3* | rs2835665 | 21:38574092A>G | - | - |
| *TTC3* | rs2835585 | [21:38460488 T>A](https://validsplicemut.cytognomix.com/view.php?targets=79127&referenceName=21&alternateBases=A&start=38460488&referenceBases=T&assemblyId=GRCh37&includeDatasetResponses=HIT) | 1 | Patient flagged for intron inclusion RNAseq reads |
| *WBP2NL* | rs17002806 | 22:42423271G>A | - | - |
| *GUSBP11* | rs3747107 | 22:23995540C>G | - | - |
| *PRAME* | rs2266988 | 22:22899234A>G | - | - |
| *PRAME* | rs2072049 | [22:22891081G>T](https://stagedvalidsplicemut.cytognomix.com/view.php?targets=80564+427772+427773) | 3 | Patients flagged for intron inclusion RNAseq reads and total intron inclusion |
| *UBASH3A* | rs1893592 | [21:43855067A>C](https://validsplicemut.cytognomix.com/view.php?targets=426812+426813+426814+426815+426816+426817+426818+426819+447351) | 9 | All patients flagged for intron inclusion; 3 patients also flagged for exon skipping reads |
| *DERL3* | rs6003906 | 22:24180049A>T | - | - |
| *ARFGAP3* | rs1018448 | 22:43206950A>C | - | - |
| *CFLAR* | rs10190751 | [2:202006096G>A](https://validsplicemut.cytognomix.com/view.php?targets=791453+554189+554190+554193+554194+791452+791454+554187+791455+813970+813971+813972+813973+917749+554188+554186+73687+428256+79783+428255+428257+428258+428259+428260+428261+428262+428263+428264+917750) | 29 | Patients flagged for intron inclusion RNAseq reads |
| *IFI44L* | rs1333973 | 1:79094081A>T | - | - |
| *LPP* | rs13076750 | [3:188059443A>G](https://validsplicemut.cytognomix.com/view.php?targets=349138+349139+633447+633448+797460+797461+922409+922410)^2^ | 8 | Patients flagged for intron inclusion and cryptic |
| *EMID1* | rs743920 | 22:29621128G>C | - | - |
| *CLDN14* | rs16994182 | 21:37882657C>G | - | - |
| *BCR* | rs16802 | 22:23632513A>G | - | - |
| *TMPRSS3* | rs8130564 | 21:43805656T>C | - | - |
| *BACE2* | rs2252576 | 21:42615293C>T | - | - |
| *CYB5R3* | rs2285141 | 22:43032870G>T | - | - |
| *FAM3B* | rs2838010 | 21:42694633A>T | - | - |
| ^1^ Hyperlinked variant (with HG19 coordinates) lead to ValidSpliceMut page for the variant of interest. ^2^ LPP exon affected is rarely used and is categorized in ValidSpliceMut as a cryptic site splicing change. As the software did not consider the affected exon, any exon skipping reads would be instead for the nearest associated exon, which explains why exon skipping is not flagged in ValidSpliceMut despite its considerable increase detected by qRT-PCR. | | | | |
