## Supplementary material for "Expression changes confirm genomic variants predicted to result in allele-specific, alternative mRNA splicing": Supp. Methods and Results

A secondary set of common SNPs (separate from the initial SNP set; limited to 24 SNPs) were evaluated for their potential influence on mRNA splicing. First, all SNVs present in ICGC (International Cancer Genome Consortium) patients were evaluated by the Shannon Pipeline (SP) to find which SNPs alter splice site strength. Common SNPs (average heterozygosity > 10% in dbSNP 150) predicted to decrease natural splice site strength by the SP (where *ΔR_i_* < -1 bit) were further evaluated. A program was written to determine what ICGC patients had these flagged SNPs (either heterozygous or homozygous). The program then created a IGV (Integrative Genomics Viewer) bash script which instructed the software to: 1) load the patient’s RNAseq BAM file [from normal tissue, if available]; 2) set the viewing window to the region of interest [300nt window range; variant position centered]; 3) Use IGV commands ‘collapse’ and ‘sort’ to group similar RNAseq reads together [atypical spliced reads tend to move to the top of the viewing window], and; 4) take a screenshot. We then manually eliminated IGV images which did not meet our gene expression criteria (exon affected by the SNP must have ≥5 RNAseq reads present). As this generated thousands of images, we limited our analysis to two ICGC patients, chosen randomly aside from pre-selecting their tissue of origin (to increase the likelihood of finding expression in these regions): DO47132 [Renal Cell Cancer] and DO52711 [Chronic Lymphocytic Leukemia]. Images were evaluated sequentially (in order of rsID value) and only concluded once the first 24 SNPs meeting these criteria were found. Note that this analysis will not determine if the alternative splicing event is significantly higher in these patients when compared to non-SNP carriers. It only determines that alternative splicing events are present in these regions. We therefore queried the ValidSpliceMut database for these SNPs, as abnormal splicing was only flagged in the database if it was found to be significant above the RNAseq a control set of normals (Shirley *et al.*, 2019).

**Results:**

Another set of 24 common SNPs were also evaluated for their potential impact on mRNA splicing by RNA-Seq analysis of ICGC patients. Those resulting in significantly decreased natural splice site strength (*ΔR_i_* < -1 bit) were analyzed for SNP-derived alternative splicing events. SNPs fulfilling these criteria expressed at sufficient levels over the region of interest were: rs6467, rs36135, rs154290, rs166062, rs171632, rs232790, rs246391, rs324137, rs324726, rs448580, rs469074, rs518928, rs624105, rs653667, rs694180, rs722442, rs748767, rs751128, rs751552, rs752262, rs832567, rs909958, rs933208, and rs1018342 (Supplemental Table X). Splicing was predicted to be leaky at all of these SNPs (Rogan et al. 1998; final R_i_ ≥ 1.6 bits), ranging from 1.1 to 3.3 bit decreases in *R_i_* value.

Alternate mRNA splicing was seen for 14 SNPs: rs6467, rs36135, rs166062, rs171632, rs448580, rs469074, rs518928, rs694180, rs722442, rs752262, rs832567, rs909958, rs933208, rs1018342; Supplemental Table X). Reads spanning these regions revealed intron retention (N=12), activation of cryptic splicing (N=4), and complete exon skipping (N=3). 11 of these SNPs (79%) exhibited splicing patterns that significantly differed from the control alleles, and were represented in the ValidSpliceMut database. Interestingly, ValidSpliceMut contained entries for 7 of 10 SNPs where alternative splicing had not been found in the two patients reported in Supplemental Table X (rs246391, rs324137, rs624105, rs653667, rs748767, rs751128, rs751552). The observed significant splicing differences relative to controls for these SNPs occurred in distinct tumor types, consistent with tissue-specific effects of these SNP on splicing.

**Supplementary Table X: Secondary Set of Common SNPs Predicted to Weaken Natural Splice Sites**

| **Gene** | **rsID** | **HGVS Notation**  **(HG19)** | ***R_i_ initial*** | ***R_i_ final*** | *Δ****R_i_*** | **Alternative Splicing Observed** |
| --- | --- | --- | --- | --- | --- | --- |
| *CYP21A2* | rs6467 | [6:32006858C>A](https://validsplicemut.cytognomix.com/view.php?targets=447403+447404+819744&referenceName=6&alternateBases=A&start=32006858&referenceBases=C&assemblyId=GRCh37&includeDatasetResponses=HIT) | 6.1 | 4.5 | -1.6 | Intron Retention; Cryptic Site Use |
| *TRIM23* | rs36135 | 5:64890479A>C | 10.0 | 7.5 | -2.5 | Intron Retention |
| *ZFYVE16* | rs166062 | 5:79773028T>G | 14.1 | 11.8 | -2.4 | Intron Retention |
| *APBB3* | rs171632 | [5:139941318A>G](https://validsplicemut.cytognomix.com/view.php?targets=447388&referenceName=5&alternateBases=G&start=139941318&referenceBases=A&assemblyId=GRCh37&includeDatasetResponses=HIT) | 2.5 | 1.4 | -1.1 | Intron Retention; Use of Alternate Acceptor |
| *SMIM8* | rs448580 | [6:88040399T>G](https://validsplicemut.cytognomix.com/view.php?targets=325307+684696+684697&referenceName=6&alternateBases=G&start=88040399&referenceBases=T&assemblyId=GRCh37&includeDatasetResponses=HIT) | 6.9 | 4.4 | -2.5 | Exon Skipping |
| *FCHSD1* | rs469074 | [5:141024136T>G](https://validsplicemut.cytognomix.com/view.php?targets=428442+428443+428444+428445+428446+428447+428448+812575+812576+812577+812578+812579+812580+812581+812582+812583&referenceName=5&alternateBases=G&start=141024136&referenceBases=T&assemblyId=GRCh37&includeDatasetResponses=HIT) | 10.4 | 7.1 | -3.3 | Intron Retention |
| *KIFAP3* | rs518928 | [1:169890933A>G](https://validsplicemut.cytognomix.com/view.php?targets=447481&referenceName=1&alternateBases=G&start=169890933&referenceBases=A&assemblyId=GRCh37&includeDatasetResponses=HIT) | 15.6 | 14.5 | -1.1 | Intron Retention |
| *CEPT1* | rs694180 | [1:111726213A>G](https://validsplicemut.cytognomix.com/view.php?targets=447444) | 9.6 | 7.0 | -2.6 | Intron Retention |
| *ADCY10P1* | rs722442 | [6:41089681A>G](https://validsplicemut.cytognomix.com/view.php?targets=411458+814669+863502&referenceName=6&alternateBases=G&start=41089681&referenceBases=A&assemblyId=GRCh37&includeDatasetResponses=HIT) | 7.5 | 4.9 | -2.5 | Exon Skipping |
| *MICAL1* | rs752262 | [6:109770999G>C](https://validsplicemut.cytognomix.com/view.php?targets=427495+427496+427497+427498+427499+427500+427501+427502+427503+427504+427505+447412+792832&referenceName=6&alternateBases=C&start=109770999&referenceBases=G&assemblyId=GRCh37&includeDatasetResponses=HIT) | 4.0 | 2.7 | -1.4 | Intron Retention |
| *MAP3K1* | rs832567 | [5:56152416C>A](https://validsplicemut.cytognomix.com/view.php?targets=82130+432785+432786+432787+432788+432789&referenceName=5&alternateBases=A&start=56152416&referenceBases=C&assemblyId=GRCh37&includeDatasetResponses=HIT) | 6.8 | 5.0 | -1.7 | Exon Skipping; Intron Retention |
| *METTL13* | rs909958 | [1:171763522C>A](https://validsplicemut.cytognomix.com/view.php?targets=80782+83375+447482+447483&referenceName=1&alternateBases=A&start=171763522&referenceBases=C&assemblyId=GRCh37&includeDatasetResponses=HIT) | 11.4 | 9.9 | -1.4 | Cryptic Site Use |
| *DDX39B* | rs933208 | 6:31506648G>T | 4.8 | 3.7 | -1.1 | Intron Retention; |
| *CCT7* | rs1018342 | [2:73471653T>G](https://validsplicemut.cytognomix.com/view.php?targets=447970+447971&referenceName=2&alternateBases=G&start=73471653&referenceBases=T&assemblyId=GRCh37&includeDatasetResponses=HIT) | 5.9 | 4.4 | -1.5 | Intron Retention; Cryptic Exon use |
| *PPIP5K2* | rs154290 | 5:102537200T>G | 12.5 | 11.3 | -1.3 | Wildtype Only |
| *MYSM1* | rs232790 | 1:59131311G>T | 10.4 | 8.7 | -1.7 | Wildtype Only |
| *PDGFRB* | rs246391 | [5:149497177T>C](https://validsplicemut.cytognomix.com/view.php?targets=78727+921594+921595+921596+921597+921599+921600&referenceName=5&alternateBases=C&start=149497177&referenceBases=T&assemblyId=GRCh37&includeDatasetResponses=HIT) | 6.2 | 3.6 | -2.6 | Wildtype Only |
| *AARS2* | rs324137 | [6:44273546A>C](https://validsplicemut.cytognomix.com/view.php?targets=447409) | 10.2 | 8.9 | -1.3 | Wildtype Only |
| *USO1* | rs324726 | 4:76722353G>A | 11.8 | 8.8 | -3.0 | Wildtype Only |
| *KIF13A* | rs624105 | [6:17855864G>C](https://validsplicemut.cytognomix.com/view.php?targets=429720&referenceName=6&alternateBases=C&start=17855864&referenceBases=G&assemblyId=GRCh37&includeDatasetResponses=HIT) | 14.1 | 13.0 | -1.1 | Wildtype Only |
| *TNFRSF1B* | rs653667 | [1:12251808T>G](https://validsplicemut.cytognomix.com/view.php?targets=921006+921007+921008+921009+921010&referenceName=1&alternateBases=G&start=12251808&referenceBases=T&assemblyId=GRCh37&includeDatasetResponses=HIT) | 3.7 | 2.4 | -1.3 | Wildtype Only |
| *CCDC93* | rs748767 | [2:118731573G>A](https://validsplicemut.cytognomix.com/view.php?targets=78766+447505+447506&referenceName=2&alternateBases=A&start=118731573&referenceBases=G&assemblyId=GRCh37&includeDatasetResponses=HIT) | 4.6 | 3.4 | -1.1 | Wildtype Only |
| *CAPN2* | rs751128 | [1:223951841T>C](https://validsplicemut.cytognomix.com/view.php?targets=80842+80843+80844+447491&referenceName=1&alternateBases=C&start=223951841&referenceBases=T&assemblyId=GRCh37&includeDatasetResponses=HIT) | 5.3 | 4.2 | -1.1 | Wildtype Only |
| *FYCO1* | rs751552 | [3:46016851A>T](https://validsplicemut.cytognomix.com/view.php?targets=78853+447520+447521&referenceName=3&alternateBases=T&start=46016851&referenceBases=A&assemblyId=GRCh37&includeDatasetResponses=HIT) | 6.9 | 4.7 | -2.2 | Wildtype Only |
| Thick bars separate SNP-affected exons with and without RNAseq-observed alternate splicing events; If present, variant coordinates are hyperlinked to the ValidSpliceMut database. | | | | | | |
